# Ribosomal A-site interactions with near-cognate tRNAs drive stop codon readthrough

**DOI:** 10.1101/2023.06.19.543857

**Authors:** Zuzana Čapková Pavlíková, Petra Miletínová, Adriana Roithová, Klára Pospíšilová, Kristína Záhonová, Ambar Kachale, Thomas Becker, Ignacio M. Durante, Julius Lukeš, Zdeněk Paris, Petra Beznosková, Leoš Shivaya Valášek

## Abstract

tRNAs serve as a dictionary for the ribosome translating the genetic message from mRNA into a polypeptide chain. Besides this canonical role, tRNAs are involved in other processes like programmed stop codon readthrough (SC-RT). There, tRNAs with near-cognate anticodons to stop codons must outcompete release factors and incorporate into the ribosomal decoding center to prevent termination and allow translation to continue. However, not all near-cognate tRNAs promote efficient SC-RT. Here, we demonstrate that those that do, establish critical contacts between their anticodon stem (AS) and ribosomal proteins Rps30/eS30 and Rps25/eS25 forming the decoding site. Unexpectedly, the length and well-defined nature of the AS determines the strength of these contacts, which is reflected in organisms with reassigned stop codons. These findings open a new direction in tRNA biology that should facilitate the design of artificial tRNAs with specifically altered decoding abilities.

---

Occurrence of a stop codon in the decoding site (A site) of the elongating ribosome triggers uneven competition between the termination ternary complex, composed of eukaryotic release factors eRF1 and eRF3 and guanosine triphosphate (GTP), and near-cognate tRNAs recognizing two of the three bases of any of the three stop codons (reviewed in^1^). Higher energetic stability of the interaction between the UAG or UAA or UGA stop codon and the eRF1-eRF3-GTP complex in comparison to a corresponding near-cognate tRNA favors the former competitor in 99.9% of cases^2^. Stop codon recognition by eRF1 then stimulates GTP hydrolysis on eRF3, upon which eRF1 with the assistance of the ATPase ABCE1 (ATP binding cassette subfamily E member 1) mediates the release of the nascent peptide^3–5^. The post-termination complex is then disassembled, enabling its constituents to participate in further rounds of translation.

In the remaining 0.1% of cases, near-cognate tRNAs usurp the stop codon to avoid termination and instead allow the so-called stop codon readthrough (SC-RT) (reviewed in ^6,7^). As a result, C-terminally extended proteins are produced either as an infrequent failure of the system or, if programmed by various features acting *in trans* and/or *in cis*, as a desirable variant of a given protein with newly acquired functional qualities and/or cellular localization. Among the latter features belong the identity of the stop codon and surrounding sequence contexts^2,8–11^, proximal RNA structures^12^, RNA modifications^13,14^, presence of RNA binding proteins^15^ and eukaryotic initiation factor eIF3^16,17^, and the availability and quality of aminoacylated (aa) near-cognate tRNAs^18–21^.

Interestingly, not all near-cognate tRNAs are capable of promoting efficient SC-RT^18,21^. Those that can are called readthrough inducing (rti) tRNAs and – at least in *Saccharomyces cerevisiae* – include tRNA^Trp^_CCA_ and tRNA^Cys^_GCA_ for UGA with the 3^rd^ base wobble, the M iso-acceptor of tRNA^Gln^ for UAG with the 1^st^ base wobble, and tRNA^Tyr^_GUA_ for UAR (UAA and UAG collectively) stop codons with the 3^rd^ base wobble^19,20^. Virtually nothing is known about the qualities that significantly increase the odds of rti-tRNAs competing with eRF1 for the stop codon, nor about the underlying molecular mechanism of their preferred selection by the decoding center.

One hypothesis to understand how the rti-tRNAs get accepted by the ribosome that is poised to terminate can be found in the analogy with the 3^rd^ base wobbling during elongation. Cognate codon-anticodon recognition in the decoding center of the ribosome is read out by the two conserved decoding bases (A1492 and A1493 in bacteria), which are flipped out of helix (h) 44 of 18S rRNA to check a proper A-helix minor grove geometry of the mRNA:tRNA decoding complex. A cognate complex then induces a conformational change in the ribosomal small subunit, referred to as domain closure^22,23^. This movement was proposed to induce a tighter fit around the codon-anticodon minihelix and to help activate the GTP hydrolysis on eukaryotic elongation factor 1A (eEF1A). Domain closure appears to be a specific response to aa-tRNA selection and does not occur in the presence of eRF1^24^. Since the h44 decoding bases sense indirectly only the first two codon pairs, enabling the +3 position to participate in the wobble interactions^25^, the same architecture is believed to be induced with near-cognate tRNAs^26^. In other words, in order to be selected, near-cognate tRNAs have to go through the same activated state as fully cognate tRNAs. However, in physiological conditions, the probability of reaching the activated state is much more favored for the cognate interactions than for the non-cognate ones^24^, which is reflected in increased ribosome occupancy at the wobble codons^27,28^. The same is most likely true for tRNAs near-cognate to the stop codons with the 3^rd^ base wobble regardless of their ability to boost SC-RT or not.

The situation might be even more complicated with near-cognate tRNAs with the 1^st^ base wobble, one of which – the M iso-acceptor tRNA^Gln^_CUG_ (tRNA^Gln^_CUG_[M]) – is a very potent inducer of SC-RT compared to others from the same family, such as tRNA^Arg^, tRNA^Glu^, tRNA^Gly^, and tRNA^Lys18,19,21^. Given these differences, it is clear that in addition to the minor groove geometry check carried out by specific 18S rRNA bases, other factors influence the tRNA selection process, providing rti-tRNAs with the advantage of being selected at a significantly higher frequency than other near-cognate tRNAs. For example, in addition to 18S rRNA, four small ribosomal proteins (RPSes), namely Rps23/uS12, Rps15/uS19, Rps25/eS25 and Rps30/eS30, are part of the decoding site, with all of them having profound roles during codon sampling^24,29–31^. To name a few, it was proposed that the contacts with the codon-anticodon minihelix made by Rps15/uS19, and especially by Rps30/eS30 could increase the stability of aa-tRNAs during initial selection and accommodation^24,29^. Therefore, the question arises whether RPSes could play a discriminatory role.

Another factor that may influence SC-RT could lie directly in the nature of the tRNA backbone. Indeed, we have recently shown that tRNA^Trp^_CCA_ with a short anticodon stem (AS; 4-base pairs [bp] instead of the canonical 5-bp), which occurs naturally in several protists with all three stop codons reassigned to sense codons, markedly boosts SC-RT on UGA^32^. Curiously, this special quality of the 4-bp tRNA^Trp^_CCA_ originating outside of the anticodon loop represents one of the critical life-sustaining measures in these organisms with an altered genetic code.

Here, with help of the M iso-acceptor of tRNA^Gln^_CUG_, and tRNA^Trp^_CCA_ and tRNA^Cys^_GCA_ we show that not only the length but also the nature of the AS (in particular the base composition of its 4^th^ 28:42 base pair) critically contribute to the increased propensity of rti-tRNAs to SC-RT on UAR and UGA, respectively, at least in *S. cerevisiae* and *Trypanosoma brucei*. Bioinformatics analyses revealed that ciliates with UAR-to-glutamine reassignment have genomic bias towards this particular base pair composition with pyrimidine at position 28 and purine at position 42 of their tRNA^Gln^_UUG_. Using yeast genetics we have identified contacts that the AS of these rti-tRNAs and of rti-tRNA^Tyr^_GUA_ establish with specific amino acid residues of Rps30/eS30 and Rps25/eS25, respectively. Both of these proteins form the ribosomal decoding center. Disruption of these contacts greatly affects SC-RT-promoting potential that these rti-tRNAs normally allow. Therefore, we propose that, in addition to the degree of complementarity of the codon-anticodon minihelix and its interactions with the specific 18S rRNA decoding bases, tRNAs establish additional contacts with the decoding center components through their backbone, in particular through their AS. These so far unrecognized interactions could further propagate a tighter fit around the codon-anticodon minihelix, thereby increasing the likelihood that both cognate and specific near-cognate aa-tRNAs reach an activated state to be selected.

## RESULTS

### Primary sequence of tRNA^Gln^ determines the efficiency of stop codon readthrough in *S. cerevisiae* and *T. brucei*

We previously observed that whereas the tRNA^Gln^_CUG_ M iso-acceptor (tRNA^Gln^_CUG_[M]) dramatically induced SC-RT at all four UAG-N tetranucleotides, the other tRNA^Gln^_UUG_ iso-acceptors, such as tRNA^Gln^_UUG_[B] and tRNA^Gln^_UUG_[L], promoted SC-RT at their corresponding near-cognate UAA-N tetranucleotides only marginally^19^. Remarkably, the nuclear genome of the budding yeast *S. cerevisiae* contains altogether 10 genes encoding tRNA^Gln^. Whereas 9 of them encode different iso-decoders of the UUG anticodon iso-acceptor, there is only a single gene encoding the CUG anticodon iso-acceptor tRNA^Gln^_CUG_[M]. Apart from the distinct anticodons of the CUG M iso-acceptor *versus* the remaining 9 UUG iso-decoders, there are only minor variations in their backbone sequence embodied by three specific bases. These are A *versus* G at position 42 of the AS of the M iso-acceptor (also present in the tRNA^Gln^_UUG_[E2] iso-decoder), A *versus* G at position 51 in the pseudouridine (T) loop, and G *versus* A at position 66 in the acceptor stem (Fig. 1A; Extended Data Fig. 1). Notably, none of these bases is subject to otherwise widespread tRNA modifications (https://www.tmodbase.com)^33,34^. These differences prompted us to investigate the mechanistic basis for the elevated SC-RT-inducing potential of the tRNA^Gln^_CUG_[M] iso-acceptor. We wondered whether it is due to the differing anticodon and its modified nucleotide(s) or due to nucleotide variations in the primary sequence of the tRNA^Gln^ backbone.

**Figure 1.**
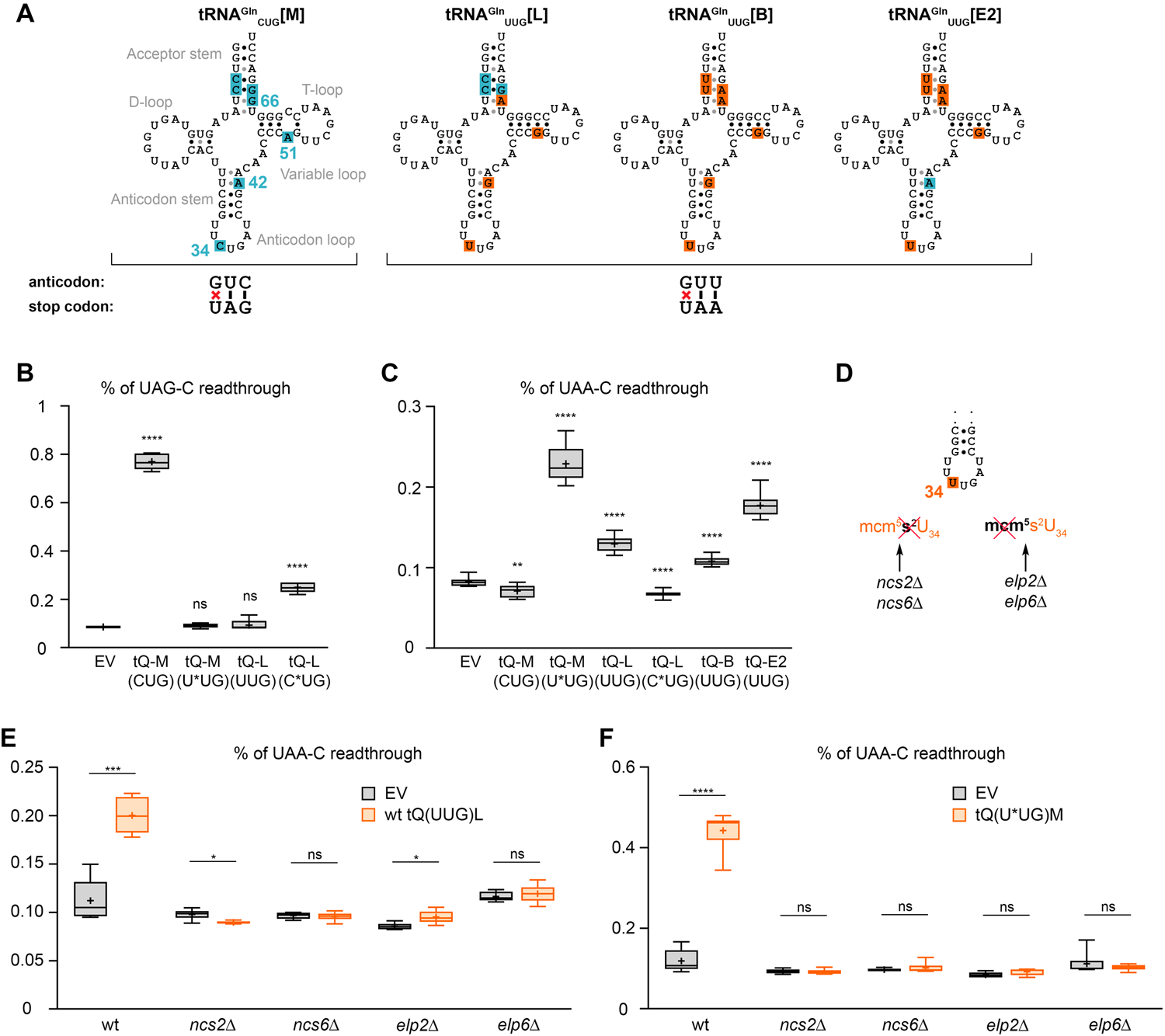
The efficiency of SC-RT of tRNA^Gln^ is influenced neither by the nature of anticodons, nor by their modifications. (A) Predicted secondary structures of all tRNA^Gln^ studied therein (retrieved from the GtRNAdb^69^). The bases differing between the CUG and UUG iso-acceptors are shown in blue and orange, respectively. (B - C) The nature of anticodons of tRNA^Gln^ does not impact the efficiency of SC-RT. The wt and indicated (*) mutant of CUG and UUG tRNA^Gln^ iso-acceptors with swapped anticodons were expressed individually (along with an empty vector - EV) in wt yeast cells and the efficiency of SC-RT on (B) UAG-C or (C) UAA-C was measured using the dual-luciferase system described in Methods. Readthrough values were normalized to the control cells (containing dual-luciferase cassette with CAA in place of a stop codon) and plotted as a percentage of readthrough. Each box in the box-and-whisker plot is represented by (B) n = 5 or (C) n ≥ 8 readthrough values. The mean value is marked (+); whiskers range from minimal to maximal values. Statistical significance was determined by the unpaired, two-tailed Welch’s *t* test; **** indicates p < 0.0001; *** p < 0.001; ** p < 0.01; * p < 0.05; ns = non-significant. (D) Scheme indicating the nature of the tRNA^Gln^_UUG_ anticodon modifications, as well as the corresponding deletion mutants of modifying enzymes tested in the study. (E - F) The mcm^5^s^2^ modification does not make a difference in the varying SC-RT potential between the M and other forms of tRNA^Gln^. (E) The wt tRNA^Gln^_UUG_[L] and (F) mutant tRNA^Gln^_U*UG_[M] iso-acceptors were expressed individually (along with EV) in wt and indicated mutant yeast cells and the efficiency of SC-RT on UAA-C was measured and evaluated as described in panels B – C; each box is represented by n ≥ 5 (E) and n = 6 values (F).

In order to test the impact of each anticodon, we took the M and L iso-acceptors (identical in the backbone sequence with the exception of the aforementioned bases) and simply swapped their anticodons creating the mutant tRNA^Gln^_U*UG_[M]^A42_A51_G66^ and tRNA^Gln^_C*UG_[L]^G42_G51_A66^ forms. Next, we expressed them individually (along with the wild-type [wt] tRNA^Gln^_UUG_[B] and tRNA^Gln^_UUG_[E2] iso-decoders) in yeast cells and measured the SC-RT efficiency using our well-established dual-luciferase system^19^, featuring Renilla and Firefly luciferases set in the same frame but separated by either the UAG or UAA stop codons. We confirmed markedly induced SC-RT of tRNA^Gln^_CUG_[M] at UAG-C (Fig. 1B). In addition, we observed that substituting its anticodon from CUG to UUG (tRNA^Gln^_U*UG_[M]^A42_A51_G66^) did not impair the ability of this anticodon-swapped mutant tRNA^Gln^ to promote efficient SC-RT also on UAA (Fig. 1C). (Please note that absolute values are not comparable between experiments shown in Figure 1B and C because the intrinsic ability of stop codons to allow natural readthrough differs among them, with UGA being the most readthrough-permissive stop codon, while UAA the least (UGA > UAG > UAA; reviewed in^35^). Therefore, we conclude that the impact of the stop codon identity *per se* is negligible and that the readthrough potential of tRNA^Gln^_CUG_[M] seems to lie predominantly in its primary sequence. In strong support, the reciprocal substitution of the anticodon from UUG to CUG in the tRNA^Gln^_C*UG_[L]^G42_G51_A66^ mutant resulted in comparable levels of SC-RT on UAG, as in the case of the wt tRNA^Gln^_UUG_[L] on UAA (Fig. 1B and C).

For consistency, all tRNA constructs created in this study have been flanked by the M iso-acceptor untranslated regions (“M” UTRs). Importantly, a direct comparison of all studied UUG iso-decoders flanked by either the genuine or “M” UTRs revealed no difference in their SC-RT-inducing potential (Extended Data Fig. 2). Northern blotting revealed that most of the tRNA^Gln^ variants employed in this study are well expressed, as shown previously^19^ and here (Extended Data Fig. 3A); only two tRNA^Gln^ variants did not reach elevated levels compared to empty vector (EV), as explained below. In any case, these control experiments rule out that the observed differences are due to differing levels of individual tRNA variants and/or their integrity issues.

Importantly, while no information exists about modifications of the CUG anticodon and/or its adjacent nucleotides in the M iso-acceptor^14^, the UUG anticodon of all tRNA^Gln^_UUG_ iso-decoders undergoes the 5-methoxycarbonylmethyl-2-thiouridine (mcm^5^s^2^) modification at U_34_ during the tRNA maturation process (Fig. 1D)^36^. Hence, we were wondering whether this modification could compromise the SC-RT potential of the tRNA^Gln^_UUG_ iso-decoders with the UUG anticodon sampling the UAA stop codon in the ribosomal A site. Therefore, we expressed the tRNA^Gln^_UUG_[L] iso-decoder in yeast strains individually deleted for genes required for the U_34_ modification; namely *ncs2Δ, ncs6Δ, elp2Δ*, and *elp6Δ* (kindly provided by Sebastian Leidel) (Fig. 1D). We assumed that if mcm^5^s^2^ at U_34_ negatively influences SC-RT, we should record higher SC-RT values. On the contrary, for all four tested strains the SC-RT efficiency of the wt tRNA^Gln^_UUG_[L] dropped to the “EV” background levels (Fig. 1E). Essentially the same effect was observed when the mutant tRNA^Gln^_U*UG_[M]^A42_A51_G66^ iso-acceptor was expressed in these deletion strains (Fig. 1F). Northern blotting revealed that neither of these deletions markedly impacted the expression levels of these two tRNAs (Extended Data Fig. 3B). Please note that the Northern probe was designed to fully match bases 43 through 72 of the tRNA^Gln^_CUG_[M] molecule. Therefore, the recognition specificity of tRNA^Gln^_UUG_[L] is reduced due to two internal mismatches at positions A66 and G51, which in turn magnifies the relative noise signal in the EV lanes observed in the experiment with tRNA^Gln^_UUG_[L] (upper three lanes), since this probe naturally recognizes all other native tRNA^Gln^_UUG_ isodecoders.

Collectively, these results thus not only correlate with the previous work^14^, but also clearly suggest that it is not the modification of mcm^5^s^2^ that causes the differential SC-RT potential between the M and all other forms of tRNA^Gln^. On the contrary, it modestly promotes the mis-incorporation of the tRNA^Gln^_UUG_ iso-decoders at the UAA stop codon. Therefore, we ruled out the impact of the anticodon and its modifications. To test the effect of the tRNA^Gln^ primary sequence, we individually or in various combinations substituted the bases of wt tRNA^Gln^_CUG_[M]^A42_A51_G66^ at positions 42, 51 and 66 with those occurring in the L iso-acceptor. While A51G increased SC-RT, G66A significantly decreased it and, strikingly, A42G displayed exactly the same effect as the triple mutation (in tRNA^Gln^_CUG_[M]^A42G_A51G_G66A^), i.e., the most pronounced decrease (Fig. 2A). Note that the triple mutant is – apart from the anticodon – fully identical to the tRNA^Gln^_UUG_[L]^G42_G51_A66^ iso-acceptor, resembling all other tRNA^Gln^_UUG_ iso-decoders (Fig. 1A). Consequently, tRNA^Gln^_CUG_[M]^A42G_A51G_G66A^ generates weaker Northern blot signal due to two internal mismatches with the Northern probe used as described above (Extended Data Fig. 3A). Since the effect of the A42G_G66A double substitution practically mimicked that of A42G alone (Fig. 2A), we conclude that the A42 base is the major readthrough effector of the tRNA^Gln^_CUG_[M] iso-acceptor. In agreement, the tRNA^Gln^_UUG_[E2] iso-decoder that naturally carries adenine at position 42 – in contrast to all other tRNA^Gln^_UUG_ iso-decoders – showed the most efficient SC-RT among them (Fig. 1C).

**Figure 2.**
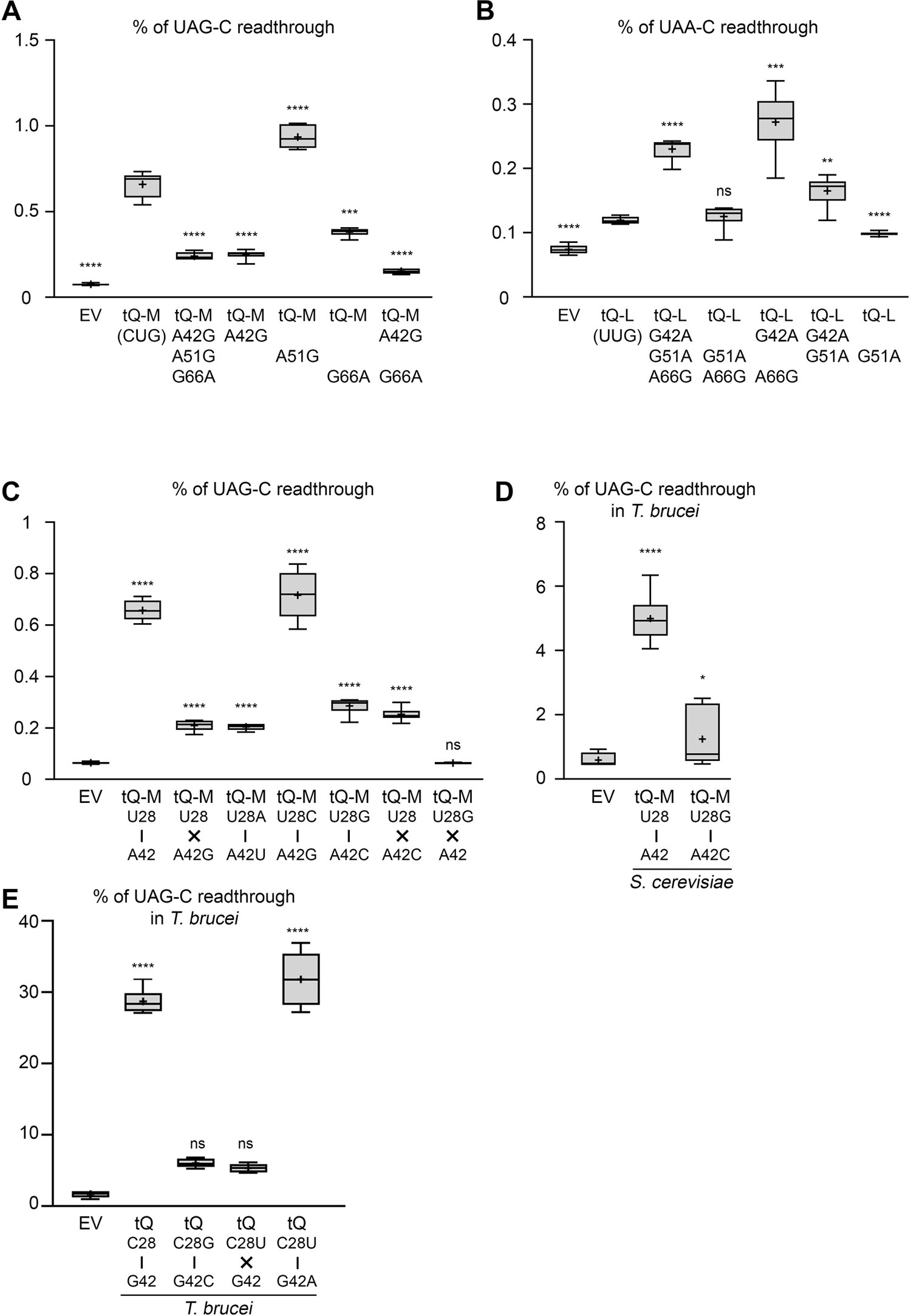
Primary sequence of tRNA^Gln^_CUG_ determines the efficiency of stop codon readthrough in *S. cerevisiae* and *T. brucei*. (A - B) The wt and mutant (A) CUG and (B) UUG tRNA^Gln^ iso-acceptors with indicated backbone substitutions were expressed individually (along with EV) in wt yeast cells and the efficiency of SC-RT on (A) UAG-C or (B) UAA-C was measured and evaluated as described in Fig. 1B – C; each box is represented by n ≥ 5 values. (C) The nature of the 28:42 base pair of the AS is critical for the increased propensity of tRNA^Gln^_CUG_[M] iso-acceptor for SC-RT. The wt and indicated mutant variants of the tRNA^Gln^_CUG_[M] iso-acceptor with substitutions of their 28:42 base pair were expressed individually (along with EV) in wt yeast cells and the efficiency of SC-RT on UAG-C was measured and evaluated as described in Fig. 1B – C; each box is represented by n = 6 values. The nature of hydrogen bonding between bases 28 and 42 of the AS is indicated by a vertical black line (Watson-Crick [W-C] base paring) or a black cross (no W-C paring). (D) The wt and the A42G mutant of the tRNA^Gln^_CUG_[M] iso-acceptor were expressed individually (along with EV) in the procyclic forms of *T. brucei* bearing dual-luciferase cassette with in-frame UAG-C. Transformed cells were processed for UAG readthrough measurements as described in Methods. Readthrough values were normalized to those of the control cell line (containing dual-luciferase cassette without in-frame UAG) and evaluated as described in Fig. 1B – C; each box is represented by n = 9 values (3 individual experiments each including 3 biological replicates). (E) The wt and the indicated mutant variants of the *T. brucei* tRNA^Gln^_CUG_ were expressed individually (along with EV) in the procyclic forms of *T. brucei* bearing dual-luciferase cassette with in-frame UAG-C. Transformed cells were processed for UAG readthrough measurements as described in Methods. Readthrough values were normalized to those of the control cell line (containing dual-luciferase cassette without in-frame UAG) and evaluated as described in Fig. 1B – C; each box is represented by n = 9 values (3 individual experiments each including 3 biological replicates).

Next, we reversed our approach and instead of diminishing/eliminating the efficiency of SC-RT of the wt tRNA^Gln^_CUG_[M]^A42_A51_G66^iso-acceptor by targeted substitutions (Fig. 2A), we took the wt tRNA^Gln^_UUG_[L]^G42_G51_A66^ iso-decoder with its natural UUG anticodon and attempted to boost its propensity for SC-RT with reciprocal substitutions at positions 42, 51 and 66. As can be seen in Fig. 2B, we obtained data inversely reflecting those shown in Fig. 2A. Altogether, these findings clearly illustrate that there are features in the backbone of the tRNA^Gln^_CUG_[M] iso-acceptor – with the prevalent participation of A42 in its AS – that renders this particular tRNA highly efficient in promoting SC-RT.

### Nature of the 28:42 base pair of the anticodon stem is critical for increased propensity of tRNA^Gln^_CUG_[M] iso-acceptor for SC-RT

Adenine 42 of the tRNA^Gln^_CUG_[M] iso-acceptor base pairs with U28 in its AS (Fig. 1A). To investigate to what extent the Watson-Crick (W-C) base pairing between the 28:42 pair contributes to the efficiency of readthrough, we substituted bases at these positions with those either breaking base pairing or restoring it by different means. To the best of our knowledge, none of these bases of this particular tRNA is subject to modifications^34,37^. Of note, the position 28 is known to be modified by pseudouridine synthase Pus1 in several tRNAs but not in tRNA^Gln^ (^37^). In agreement, deletion of *PUS1* had no effect on the efficiency of SC-RT of tRNA^Gln^_CUG_[M] in our previous experiments^19^. First, we tested the variant retaining the W-C base pairing but with the U28:A42 swapped to create the U28A:A42U pair. Interestingly, its UAG-C SC-RT activity dropped by ∼3-fold, i.e. to the same level as when the base pairing was broken by the key A42G substitution (Fig. 2C). To the contrary, restoring the base-paring of A42G by introducing the U28C substitution behaved as wt tRNA^Gln^_CUG_[M]. Swapping the U28C:A42G pair, i.e., creating U28G:A42C, however, also led to ∼3-fold drop in the SC-RT efficiency, similarly as in case of the U28A:A42U swap (Fig. 2C). As expected, the pyrimidine:pyrimidine (U28:A42C) pair did not show elevated SC-RT levels (Fig. 2C). The purine:purine (U28G:A42 and U28A:A42) pairs displayed no measurable activity above EV (Fig. 2C and data not shown for U28A:A42), indicating that these substitutions produce non-functional tRNAs. Consistently, Northern blotting confirmed increased expression (compared to EV) of all tRNAs but the latter tRNA^Gln^_CUG_[M]^U28G^ (Extended Data Fig. 3A). (Lower signal of tRNA^Gln^_CUG_[M]^A42G_A51G_G66A^ is explained above). These results thus clearly demonstrate that besides purine:purine variants (tRNA^Gln^_CUG_[M]^U28G^ and tRNA^Gln^_CUG_[M]^U28A^), overall integrity, charging and functionality of all other mutants is not affected, because their activity never falls below the ∼3-fold stimulation level of tRNA^Gln^_C*UG_[L] over EV displayed in Figure 1B. Taken together, our findings indicate that in addition to the W-C base pairing of the 28:42 pair, pyrimidine is strongly preferred at position 28 and purine at position 42 for the tRNA^Gln^_CUG_[M] iso-acceptor to stimulate efficient SC-RT.

To further support the key role of this specific arrangement, we expressed the wt tRNA^Gln^_CUG_[M] iso-acceptor and its purine:pyrimidine U28G:A42C mutant variant in *T. brucei*, where we recently established the SC-RT dual lucifierase system and where conserved effects of mutations affecting the length of the AS of tRNA^Trp^_CCA_ were initially observed^32^. Whereas wt tRNA^Gln^_CUG_[M] exhibited very efficient SC-RT even in this parasitic protist evolutionary very distant from yeast, the U28G:A42C mutation completely eliminated it (Fig. 2D). Furthermore, *T. brucei* contains one gene encoding tRNA^Gln^_UUG_ and two genes for tRNA^Gln^_CUG_. All three observe the pyrimidine:purine rule with C28 and G42 and the latter iso-acceptor promotes efficient SC-RT in *T. brucei* (Fig. 2E). While swapping C with G (in C28G:G42C) or disrupting the W-C base-pairing by the C28U substitution completely eliminated the SC-RT potential of tRNA^Gln^_CUG_, substituting pyrimidine with pyrimidine and purine with purine (in C28U:G42A) preserved efficient readthrough (Fig. 2E). Northern blotting confirmed stable expression of all tRNAs tested (Extended Data Fig. 3C). Overall, these results strongly suggest a general nature of preferential selection of tRNA^Gln^ with pyrimidine 28 and purine 42 forming the 4^th^ AS pair by the ribosome.

### Ciliates with UAR-to-glutamine reassignment have genomic bias towards the 28:42 base pair

The ciliates (*Ciliophora*) are well-known for their propensity to stop codon reassignments^38^. In several ciliate lineages, UAR stop codons occur within their coding sequences, being reassigned to encode glutamine. Functionally, these organisms require tRNA^Gln^ to allow SC-RT on in-frame UAR codons for the synthesis of full-length proteins. Therefore, we asked whether there is a genomic bias in UAR-to-glutamine reassigned ciliates towards tRNAs^Gln^-encoding genes with the 4^th^ AS base pair observing the pyrimidine 28 : purine 42 rule as described above, which allows more efficient SC-RT. To investigate this, we aligned tRNAs^Gln^ from two groups of ciliates: those with the canonical genetic code (“stop”) and those with the UAR-to-glutamine (“Q”) reassignment, and further split these tRNAs according to their anticodon into two subgroups (“CTG” *versus* “TTG”) (Extended Data Excel File 1A). These four subgroups were further divided into those having the pyrimidine 28 : purine 42 base pair *versus* any other combination. We then counted fractions of the individual 28:42 base pair identities within all these subcategories and expressed them as a ratio of the pyrimidine 28 : purine 42 base pair identities *versus* any other combination (pyr:pur/other) for the four major subgroups (Extended Data Table 1; Extended Data Excel File 1B). These results revealed that the pyr:pur/other ratio is 1:3 for “Q” and 1:1 for “stop” in the CTG subgroups, whereas it is 2:1 for “Q” and 1:2 for “stop” in the TTG subgroups, showing expected bias for the latter subgroup.

To explain why the CTG group lacks this bias, we analyzed all available predicted proteomes of ciliates with reassigned TAR codons (out of total 44 analyzed genomic assemblies only 12 come with predicted proteomes) (Extended Data Excel File 1A). In these coding sequences, we determined the number of codons that encode glutamine, i.e., CAA, CAG, TAA and TAG, and tested the presumption that the distribution of all four codons encoding glutamine is the same (i.e., the expected numbers are [CAA+CAG+TAA+TAG]/4). The χ^2^ p-value was < 0.00001 for each organism, suggesting that the differences in codon abundancies are significant (Extended Data Excel File 1C). Notably, with the single exception of *Halteria grandinella*, TAA (with the TTG anticodon) is the most frequently used codon in this set, and in all cases its representation is at least 2-fold higher (up to 6.4-fold) than that of TAG (CTG anticodon) (Extended Data Fig. 4). Although the expression levels of tRNAs^Gln^ in these organisms are not known, our finding of the biased Gln codon representation in favor of TAA offers a logical explanation why the expected bias for the pyrimidine 28 : purine 42 base pair was not observed also in the tRNAs^Gln^_CUG_ group, which was therefore not selected for. On the contrary, it seems that tRNAs^Gln^_UUG_ have been under strong selection to prioritize the pyrimidine 28: purine 42 base pair identity in their AS to boost readthrough of the most abundant in-frame TAA codon.

### Interplay between tRNA^Gln^ primary structure and factors modulating SC-RT

Previously, we and others showed that the identity of a nucleotide following the termination codon (position/base +4) defines a so-called stop codon tetranucleotide, which determines the preferences of rti-tRNAs for a given tetranucleotide^11,19^. In this respect, as aforementioned, we demonstrated that tRNA^Gln^_CUG_[M] was able to induce SC-RT at all UAG-N tetranucleotides with the following preferences: -G > -A > -C > - U^19^. To examine whether the tRNA^Gln^_CUG_[M] primary backbone sequence has an impact on this UAG-N preference, we employed the dual-luciferase assay with selected, individually expressed substitution variants of tRNA^Gln^_CUG_[M]. All of them incorporated at the UAG-N tetranucleotides with the same trend as wt (Fig. 3A). Essentially the same result was observed with two naturally occurring readthrough-inducing hexanucleotide sequences downstream of the UAG stop codon, in particular the tobacco mosaic virus (TMV) UAG-CAATTA context (UAG-TMV) and the UAG-CAACTA context of the yeast *BSC4* gene (UAG-BSC4) (Fig. 3B). Therefore, as in case of the stop codon *per se*, there seems to be no functional interplay between the key tRNA backbone bases and the stop codon context including the key +4 base of the stop codon tetranucleotide.

**Figure 3.**
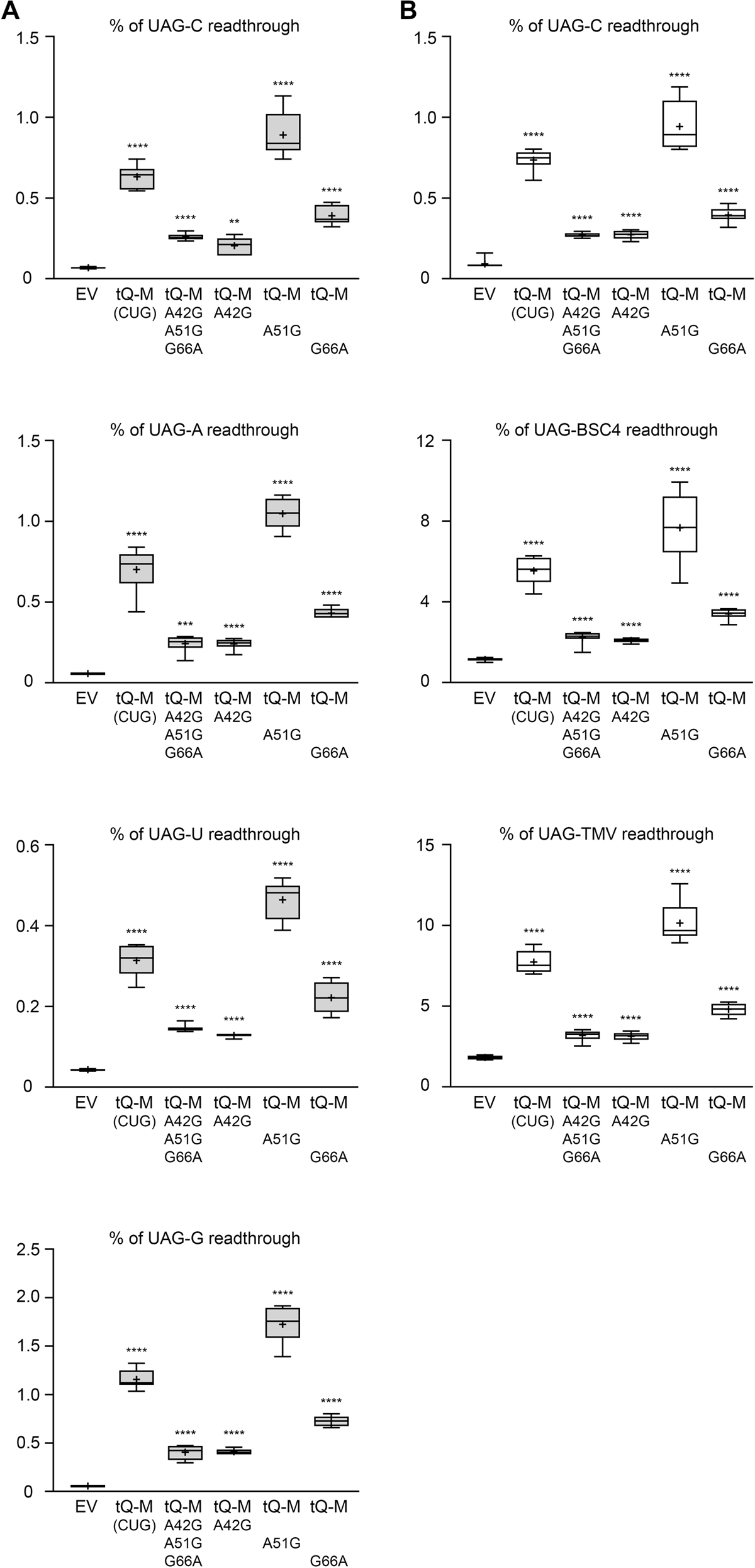
Stop codon context has no impact on the SC-RT efficiency of the tRNA^Gln^_CUG_[M] iso-acceptor. (A - B) The wt and mutant variants of the tRNA^Gln^_CUG_[M] iso-acceptor with indicated backbone substitutions were expressed individually (along with EV) in wt yeast cells and the efficiency of SC-RT on (A) UAG-N or (B) the UAG-C, -BSC4 and –TMV stop codon contexts was measured and evaluated as described in Fig. 1B – C; (A) each box is represented by n = 6 values (2 individual experiments each including 3 biological replicates); (B) each box is represented by n ≥ 7 values (2 individual experiments each including n ≥ 3 biological replicates).

Earlier we also revealed that in addition to its indispensable initiation roles^39^, the multisubunit translation initiation complex eIF3 also controls termination and critically promotes programmed SC-RT on all three stop codons. This eIF3 function is evolutionary conserved^16,17^. Detailed analysis of the UGA and UAA stop codons and all near-cognate tRNAs suggested that – at least in case of these two stop codons – eIF3 promotes incorporation of only those rti-tRNAs that have a mismatch at the third (wobble) position; i.e., rti-tRNA^Trp^_CCA_ and rti-tRNA^Cys^_GCA_ for UGA, and rti-tRNA^Tyr^_GUA_ for UAA. Here, we asked whether eIF3 also contributes to UAG SC-RT promoted by the tRNA^Gln^_CUG_[M] iso-acceptor, which has a mismatch at the first position. And if yes, whether there is some interplay between eIF3 and the tRNA^Gln^_CUG_[M] primary structure. Expressing wt tRNA^Gln^_CUG_[M] in the yeast strain with either wt eIF3 (*TIF35*) or carrying the *tif35-KLF* mutation, compromising its termination role^17^, revealed that incorporation of tRNA^Gln^_CUG_[M] at the UAG-C stop codon was markedly impaired in the *tif35-KLF* mutant (Fig. 4A; note different scales and compare “tQ_CUG_“ between both panels), as observed earlier with other two stops^11,17^. Next, we expressed mutant tRNA^Gln^_CUG_[M]^A42G_A51G_G66A^ in both strains seeking a genetic interaction with *tif35-KLF*. However, none was observed (Fig. 4A); the fold decrease between wt tRNA^Gln^_CUG_[M] expressed in wt *versus tif35-KLF* cells was similar to that of tRNA^Gln^_CUG_[M]^A42G_A51G_G66A^ (∼4-fold). These results imply that eIF3: 1) significantly promotes SC-RT of all four rti-tRNAs (tRNA^Cys^_GCA_, tRNA^Trp^_CCA_, tRNA^Tyr^_GUA_, tRNA^Gln^_CUG_[M]) regardless of whether the mismatch occurs at the first or third positions; and 2) does not facilitate the ability of the tRNA^Gln^_CUG_[M] iso-acceptor to markedly boost SC-RT. For mechanistic explanation of the eIF3 role in SC-RT please see^39^.

**Figure 4.**
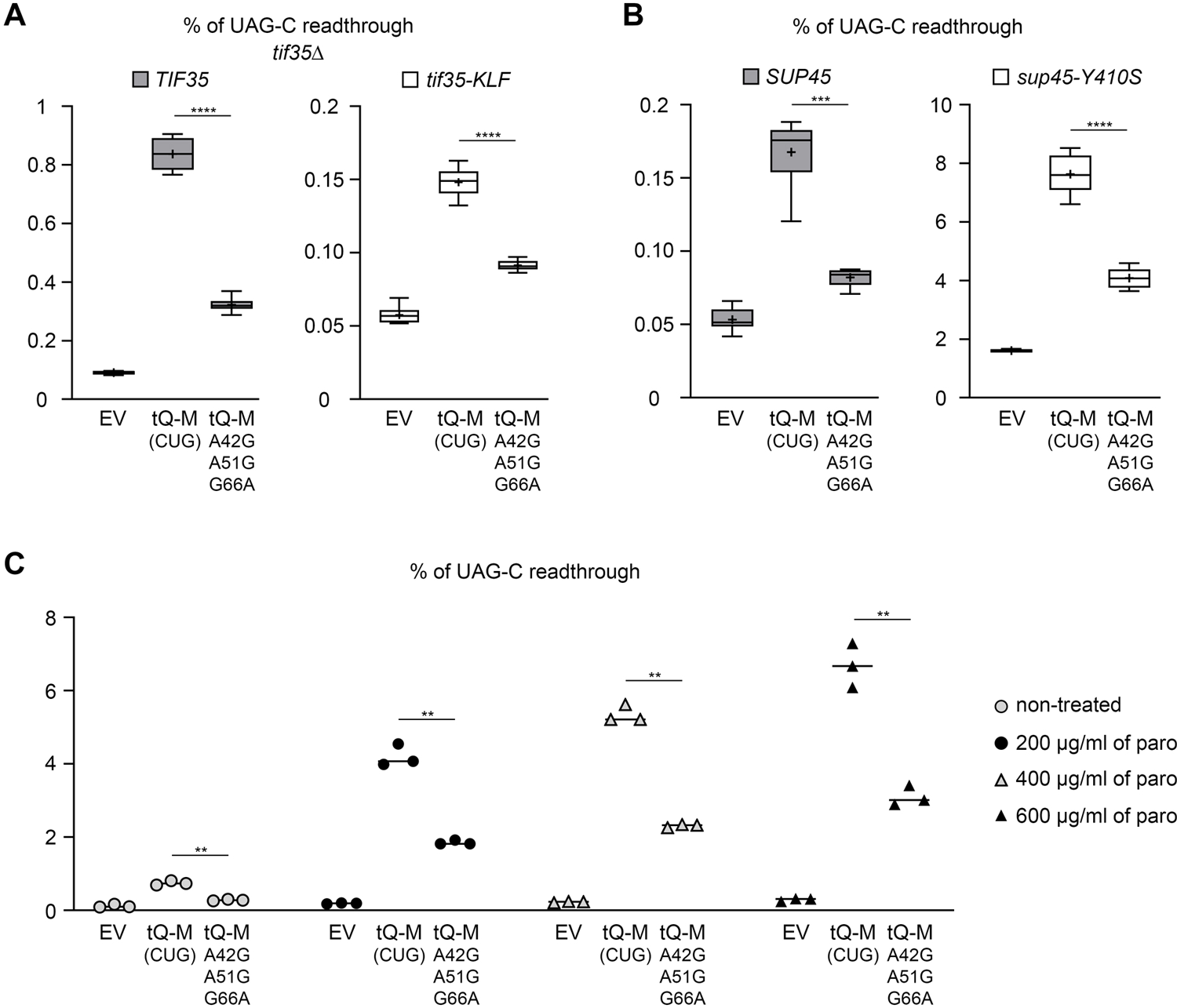
Interplay between the tRNA^Gln^_CUG_ primary structure and factors modulating SC-RT. (A) SC-RT-promoting translation initiation factor eIF3 does not influence the SC-RT efficiency of the tRNA^Gln^_CUG_[M] iso-acceptor. The wt and a mutant variant of the tRNA^Gln^_CUG_[M] iso-acceptor with indicated backbone substitutions were expressed individually (along with EV) in wt and *tif35-KLF* mutant yeast cells and the efficiency of SC-RT on UAG-C was measured and evaluated as described in Fig. 1B - C; each box is represented by n = 6 values (2 individual experiments each including 3 biological replicates). (B) Disrupting the eRF1-eRF3 interaction by a specific mutation of eRF1 does not influence the SC-RT efficiency of the tRNA^Gln^_CUG_[M] iso-acceptor. The wt and a mutant variant of the tRNA^Gln^_CUG_[M] iso-acceptor with indicated backbone substitutions were expressed individually (along with EV) in wt and *sup45-Y410S* mutant yeast cells and the efficiency of SC-RT on UAG-C was measured and evaluated as described in Fig. 1B - C; each box is represented by n = 6 values. (C) Deformation of the near-cognate codon-anticodon helix during the codon sampling step by paromomycin does not influence the SC-RT efficiency of the tRNA^Gln^_CUG_[M] iso-acceptor. The wt and a mutant variant of the tRNA^Gln^_CUG_[M] iso-acceptor with indicated backbone substitutions were expressed individually (along with EV) in wt yeast cells in the presence of an increasing concentration of paromomycin and the efficiency of SC-RT on UAG-C was measured and evaluated as described in Fig. 1B – C; each box is represented by n = 3 values.

Next, we examined the wt and mutant tRNA^Gln^_CUG_[M] in the background of the *sup45-Y410S* mutation of eRF1 (encoded by *SUP45*). This mutant directly disrupts the eRF1-eRF3 interaction, which in turn markedly weakens the eRF1 binding to the A-site-situated stop codon, resulting in robust SC-RT^40^. We wondered whether reducing the A-site-binding affinity of eRF1, as the tRNA’s major competitor for this site, would mitigate differences between the wt and mutant tRNA^Gln^_CUG_[M] (Fig. 4B). This was not the case, as both wt and mutant tRNA^Gln^_CUG_[M] (∼45-50-fold) increased SC-RT to similarly robust extent in the *sup45-Y410S* mutant *versus* wt cells. This result clearly demonstrates that the documented difference in the propensity for SC-RT between M and other forms of tRNA^Gln^ does not originate from varying ability of these tRNAs to outcompete eRF1 during stop codon recognition.

Paromomycin is a widely used drug for translation termination studies relaxing the A-site codon decoding pocket. In particular, its binding to the A-site results in deformation of the near-cognate codon-anticodon helix during the codon sampling step, after which the ribosome does not actively sense the correct W-C base pairing geometry and thus fails to discriminate against near-cognate tRNAs^41,42^. Since we previously showed that paromomycin robustly potentiates the tRNA^Gln^_CUG_[M] ability to boost SC-RT at UAG-N^19^, we examined whether it elicits varying effects on tRNA^Gln^_CUG_ variants differing in their backbone sequence. Cells expressing wt or mutant tRNA^Gln^_CUG_[M] in the presence of increasing concentration of paromomycin (200 µg/ml, 400 µg/ml, and 600 µg/ml) displayed expectedly much higher levels of SC-RT, nonetheless, the wt/mutant ratio in any of these concentrations remained practically unchanged (Fig. 4C). Please note that tested paromomycin concentration had no significant impact on cell viability (data not shown). This suggests that paromomycin-induced changes of the codon decoding pocket geometry, preventing the ribosome from discriminating against near-cognate tRNAs, do not affect the stimulatory effect of the tRNA^Gln^_CUG_ backbone bases on SC-RT. Where does this effect come from then?

### Ribosomal protein Rps30/eS30 promotes incorporation of the tRNA^Gln^_CUG_[M] iso-acceptor to the A site

Next, we wondered whether the tRNA backbone bases establish direct interactions with the A-site ribosomal components outside of the codon decoding pocket to facilitate the codon sampling and/or accommodation processes. Previous reports suggested that the aa-tRNA interacts with h18 and h30 of 18S rRNA and small ribosomal proteins Rps23/uS12, Rps15/uS19, Rps25/eS25, Rps30/eS30, and/or Rps31/eS31^24,29–31^. Based on the available data and RCSB PDB database searches we focused on the latter three RPSes because their flexible domains could stretch out to contact the AS base pair 28:42 (Fig. 5A; please note that the tRNA model shown in this figure [based on *E. coli* tRNA^Lys^] represents a mixture of tRNAs and is only used for illustration purposes; it does not represent tRNA^Gln^). In particular, the N-terminus of Rps30/eS30, the C-terminus of Rps15/uS19, and/or the N-terminus of Rps25/eS25^24,29,30^. Based on the available literature, the most intriguing contacts could be mediated by the first five extreme N-terminal residues and Arg10 of Rps30/eS30^24,29^ (Fig. 5B). To test the prospective role of Rps30/eS30 in facilitating the tRNA^Gln^_CUG_[M] accommodation in the A site, we generated several site-specific substitutions of the first ten N-terminal residues of Rps30/eS30 tagged with a FLAG tag at its C-terminus and expressed them in a yeast *rps30a rps30b* double deletion strain. None of the mutants displayed a severe growth phenotype at any tested temperatures (Fig. 5C and data not shown). Interestingly, whereas *rps30b-Δ2-6-FLAG, -5AAAAAA10-FLAG, -K3G-FLAG* and *-H5G-FLAG* mutants robustly decreased SC-RT on UAG-BSC4, *rps30b-A2G-FLAG* and *-V4A-FLAG* increased it, while *rps30b-R10A-FLAG* had a minimal effect (Fig. 5D). Similar results were obtained with UAA-BSC4 and UGA-BSC4 stop codons (Extended Data Fig. 5A – B). (Note, that the robustness of the increase in SC-RT induced by *rps30b-V4A-FLAG* exclusively on UAA and a significant decrease of *rps30b-R10A-FLAG* exclusively on UGA are discussed below.) Altogether, these results clearly implicate the N-terminus of Rps30/eS30 in controlling the stop codon decoding process.

**Figure 5.**
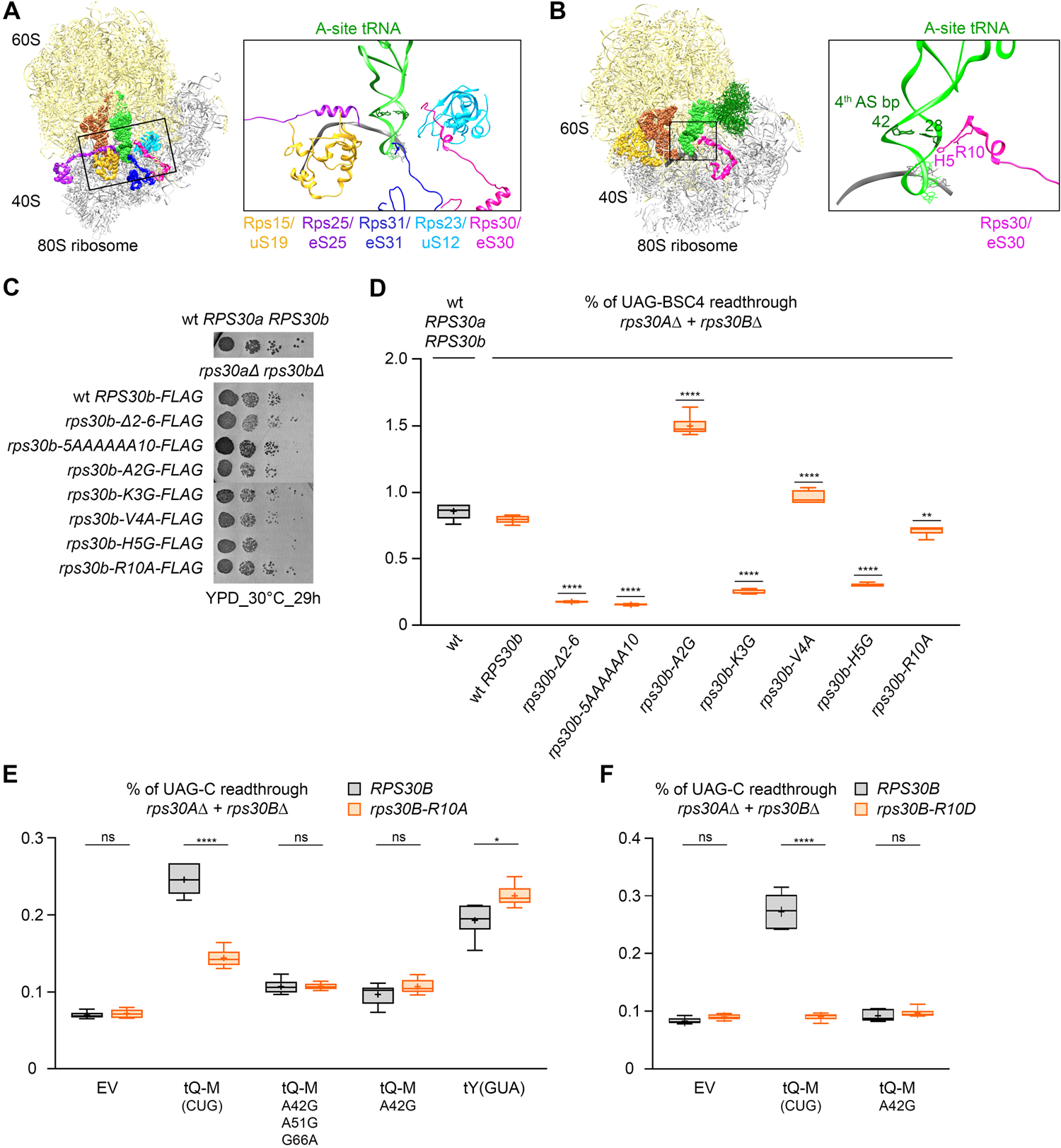
Ribosomal protein Rps30/eS30 promotes incorporation of the tRNA^Gln^_CUG_[M] iso-acceptor to the ribosomal A site. (A) Model of the yeast 80S ribosome (PDB-ID 7RR5; left) shown from the top and zoomed-in to highlight the A site (right). Shown are prospective contacts between the tRNA AS and ribosomal proteins in the A site. The 28:42 base-pair is depicted in dark green with spheres (left panel) or with sticks (right panel). Note that the terminal tails of Rps15/uS19, Rps25/eS25, Rps31/eS31, Rps23/uS12 and Rps30/eS30 are known to be flexible and the exact positions and the displayed length of fitted amino acids vary between different published structures. The last amino acids fitted here are as follows: Gly129 of the Rps15/uS19 C-terminus; Gln6 of the Rps25/eS25 N-terminus; Arg4 of the Rps31/eS31 N-terminus; full-length Rps23/uS12; and Ala2 of the Rps30/eS30 N-terminus. (B) Model of the rabbit 80S ribosome (PDB-ID 5LZS; left) zoomed-in to highlight the A site (right). Shown are prospective contacts between the 4^th^ base-pair of the tRNA AS and the ribosomal protein Rps30/eS30 in the A site. The A-site tRNA is shown in green (the 4^th^ base-pair of its AS in dark green) and Rps30/eS30 in magenta with His5 and Arg10 highlighted. (C) Growth rate analysis of the N-terminal substitution mutants of Rps30/eS30. The wt and indicated Rps30/eS30 mutant yeast cells were spotted in four serial 10-fold dilutions on YPD media and incubated for 29 h at 30 °C. (D) Rps30/eS30 mutants differentially modulate the SC-RT efficiency at the UAG stop codon. The wt and indicated Rps30/eS30 mutant yeast cells were tested for the efficiency of SC-RT in the UAG-BSC4 stop codon context as described in Fig. 1B – C; each box is represented by n ≥ 5 values. (E) R10 of Rps30/eS30 may potentiate the SC-RT-promoting ability of tRNA^Gln^_CUG_[M] by directly contacting the 28:42 base pair of its AS. The wt and indicated mutant variants of the tRNA^Gln^_CUG_[M] iso-acceptor, as well as control tRNA^Tyr^_GUA_, were expressed individually (along with EV) in wt and *rps30B-R10A* mutant yeast cells and the efficiency of SC-RT on UAG-C was measured and evaluated as described in Figure 1B – C; each box is represented by n ≥ 5 values. (F) Substituting R10 of Rps30/eS30 for D completely eliminates the SC-RT-promoting ability of tRNA^Gln^_CUG_[M]. The wt and A42G mutant variants of the tRNA^Gln^_CUG_[M] iso-acceptor were expressed individually (along with EV) in wt and *rps30B-R10D* mutant yeast cells and the efficiency of SC-RT on UAG-C was measured and evaluated as described in Figure 1B – C; each box is represented by n ≥ 5 values.

Next, we individually expressed wt and mutant forms of the tRNA^Gln^_CUG_[M] iso-acceptor in Rps30/eS30 wt and mutant strains. Except for *R10A* (Fig. 5E and Extended Data Fig. 6A), none of the Rps30/eS30 mutations displayed any interaction with tRNA^Gln^_CUG_[M] mutants (Extended Data Fig. 6B and data not shown). Intriguingly, the *R10A* mutation dramatically reduced the SC-RT-inducing potential of the wt tRNA^Gln^_CUG_[M] iso-acceptor when compared to the EV control and to the A42G mutants in both tested stop codon contexts (Fig. 5E and Extended Data Fig. 6A). Importantly, the SC-RT-inducing potential of other UAG rti-tRNA, namely tRNA^Tyr^_GUA_ was not affected at all by *rps30-R10A* (Fig. 5E and Extended Data Fig. 6A). Please note that the rps30b-R10A-FLAG mutant affected neither the polysome content nor the 60S/40S ratio (Extended Data Fig. 7A - B). Also note that the reason for the overall lower SC-RT values obtained with wt tRNA^Gln^_CUG_[M] (Fig. 5E and Extended Data Fig. 6A *versus* for example Fig. 1B) is a different genetic background that was used to delete both copies of *RPS30*-encoding genes.

Finally, we reversed the charge in the R10D mutant and observed that it completely eliminated the ability of tRNA^Gln^_CUG_[M] to promote SC-RT (Fig. 5F); note this mutation did not affect basal SC-RT level of cells expressing only EV, although it led to a mild slow growth phenotype (data not shown). Taken together, these findings strongly support the view that R10 of Rps30/eS30, most probably owing to its positive charge, potentiates the SC-RT ability of tRNA^Gln^_CUG_[M] by directly contacting the 28:42 base pair of its AS.

### Interactions of the A-site ribosomal proteins and rti-tRNAs drive SC-RT

We recently discovered that shortening the AS of *S. cerevisiae* tRNA^Trp^_CCA_ from canonical 5-bp to 4-bp, while preserving its overall length, robustly (by ∼7-fold) increased the SC-RT potential of the shorter AS tRNA variant^32^. Remarkably, this “short AS” tRNA^Trp^_CCA_ naturally occurs in a handful of eukaryotes with all stop codons reassigned to sense codons^32^. Thus, together with the tRNA^Gln^_CUG_[M] described here, we found two cases of naturally occurring tRNA variants with markedly increased SC-RT potential compared to other variants encoding the same amino acids. This may suggest that it is the extra non-covalent interactions between the tRNA backbone and the A-site components, as demonstrated here for tRNA^Gln^, which generally confer all rti-tRNAs the advantage of increased SC-RT over other near-cognate tRNAs. To test that, we subjected the aforementioned C-terminus of Rps15/uS19 and the N-terminus of Rps25/eS25 (both projecting into the A site^30,31^) to a thorough mutagenesis. Subsequently, we examined the effect of the resulting mutants, together with all our Rps30/eS30 mutants, on propensity of all four rti-tRNAs to increase SC-RT on corresponding stop codons in a complex approach.

While the R10A mutation significantly reduced (by ∼30%) the SC-RT potential of the 5-bp long AS tRNA^Trp^_CCA_, as in the case of the EV control, the 4-bp long AS variant with much higher SC-RT became resistant to this reduction; it even showed a statistically insignificant but repeatedly observed modest increase (Fig. 6A). These results further support the idea R10 of eS30 can somehow monitor the geometry of the AS unless it is disturbed by mutations changing the nature of the AS bases or its length. In contrast, neither *A2G* nor *V4A* mutations of Rps30/eS30 showed any effect with any length variant of tRNA^Trp^_CCA_ (Fig. 6A). Neither these mutations nor, importantly, the *R10A* mutation affected the SC-RT potential of tRNA^Tyr^_GUA_, whose 4^th^ base pair does not obey the pyrimidine : purine rule (it has A : U at this position) (Fig. 5E, Extended Data Fig. 6, and data not shown). The tRNA^Tyr^_GUA_ thus serves as a critical specificity control.

**Figure 6.**
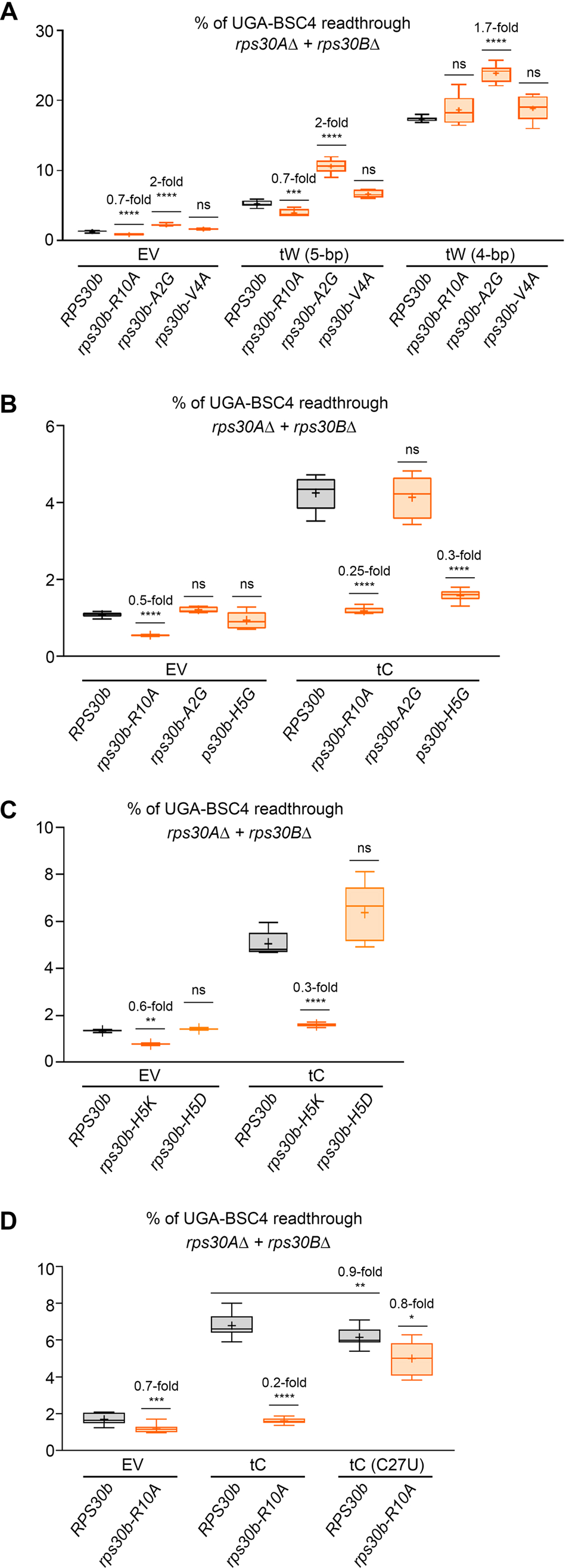
Arg 10 of Rps30/eS30 ensures incorporation of all rti-tRNAs to the A site depending on the well-defined identity of the 28:42 base pair. (A) R10 of Rps30/eS30 potentiates the SC-RT ability of the wt tRNA^Trp^_CCA_ (with the 5-bp long AS) while having no influence over its 4-bp long mutant variant. The wt and a mutant variant of tRNA^Trp^_CCA_ were expressed individually (along with EV) in wt and indicated *rps30B* mutant yeast cells and the efficiency of SC-RT on UAG-BSC4 was measured and evaluated as described in Fig. 1B - C; each box is represented by n ≥ 5 values (2 individual experiments each including ≥ 3 biological replicates). (B) H5 and R10 of Rps30/eS30 potentiates the SC-RT ability of tRNA^Cys^_GCA_. The wt tRNA^Cys^_GCA_ was expressed (along with EV) in wt and indicated *rps30B* mutant yeast cells and the efficiency of SC-RT on UGA-BSC4 was measured and evaluated as described in Fig. 1B - C; each box is represented by n ≥ 5 values. (C) Substituting H5 of Rps30/eS30 for K or D shows varying effects on the SC-RT-promoting ability of tRNA^Gln^_CUG_[M]. The wt tRNA^Cys^_GCA_ was expressed (along with EV) in wt and indicated *rps30B* mutant yeast cells and the efficiency of SC-RT on UGA-BSC4 was measured and evaluated as described in Fig. 1B - C; each box is represented by n ≥ 5 values. (D) Mutant tRNA^Cys^_GCA_ (C28U : G42) becomes insensitive to the *R10A* mutation of Rps30/eS30. The wt and mutant tRNA^Cys^_GCA_ were expressed (along with EV) in wt and the *R10A* mutant of Rps30/eS30 and the efficiency of SC-RT on UGA-BSC4 was measured and evaluated as described in Fig. 1B - C; each box is represented by n ≥ 5 values (2 individual experiments each including ≥ 3 biological replicates).

In case of tRNA^Cys^_GCA_, *R10A* and mainly *H5G* as the only mutations also reduced its SC-RT potential (Fig. 6B and data not shown). Unexpectedly, substituting basic H5 for acidic D had no effect, whereas replacing H5 for also basic K (or hydrophobic A) displayed a similar phenotype to that of the H5G mutant (Fig. 6C and data not shown). At present, we have no explanation for these observations. Remarkably, the 4^th^ base pair of wt tRNA^Cys^_GCA_ obeys the pyrimidine : purine rule (C : G) and when we disrupted its W-C character with the C27U : G41 mutation, the SC-RT-promoting activity of mutant tRNA^Cys^_GCA_^C27U^, which modestly but still significantly dropped in wt cells, became virtually insensitive to the *R10A* mutation of Rps30/eS30 (Fig. 6D). Together, we propose that the N-terminus of Rps30/eS30, specifically R10 and H5, contacts the top of the AS of tRNA^Gln^_CUG_[M], tRNA^Trp^_CCA_, and tRNA^Cys^_GCA_ to facilitate their accommodation in the A-site.

While mutations of Rps30/eS30 had no effect on the SC-RT potential of tRNA^Tyr^_GUA_, a fully viable double deletion of both Rps25/eS25-encoding genes (*RPS25A* and RPS25*B*), as well as the deletion of 12 N-terminal amino acids in *rps25-Δ2-13* expressed as a sole allele nearly eliminated its SC-RT potential (Fig. 7A and C). At the same time, these mutants had minimal impact on other three rti-tRNAs, despite their overall drastic impact on the SC-RT efficiency (Fig. 7). These findings indicate that the N-terminus of Rps25/eS25 most probably contacts specifically tRNA^Tyr^_GUA_ to achieve its increased stabilization in the A site, as in case of Rps30/eS30 and the other three rti-tRNAs.

**Figure 7.**
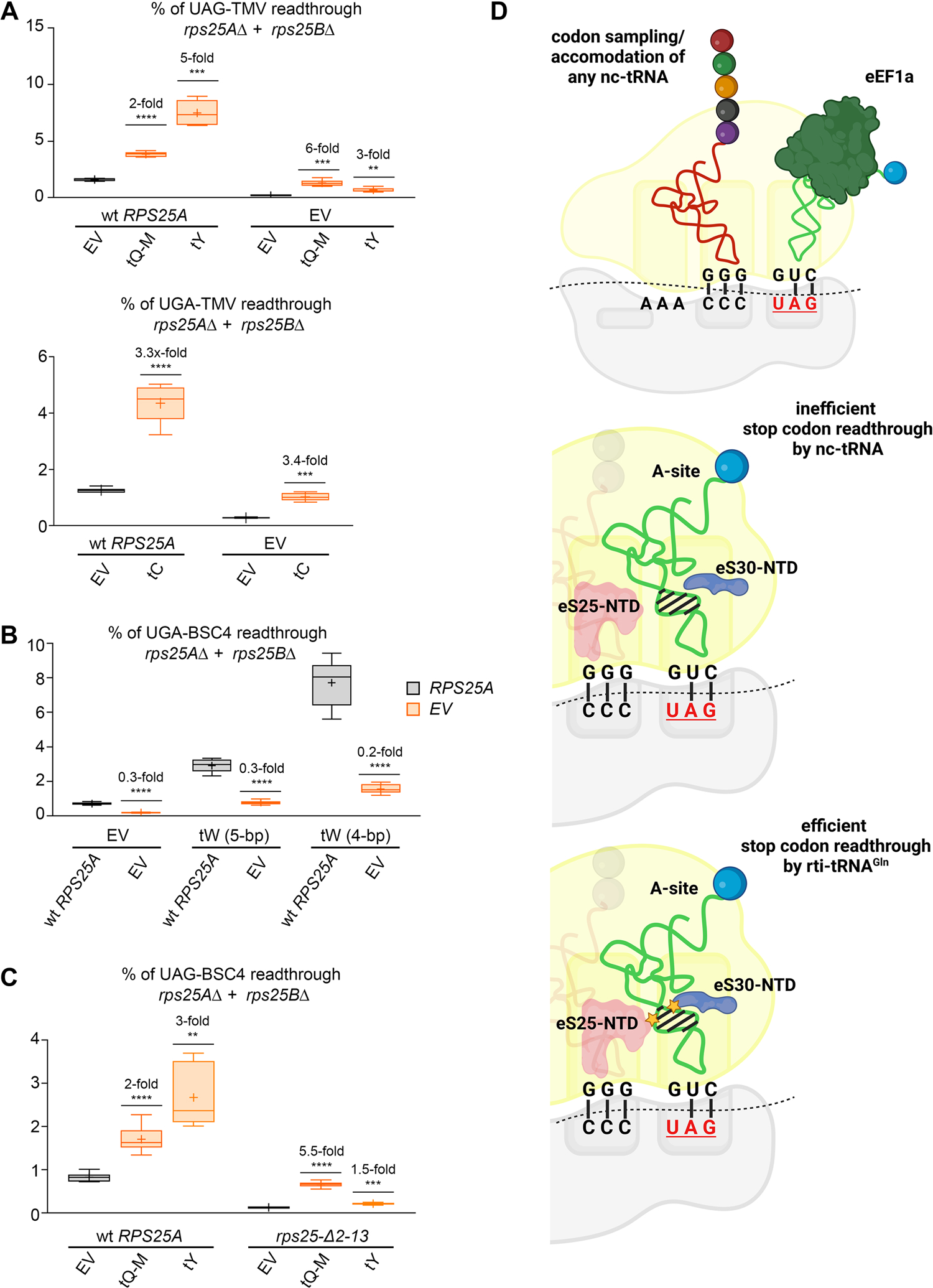
The extreme N-terminal residues of Rps25/eS25 promote incorporation of the rti-tRNA^Tyr^_GUA_ to the ribosomal A site. (A) Loss of Rps25/eS25 protein diminishes the SC-RT ability of tRNA^Tyr^_GUA_ as the only rti-tRNA. The wt tRNA^Gln^_CUG_[M], tRNA^Tyr^_GUA_, and tRNA^Cys^_GCA_ rti-tRNAs were expressed individually (along with EV) in wt and *rps25A rps25B* double deletion cells and the efficiency of SC-RT on corresponding stop codons was measured and evaluated as described in Fig. 1B – C; each box is represented by n ≥ 5 values. (B) Loss of Rps25/eS25 protein has no influence over the SC-RT potential of wt and mutant tRNA^Trp^_CCA_. The wt and a mutant variant of tRNA^Trp^_CCA_ were expressed individually (along with EV) in wt and *rps25A rps25B* double deletion cells and the efficiency of SC-RT on UGA-BSC4 was measured and evaluated as described in Fig. 1B - C; each box is represented by n = 6 values. (C) The extreme N-terminal residues of Rps25/eS25 promote incorporation of the rti-tRNA^Tyr^_GUA_ to the ribosomal A site. The wt tRNA^Gln^_CUG_[M] and tRNA^Tyr^_GUA_ were expressed individually (along with EV) in wt and *rps25-Δ2-13* mutant cells and the efficiency of SC-RT on UAG-BSC4 was measured and evaluated as described in Fig. 1B – C; each box is represented by n ≥ 5 values. (D) Model depicting the proposed molecular mechanism of efficient SC-RT. The additional contacts that selected near cognate tRNAs; i.e. rti-tRNAs, make with N-terminal (NTD) domains of Rps25/eS25 and Rps30/eS30 (marked by yellow asterisks) via their AS increase their odds of being selected during termination to promote efficient SC-RT.

Finally, robust mutational analysis of Rps15/uS19 revealed only two mutants with markedly increased SC-RT potential; in particular *rps15-R130A* and *rps15-128AAAAAAA134* (Extended Data Fig. 8A). However, we did not observe any specific interactions with either of rti-tRNAs or their mutants (Extended Data Fig. 8B - C).

## DISCUSSION

In most non-canonical genetic codes, the UAG and UAA stop codons share near-cognate tRNAs delivering the same amino acids^43^, which differ from those recognizing the UGA stop codon. In agreement with the genetic code standardly allowing wobble decoding of the 3^rd^ codon base and a well-documented tolerance for the 1^st^ base wobble decoding during recoding events, glutamine, tyrosine, and lysine are incorporated during UAR readthrough, whereas arginine, cysteine, and tryptophan are incorporated during UGA readthrough^18,21^. The efficiency of their incorporation depends on many factors, one of which is the cellular level that may vary from tissue to tissue. Nonetheless, when the relevant tRNAs were expressed individually to similar levels in *S. cerevisiae*, only four of them stood out as very potent facilitators of SC-RT, namely glutamine, cysteine, tyrosine, and tryptophan^19^. (Of note, the latter two were tested also in human cell lines with similar results^20^.) Why only these four?

Here and in our recent work^32^ we demonstrated that – at least for tRNA^Gln^_CUG_[M], tRNA^Trp^_CCA_, and tRNA^Cys^_GCA_ – the answer lies surprisingly not in the bases of their anticodon loop and/or their modifications, but most probably in their AS. In case of tRNA^Gln^_CUG_[M], the nature of the 28:42 anticodon base pair is sensed by the Arg10 of Rps30/eS30 - possibly owing to its positive charge, whose substitution to Ala greatly mitigates the ability of the tRNA^Gln^_CUG_[M] iso-acceptor to promote efficient SC-RT on UAG, and so does the alteration of the pyrimidine:purine character of its 28:42 base pair. Similarly, *R10A*, as well as *H5G/K/A* mutations also reduced the SC-RT potential of wt tRNA^Cys^_GCA_, which also obeys the pyrimidine 28 : purine 42 rule. Since we obtained fully consistent data with wt and mutant variants of *S. cerevisiae* tRNA^Gln^_CUG_[M] and *T. brucei* tRNA^Gln^_CUG_ in *T. brucei* (Fig. 2D - E), we conclude that the importance of the pyrimidine:purine nature of the 4^th^ AS base pair of at least these two rti-tRNAs is evolutionary conserved. In agreement, working with the *Escherichia coli* tRNA^Trp^_CCA_ with the artificial GUC anticodon complementary to the Glu codon CAG, it was shown that mutations disrupting base-paring between G27 and A43 (i.e. the bases forming the 5^th^ pair of its AS) turned this tRNA into a potent synthetic suppressor permitting SC-RT at UAG^44^.

Among the Rps30/eS30 N-terminal residues, mainly the basic Arg10 residue seems to monitor the length and proper base-pairing of the AS, because its Ala substitution genetically interacted also with tRNA^Trp^_CCA_ and tRNA^Cys^_GCA_ and their mutant variants. In addition, while Rps30/eS30 mutants displayed no genetic interaction with tRNA^Tyr^_GUA_, complete deletion of Rps25/eS25 or partial deletion of its N-terminal residues specifically prevented tRNA^Tyr^_GUA_ from stimulating SC-RT. These results thus further support the view that the architectures of individual tRNA are well optimized^45^. However, this kind of natural fine-tuning not only compensates for the known destabilizing effect of certain amino acids that are bound to them. Generally speaking, it also seems to ensure that specific contacts are made between the residues of the A-site RPSes and backbone bases of any tRNA to promote cognate codon/anticodon base-pairing while preventing tRNA misincorporation at non-cognate codons. In the case of rti-tRNAs, these contacts then increase their other odds to be selected during SC-RT (Fig. 7D), as we demonstrated here.

The applicability of our findings beyond the SC-RT process; i.e., generally to the canonical wobble decoding during translation elongation in, is supported by the fact that the human genome encodes ∼270 iso-decoder tRNA genes out of approximately 450 tRNA genes distributed among 49 iso-acceptor families. When the sequence diversity among iso-decoder tRNA genes was subjected to robust suppression analysis, it was concluded that the large difference in their suppression efficiency observed *in vivo* could not be explained solely by differences in tRNA folding due to their varying sequences. Rather, it was postulated that their contacts with the ribosomal components likely play a very important role in modulating their selection effectivity^46^.

With respect to Rps30/eS30, it was proposed that it may enhance the stability of a correct codon-anticodon interaction *via* its contact with the anticodon loop^24^. In the presence of a cognate aa-tRNA, the N-terminus of Rps30/eS30 becomes ordered, allowing a conserved histidine (His76, corresponding to yeast H5 studied here) to reach into a groove between the phosphate backbone of the anticodon +1 position and the two flipped-out decoding bases to form potentially stabilizing contacts. In addition, it was suggested that because this groove depends on the flipped nucleotides that accompany canonical codon-anticodon base pairing, Rps30/eS30 may preferentially stabilize cognate tRNAs to enhance discrimination^24^. Since our data strongly indicate that yeast H5, as well as R10, preferentially promote incorporation of a selected set of the nc-tRNAs to the A site, it seems that eS30 may play a dual role (especially its H5 residue). It could stabilize cognate tRNAs to enhance their discrimination during kinetic proofreading of the initial selection and/or accommodation, while stabilizing certain near-cognate tRNAs at the A site to increase their odds of being selected (Fig. 7D). Intuitively, for this specific class of tRNAs, the discriminatory role of eS30 would have to be suppressed.

In agreement with our results clearly indicating that the key AS base recognized by Arg10 is a purine at position 42 (or equivalent) base pairing with a pyrimidine at position 28 (or equivalent) of selected rti-tRNAs, all available structures of decoding complexes show that His5 and Arg10 (or the corresponding conserved residues in other organisms) are positioned close to the AS of the A/T-state tRNA^24,29,47^ (Fig. 5B). The key question is how these two residues can read out characteristic structural features (like the 4^th^ pyr28:pur42 base-pair) of the AS of some rti-tRNAs? Interestingly, the tRNA^Gln^_CUG_[M] iso-acceptor has three consecutive stacked purines in the third (G41), fourth (A42) and fifth (A43) base pair, whereas a purine-pyrimidine swap results in stacked purine-pyrimidine steps. Molecular dynamics studies demonstrated that duplex RNA (A-helix form) with purine:pyrimidine steps is less stiff than the homogenous duplexes^48^. This may be explained by the interstrand steric clashes between adjacent purine bases that are stepwise stacked^49^. We thus speculate that such a stiffer structure of the AS may facilitate highly specific interaction with positively charged Arg10 of eS30, because Arg10 would most likely contact the RNA backbone and the likelihood of forming a stable salt bridge with the phosphate backbone may be higher if the A-helix is stiffer. Yet, further studies, e.g., using cryo-electron microscopy of readthrough complexes will be needed to reveal the exact molecular basis of these interactions in the ribosomal decoding center.

Interestingly, while only a mild increase in SC-RT was observed for the *rps30-V4A* mutation on UAG and UGA, a very strong increase (more than 4-fold) was observed on UAA; similarly, while virtually no effect was observed for the *rps30-R10A* mutation on UAA and UAG, a significant drop in SC-RT (by ∼30-50%) was observed on UGA (Fig. 5D and Extended Data Fig. 5). Strikingly, an extreme N-terminal residue R8 of human Rps30/eS30, corresponding to *S. cerevisiae* R10 studied here, was directly cross-linked to the +4 position/base of the mRNA (+1 position being the first nucleotide of the stop codon in the A site) in a pre-termination ribosome^50^. This is exactly the base shown to be accommodated in the eukaryotic-specific U-turn-like conformation of the stop codon in the decoding pocket, which is formed when the N-terminal domain (NTD) of eRF1 contacts the ribosome^51,52^. It is also the base, which has a significant impact on the efficiency of SC-RT^2,8–11^. In addition, the N-terminal tail of Rps30/eS30 is, along with Rps31/eS31 and several 18S rRNA sites, known to interact with the eRF1-NTD^53^. In fact, it is believed that upon eRF1 binding, the N-terminal tail of Rps30/eS30 is shielded by the eRF1-NTD from interacting with the backbone of the stop codon as observed in a pre-termination ribosome free of eRFs^50^. Thus, we propose that Rps30/eS30, through its N-terminus, not only discriminates between individual tRNAs, but also somehow recognizes the identity of a stop codon to facilitate its recognition by eRF1, thereby controlling the balance between translation termination and SC-RT.

Besides Rps30/eS30, the AS of the A-site aa-tRNA also appears to be in contact with the 12 to 15 C-terminal residues of Rps15/uS19, depending on the species. In addition, this region makes electrostatic interactions with the P-site tRNA, the rRNA helices h44 and h69, as well as with the +4 position/base of the mRNA like R8 (*S. cerevisiae* R10) of Rps30/eS30^24,50,54^. However, we found no interactions between the C-terminal mutations of this ribosomal protein and any of rti-tRNAs. This can be explained by two previous observations. Analysis of tRNAs associated with 40S subunits and 80S ribosomes showed that the deletion of the C-terminal pentadecapeptide fragment of Rps15/uS19 does not affect binding of aa-tRNA at the A site^55^. Furthermore, interactions of the Rps15/uS19 C-terminal tail with the mRNA or the AS of the A-site tRNA are not observed in the eEF1A-bound decoding state^54^. Therefore, it seems plausible that the Rps15/uS19 establishes contact with the AS of the A-site tRNA post eEF1A dissociation and kinetic proofreading of the decoded tRNA, which contributes to efficientH5 accommodation of the A-site tRNA, thereby maintaining the optimal rate of elongation. However, this contact may not – in the case of near cognate tRNAs – be of any dramatic importance, as these tRNAs would require energetic support mainly during the initial codon sampling phase, which occurs still in the presence of eEF1A.

The N-terminal extension of the Rps25/eS25 was originally thought to be positioned between the P and E sites^56,57^. However, available structures (PDB: 7RR5) suggest that it could stretch as far as to the decoding center of the A site, where it may interact with the 1^st^ through 3^rd^ base pair of the tRNA AS. So far, the non-essential Rps25/eS25 has been implicated primarily in the translation of a specific population of mRNAs by specialized ribosomes, as well as in the cap-independent translation initiation *via* internal ribosome entry sites (IRES) and ribosome shunting^58,59^. However, our data with tRNA^Tyr^_GUA_ suggest an additional role for this protein in the control of SC-RT and, highly likely, in the aa-tRNA selection process during elongation (Fig. 7D).

Given that nonsense mutations account for approximately 11% of inherited genetic disorders in humans^60^, prioritizing SC-RT for nonsense mutation correction represents an attractive approach for treating these disorders that could substantially alleviate human disease^61^. Noteworthy, the use of readthrough-inducing drugs (RTIDs) was first tested in patients with cystic fibrosis caused by a nonsense mutation in the CFTR gene^62^. However, the RTIDs are often highly toxic and many patients are non-responsive to the treatment for unknown reasons. Therefore, the application of specialized rti-tRNAs engineered to become highly potent for SC-RT offers a tempting complement to the RTID-based drug therapy. Indeed, a combination of both approaches could mitigate the toxicity of RTIDs by reducing their administered dose, as well as maximize targeted SC-RT while minimizing off-target miscoding^20^. Another application of such tRNAs could be in our combat against human pathogens whose life cycle is dependent on SC-RT, such as various viruses. We believe that the work described here significantly contributes to the understanding of the rules dictating efficacy of SC-RT, which despite extensive study remain elusive.

## ONLINE METHODS

### *S. cerevisiae* and *T. brucei* strains and plasmids

The lists and descriptions of all strains, plasmids, primers and GeneArt Strings DNA Fragments (Invitrogen) used throughout this study can be found in Supplementary Tables 1 – 6 in the Supplementary Information.

### Stop codon readthrough assays

The majority of stop codon readthrough assays in this study were performed using a standard bicistronic reporter construct bearing a *Renilla* luciferase gene followed by an in-frame *Firefly* luciferase gene, originally developed by^63^. The two genes are separated by either a tetranucleotide termination signal (UGA-C) or, for control purposes, the CAA sense codon, followed by cytosine. In indicated cases, the termination signal and*/*or the following nucleotide context was modified. Note that mRNA levels of the reporters bearing the stop signal between *Renilla* and *Firefly* genes do not differ from the CAA sense control^19^. This system avoids possible artifacts connected to the changes in the efficiency of translation initiation associated with the nonsense-mediated decay pathway^64^, because both Renilla and Firefly enzymes initiate translation from the same AUG codon. All experiments and data analyses were carried out according to the Microtiter plate-based dual luciferase assay (Promega). Readthrough measurements were conducted from at least three biological replicates (n ≥ 3) and each experiment was repeated at least three times. The readthrough values are represented by the box-and-whisker plot with the mean value marked (+) and whiskers going from minimal to maximal value. The box always extends from the 25th to 75th percentiles. The line inside of the box is plotted at the median. Statistical significance was determined by the unpaired, two-tailed Welch’s *t* test. The readthrough calculations with raw data of *Firefly* and *Renilla* measurements are given in the Source Data for Figs. 1-7 and Extended Data Figs. 2, 5, 6, and 8.

### Northern blot analysis

To examine expression levels of studied tRNAs, total RNA from yeast was isolated by the Quick RNA miniprep assay and total RNA from T. brucei was isolated as previously described ^65^. One microgram (yeast) or 10µg (*T. brucei*) was separated on a denaturing 8M urea 8% polyacrylamide gel, and electroblotted to Zeta-probe membranes. The membrane was then UV cross-linked for 1 min followed by Northern hybridization according to the manufacturer (Bio-Rad) using with 32P-labeled oligonucleotides. Following hybridization, membranes were exposed overnight on a phosphoimager screen. Blots were analyzed using a Typhoon scanner and the ImageQuant TL software (GE Healthcare).

### Polysomal gradient analysis

The *S. cerevisiae* strains bearing the wt or mutant Rps30/eS30 were cultured in the YPD media at 30 °C to an OD_600_ ∼0.5. Cycloheximide (50 μg/ml) was added 5 min prior to harvesting by centrifugation at 1,455 g for 5 min at 4 °C. The whole cell extracts (WCEs) were prepared in the GA-HEPES buffer composed of 10 mM HEPES (pH 7.5), 62.5 mM KCl and 10 mM MgCl_2_ supplemented with 50 µg/ml cycloheximide, 1 mM DTT, 1 mM PMSF, and 1 tablet of cOmplete EDTA-free per 25 ml of buffer. Cells were broken using glass beads (∼1/2 volume) by the Bead Beater Homogenizer. After centrifugation at 15,871 g for 5 min at 4 °C, the supernatant was transferred into a pre-cooled 1.5 ml tube and cleared by centrifugation at 21,130 g at 4 °C for 10 min.

For polysome profile analysis, fifteen A_260_ units of WCEs were separated by high velocity sedimentation on a 5–45% sucrose gradient in the GA-HEPES buffer by centrifugation at 39,000 rpm at 4 °C for 2.5 h in the SW41Ti rotor (Beckman Coulter). The resulting gradients were scanned at A_254_ using Teledyne ISCO UA-6 UV/VIS gradient detector.

For the 60S/40S ratio analysis, fifteen A_260_ units of WCEs with addition of 50 mM EDTA were separated and analyzed as described above in the GA-HEPES buffer without MgCl_2_ to visualize free 40S and 60S ribosomal subunits.

### Bioinformatics

The genomes and protein-coding sequences were collected from NCBI and are listed in Extended Data Excel File 1A. The tRNAs^Gln^ were predicted by ARAGORN v1.2.38^66^ and tRNAscan-SE v2.0.5^67^. Identical sequences of one organism were deduplicated using an in-house python script (https://github.com/kikinocka/ngs/blob/master/py_scripts/deduplicate_sequence_seqs.py). This means that there are still some identical sequences in the dataset but originating from different assemblies. Sequences from ciliates with canonical genetic code (alignment_file_1) and those with reassigned TAR codons (alignment_file_2) were aligned separately using MAFFT G-INS-i v7.458^68^. Sequences with a longer variable loop were manually refined using secondary structures obtained from ARAGORN and/or tRNAscan-SE. The base pair corresponding to 28:42 pair was extracted from the alignments (Extended Data Excel File 1B) using an in-house python script (https://github.com/kikinocka/ngs/blob/master/py_scripts/nt_pair_aln.py).

Codon usage of each protein-coding sequence was determined using an in-house python script (https://github.com/kikinocka/ngs/blob/master/py_scripts/codon_usage.py). The total sum of each Gln codon was extracted for each organism as numbers observed; the expected numbers were calculated as (CAA+CAG+TAA+TAG)/4, and χ^2^ test was calculated (Extended Data Excel File 1C).

### REPORTING SUMMARY

Further information on research design is available in the Nature Research Reporting Summary linked to this paper.

## DATA AVAILABILITY

All data generated during this study are included in this published article (and its Supplementary Information files). Plasmids and yeast strains generated in this study are available upon request.

## CODE AVAILABILITY

Nothing to report.

## ACKNOWLEDGEMENTS

We are grateful to Ivana Malcová (Institute of Microbiology CAS, Prague) and Jonathan D. Dinman (University of Maryland) for providing various plasmids and to Sebastian Leidel (University of Bern) for providing yeast strains individually deleted for genes required for the U_34_ modification. This work was supported by the Czech Science Foundation grants 20-00579S (to L.S.V.) and 20-11585S (to Zd.Pa.), Lead Agency (DFG & CSF) grant 23-08669L (to L.S.V. and Zd. Pa.), and the Praemium Academiae grant provided by the Czech Academy of Sciences (to L.S.V.), and the Charles University Grant Agency project GA UK 1192819 (to Zu.Pa.). Computational resources were provided by the e-INFRA CZ project (ID:90140), supported by the Ministry of Education, Youth and Sports of the Czech Republic.

## AUTHOR CONTRIBUTIONS

Zu.Pa., P.B. and L.S.V. conceived and designed the project.

Zu.Pa., P.M., K.P., A.R., K.Z., I.M.D., A.K. and Zd. Pa. carried out all experiments and performed the data analysis.

Zu.Pa., P.M., Zd. Pa., P.B. and L.S.V. interpreted the results. L.S.V. wrote the paper with input from Zu.Pa., P.M., K.Z., T.B., J.L., Zd.Pa. and P.B.

## MATERIALS & CORRESPONDENCE

Correspondence and requests for materials should be addressed to.

Supplementary information is available for this paper.

## EXTENDED DATA FIGURE LEGENDS

**Extended Data Figure 1.**
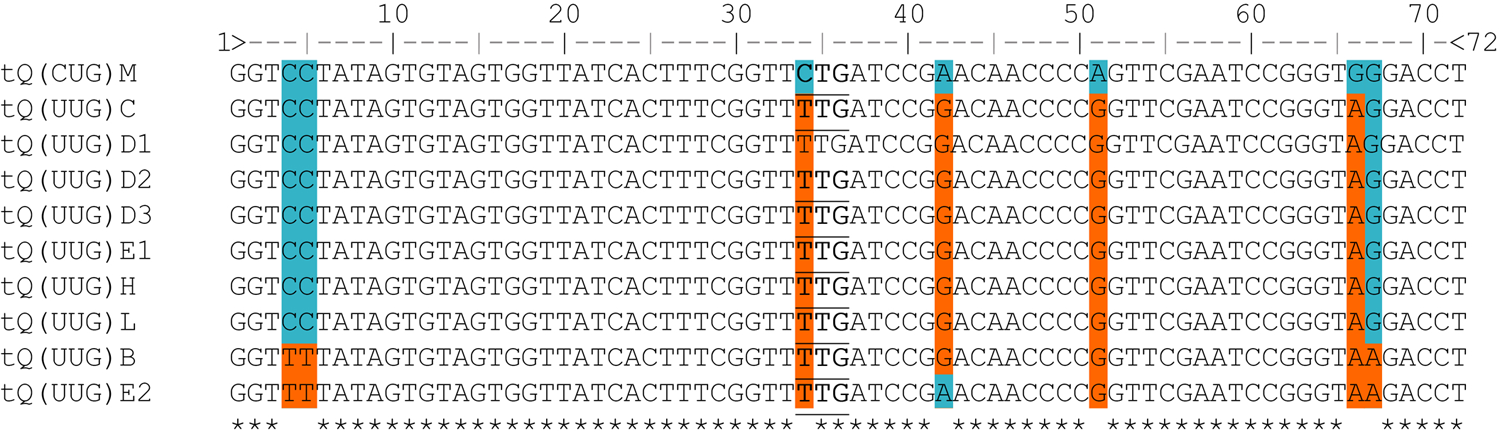
Multiple sequence alignment of *S. cerevisiae* tRNA^Gln^ iso-acceptors retrieved from the GtRNAdb^69^. Variable positions among iso-acceptors are highlighted by colors; same residues with tRNA^Gln^_CUG_[M] in blue, differing residues in orange.

**Extended Data Figure 2.**
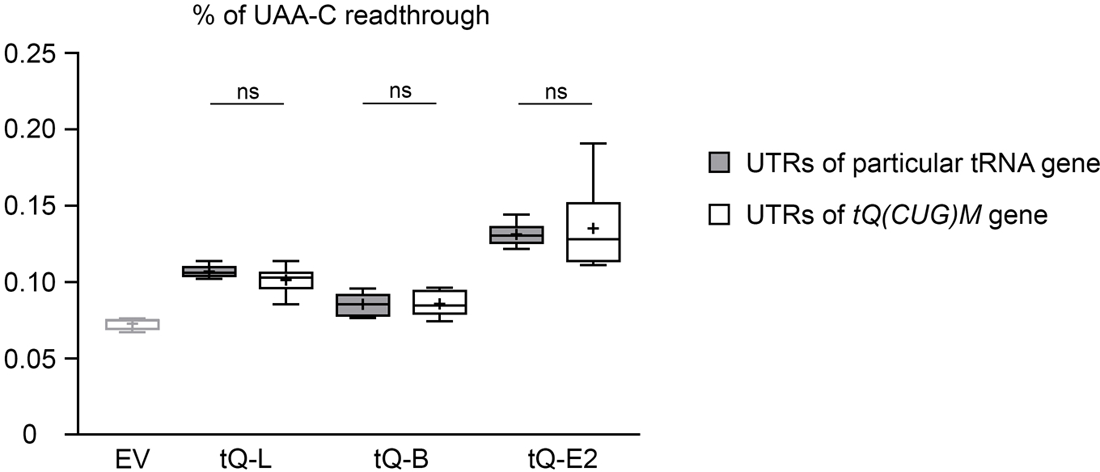
UTRs of tRNA^Gln^ iso-acceptors do not influence SC-RT efficiency. The wt tRNA^Gln^_UUG_ iso-decoders flanked by either the genuine or “tRNA^Gln^_CUG_[M]” UTRs were expressed individually (along with EV) in wt yeast cells and the efficiency of SC-RT on UAA-C was measured and evaluated as described in Fig. 1B – C; each box is represented by n ≥ 5 values.

**Extended Data Figure 3.**
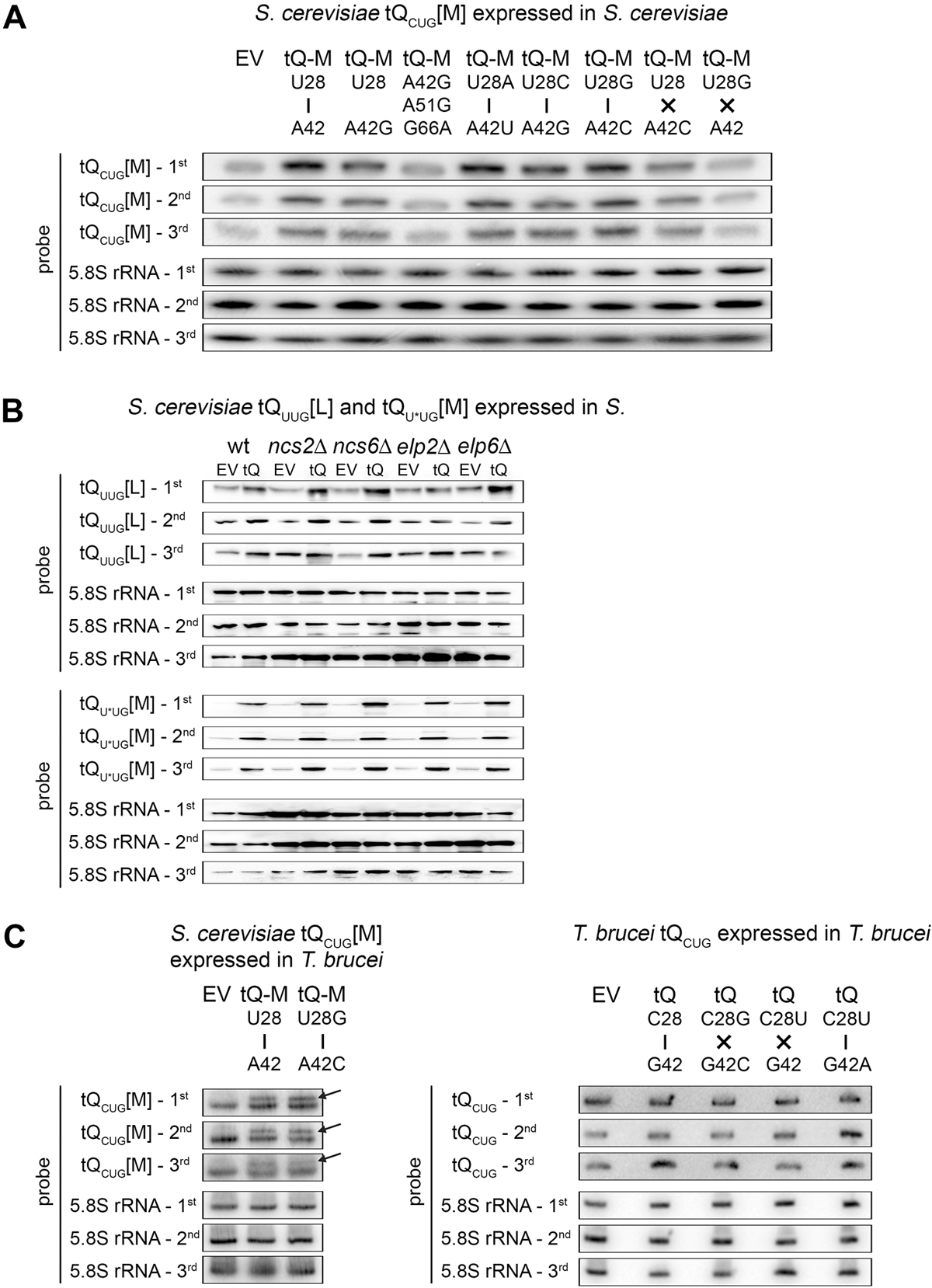
Expression levels of tRNA^Gln^ variants in *S. cerevisiae* and *T. brucei*. (A) Expression of the *S. cerevisiae* tRNA^Gln^_CUG_ [M] variants in *S. cerevisiae*. Total RNA was isolated from the wt *S. cerevisiae* strain PBH156 expressing plasmid-borne high copy wt or indicated mutant variants of tRNA^Gln^_CUG_ (tQ-M; indicated at the top of each panel). Five micrograms of total RNA was resolved on the urea gel, transferred onto a nylon membrane and hybridized with a specific [γ^32^P]ATP labeled probe against tRNA^Gln^_CUG_ [M]. The 5.8S rRNA was used as a loading control. Three biological replicates are shown; for quantifications, see Source Data Extended Data Fig. 3. (B) Expression of the *S. cerevisiae* tRNA^Gln^_UUG_[L] and tRNA^Gln^_U*UG_[M] variants in *S. cerevisiae*. Total RNA was isolated from the indicated wt and mutant *S. cerevisiae* strains expressing either empty vector or tRNA^Gln^_UUG_[L] (upper 6 panels) or tRNA^Gln^_U*UG_[M] (lower 6 panels). One microgram of total RNA was resolved on the urea gel, transferred onto a nylon membrane and hybridized with a specific DIG-labeled probe against tRNA^Gln^_CUG_ [M]. The 5.8S rRNA was used as a loading control. For details see the main text and Figure 1E and F. Three biological replicates are shown; for quantifications, see Source Data Extended Data Fig. 3. (C) Expression of the *S. cerevisiae* tRNA^Gln^_CUG_[M] and *T. brucei* tRNA^Gln^_CUG_ variants in *T. brucei*. (left) Total RNA was extracted from *T. brucei* strain 29-13 expressing the genome integrated plasmid pLEW100 bearing (tQ) variants from *S. cerevisiae* (indicated at the top of each panel). Ten µg of RNA was resolved by urea-PAGE, followed by the northern blot analysis using [γ^32^P]ATP labeled oligonucleotide specific for both tQ variants; Empty vector (EV) served as a negative control. 5.8S rRNA was used as a loading control. Three biological replicates are shown. (right) Total RNA was extracted from *T. brucei* strain 29-13 expressing the genome integrated plasmid pLEW100 bearing (tQ) variants from *T. brucei* (indicated at the top of each panel). Ten µg of RNA was resolved by urea-PAGE, followed by the northern blot analysis using [γ^32^P]ATP labeled oligonucleotide specific for tQ variants; Empty vector (EV) served as a negative control. 5.8S rRNA was used as a loading control. Three biological replicates are shown.

**Extended Data Figure 4.**
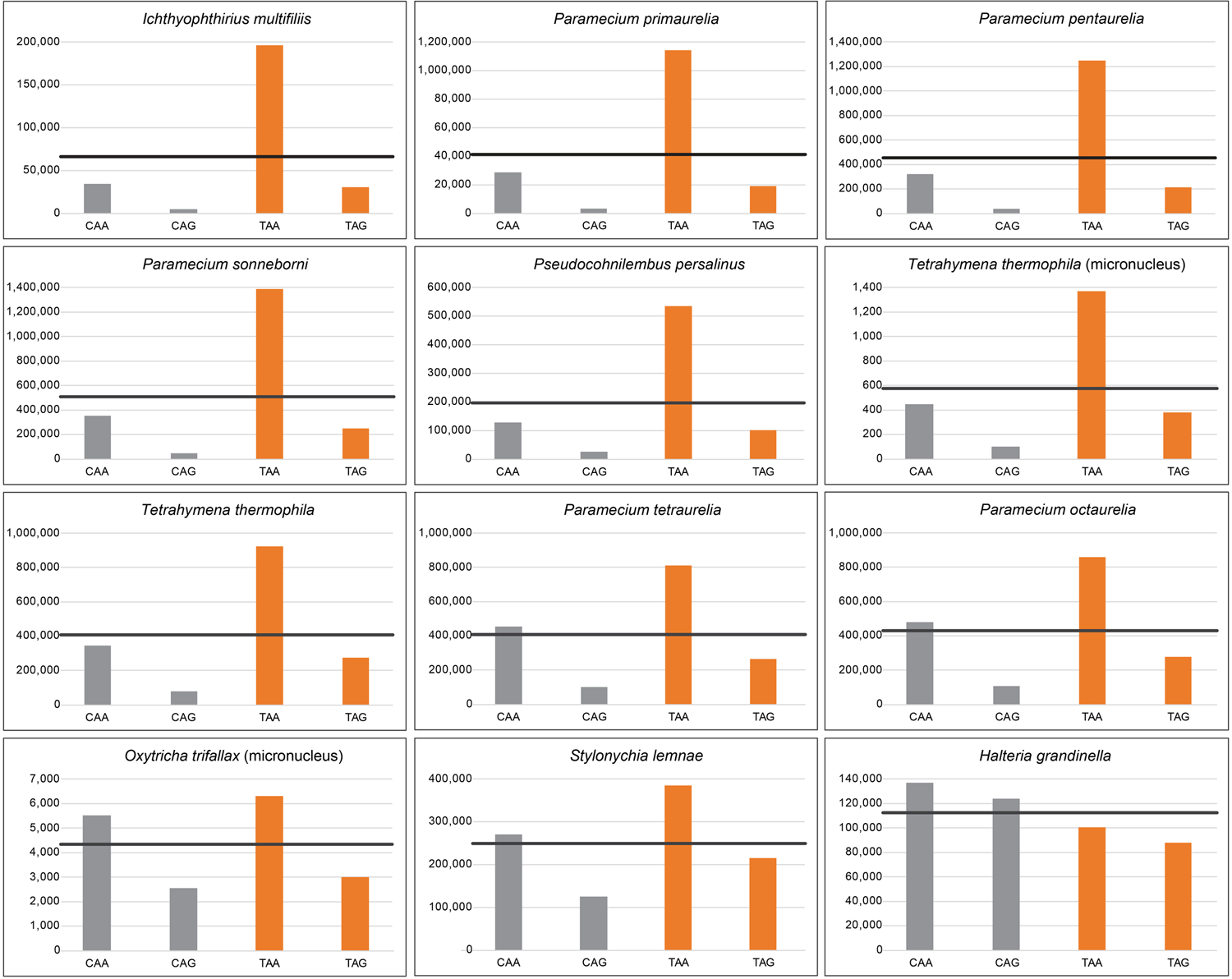
Codon usage of Gln codons in ciliates with reassigned genetic code. Protein-coding sequences originating from macronucleus (unless stated otherwise) were obtained from NCBI (Extended Data Supplementary Excel file 1A). Codon usage of all Gln codons (CAA, CAG, TAA, and TAG) was determined (Extended Data Supplementary Excel file 1C), and the observed numbers were visualized as bar plots. Expected numbers ([CAA+CAG+TAA+TAG]/4) are represented by horizontal black lines. Canonical and reassigned Gln codons are in grey and orange, respectively.

**Extended Data Figure 5.**
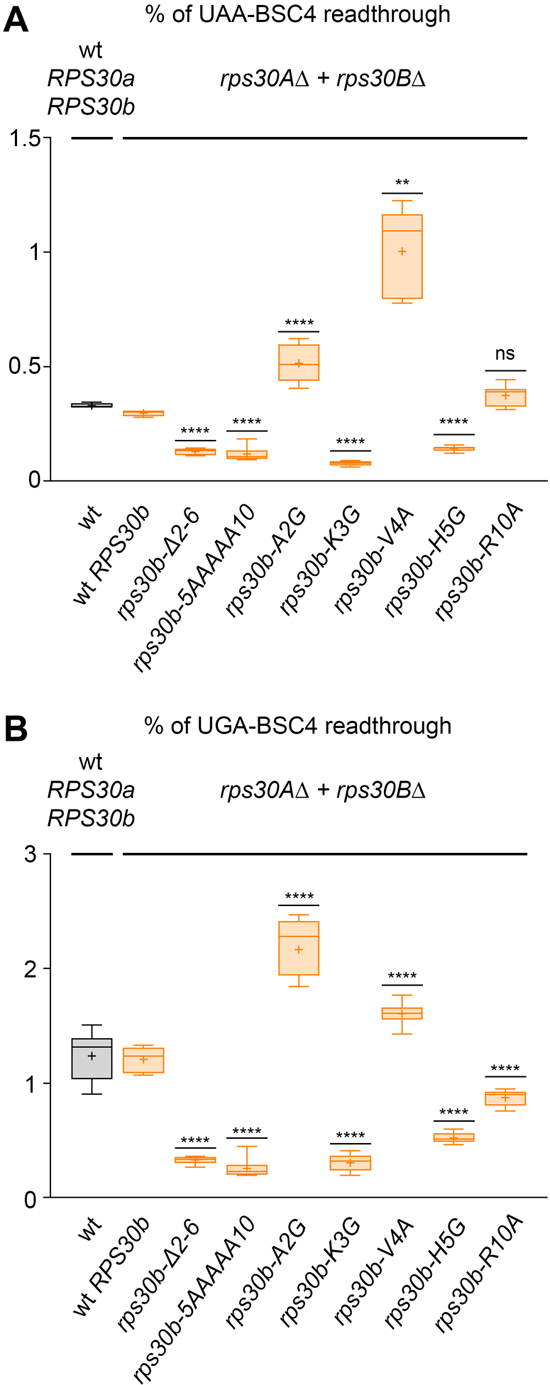
Rps30/eS30 mutants differentially modulate the SC-RT efficiency at the UAA and UGA stop codon. (A – B). The wt and indicated Rps30/eS30 mutant yeast cells were examined for efficiency of SC-RT on (A) UAA-BSC4 or (B) UGA-BSC4 as described in Fig. 1B – C; each box is represented by n ≥ 6 values (2 individual experiments each including n ≥ 3 biological replicates).

**Extended Data Figure 6.**
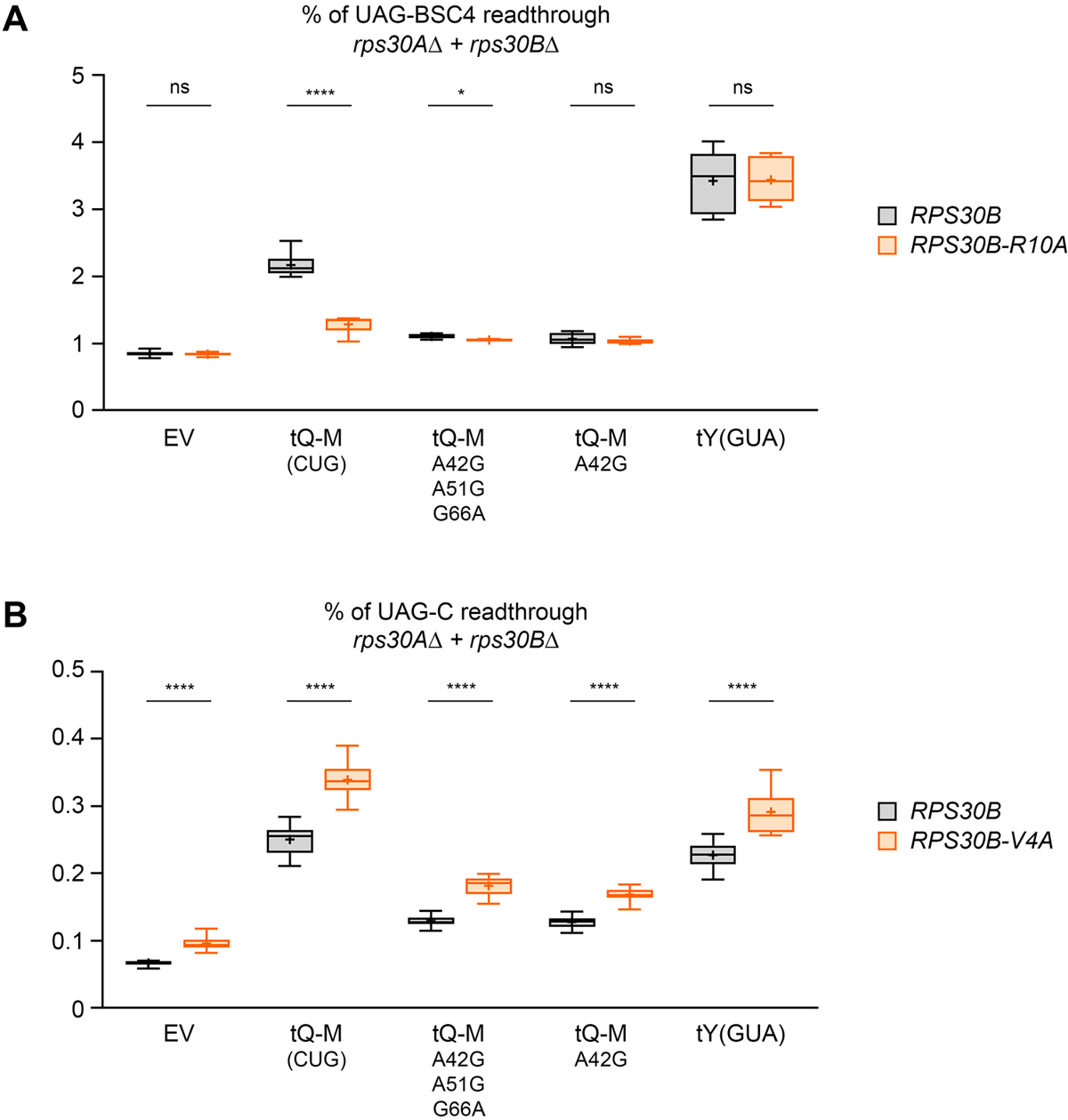
Impact of the Rps30/eS30 mutants on incorporation of the tRNA^Gln^_CUG_[M] iso-acceptor into the A site. (A) R10 of Rps30/eS30 may potentiate the SC-RT-promoting ability of tRNA^Gln^_CUG_[M] by directly contacting the 28:42 base pair of its AS. The wt and indicated mutant variants of the tRNA^Gln^_CUG_[M] iso-acceptor, as well as control tRNA^Tyr^_GUA_ were expressed individually (along with EV) in wt and *rps30B-R10A* mutant yeast cells and the efficiency of SC-RT on UAG-BSC4 was measured and evaluated as described in Fig. 1B - C; each box is represented by n ≥ 5 values. (B) V4 of Rps30/eS30 has no effect on the SC-RT-promoting ability of tRNA^Gln^_CUG_[M]. The same as in panel A only the *V4A* instead of *R10A* mutation of Rps30/eS30 was examined on UAG-C; each box is represented by n ≥ 11 values (2 individual experiments each including n ≥ 5 biological replicates).

**Extended Data Figure 7.**
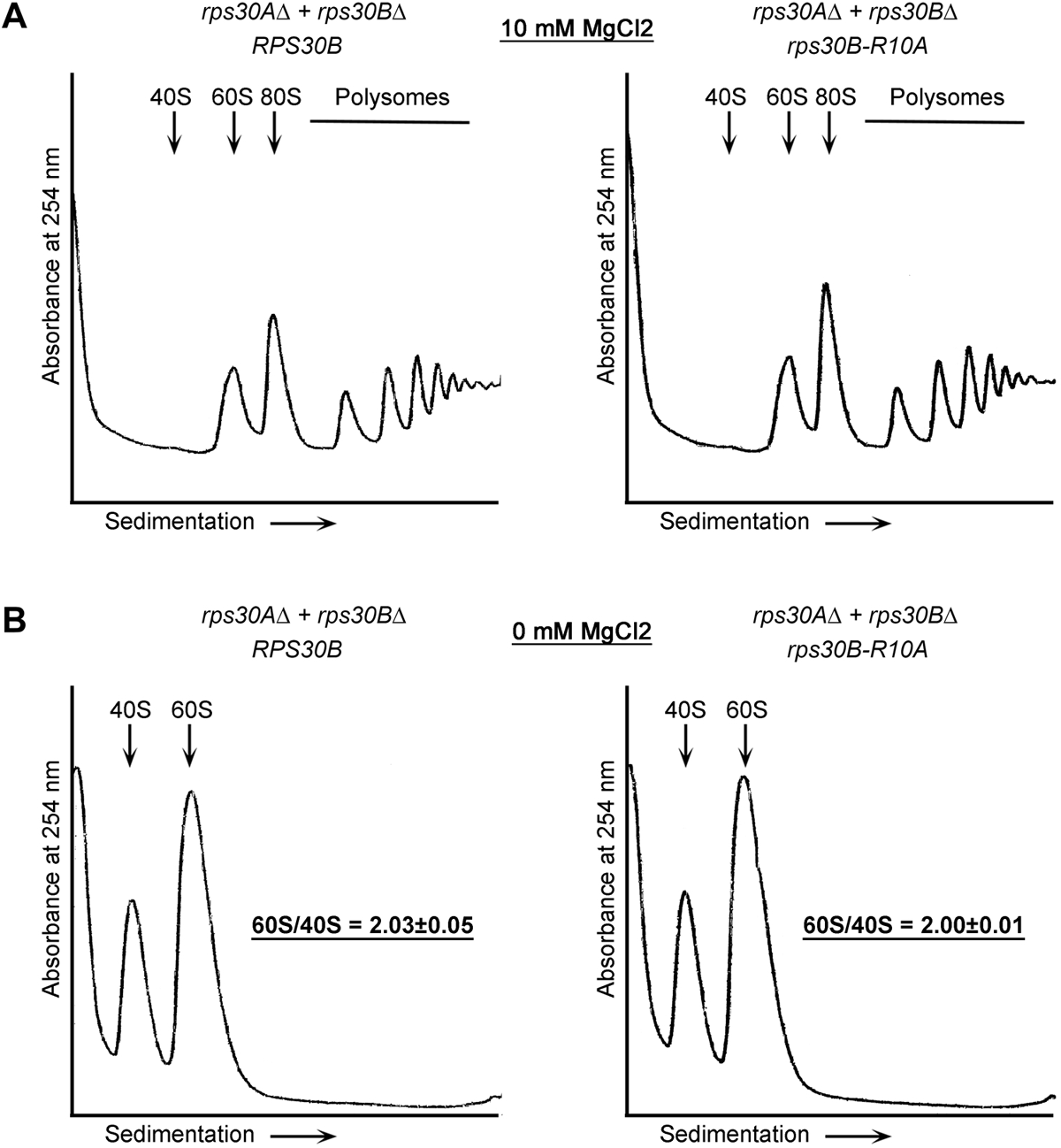
The Rps30/eS30 mutation *R10A* has no detectable impact either on the polysome content or on the 60S/40S ratio. (A – B) Polysomal profiles of wt vs. *rps30B-R10A* mutant yeast cells (A) in the presence of 10mM MgCl_2_ or (B) in the absence of MgCl_2_. Yeast cells bearing the indicated wt or mutant Rps30/eS30 were grown in the YPD medium at 30 °C to an OD_600_ ∼0.5 and processed for polysomal gradient analysis as described in Methods. Positions of 40S, 60S and 80S species are indicated by arrows; polysomes by a horizontal line. (B) The 60S/40S ratio represents the result from three independent experiments ± SD.

**Extended Data Figure 8.**
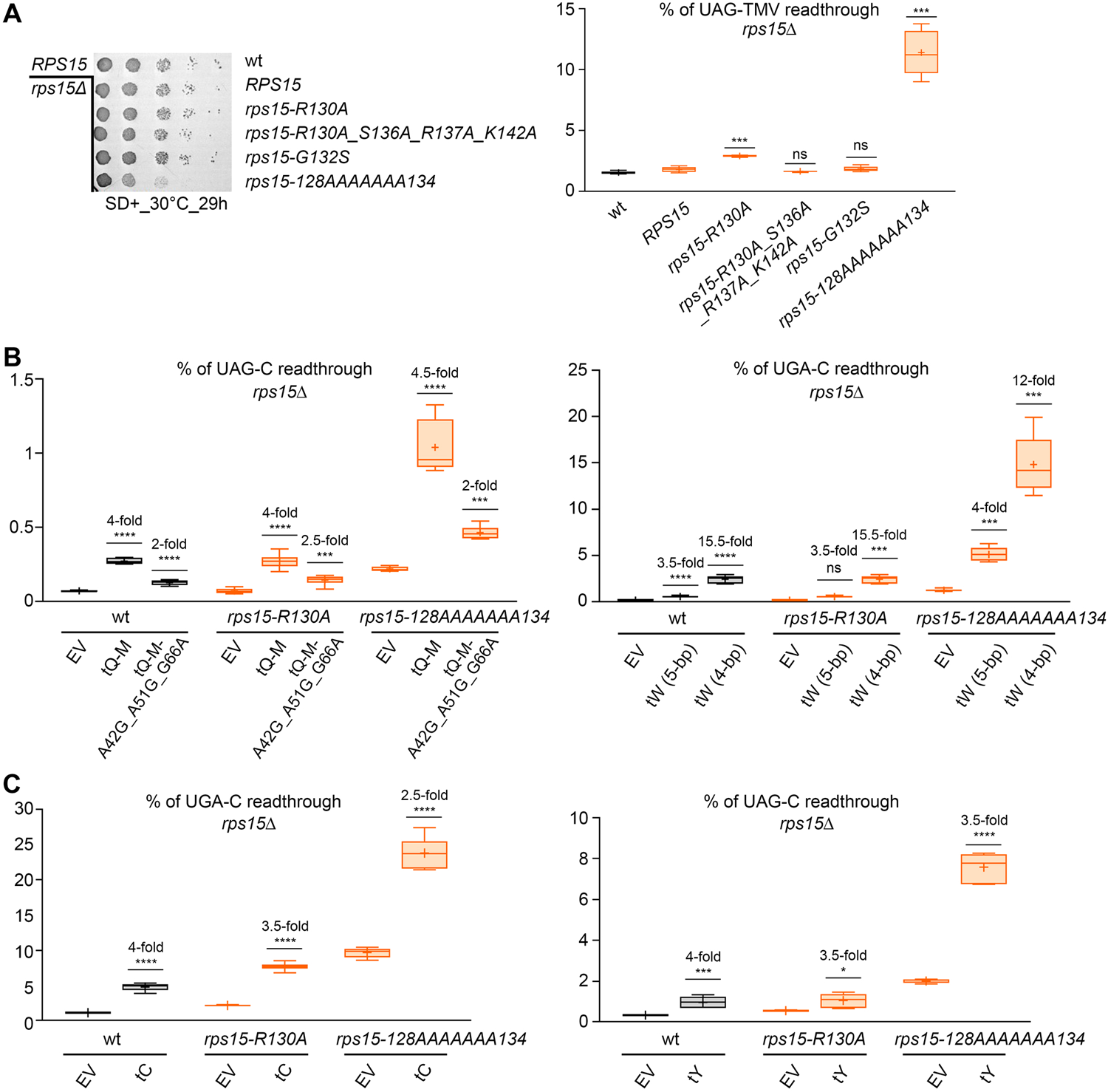
The extreme C-terminal residues of Rps19/uS19 do not influence the SC-RT efficiency of any rti-tRNAs or their mutant variants. (A) Growth rate and stop codon readthrough analysis of the C-terminal substitution mutants of Rps15/uS19. The wt and indicated Rps15/uS19 mutant yeast cells were spotted in five serial 10-fold dilutions on minimal media supplemented with required amino acids and incubated for 29 hours at 30 °C. The same cells were tested for efficiency of SC-RT in the UAG-TMV stop codon context as described in Fig. 1B - C; each box is represented by n≥ 5 values (2 individual experiments each including ≥ 2 biological replicates). (B) The extreme C-terminal residues of Rps15/uS19 do not influence the SC-RT efficiency of tRNA^Gln^_CUG_[M] and tRNA^Trp^_CCA_ or their mutant variants. The wt and indicated mutant variants of tRNA^Gln^_CUG_[M] and tRNA^Trp^_CAA_ were expressed individually (along with EV) in wt and indicated *rps15* mutant yeast cells and the efficiency of SC-RT on UAG-C and UGA-C, respectively, was measured and evaluated as described in Fig. 1B – C; each box is represented by n≥ 5 values (2 individual experiments each including ≥ 2 biological replicates). (C) The extreme C-terminal residues of Rps15/uS19 do not influence the SC-RT efficiency of tRNA^Tyr^_GUA_ and tRNA^Cys^_GCA_. The wt tRNA^Cys^_GCA_ and tRNA^Tyr^_GUA_ were expressed individually (along with EV) in wt and indicated *rps15* mutant yeast cells and the efficiency of SC-RT on UGA-C and UAG-C, respectively, was measured and evaluated as described in Fig. 1B – C; each box is represented by n ≥ 5 values.

## EXTENDED DATA TABLE TITLES AND FOOTNOTES

**Extended Data Supplementary Excel file 1.** Bioinformatics analyses of Gln tRNAs and codons. (A) List of species used for the analyses. Genetic codes and accession numbers of genomic assemblies that were obtained from NCBI are given in columns C and D, respectively. Numbers of protein-coding genes are listed in column E and links to their sources in F. (B) Listed are 28:42 base pairs for each tRNA^Gln^ for each ciliate species. (C) Summary counts (Observed) for Gln codons are listed for each species. Expected numbers were calculated as (CAA+CAG+TAA+TAG)/4, except for *Stentor coeruleus* that employs standard genetic code ^70^, and thus the expected numbers were calculated as (CAA+CAG)/2. From these numbers, bar plots were generated (see Extended Data Fig. 4) and χ^2^ was calculated.

**Extended Data Table 1.** The 28:42 base pair counts in tRNAs^Gln^ of ciliates with canonical genetic code (UAR = stop) and those with reassigned UAR codons (UAR = Q).

