## Supplementary material for "Ribosomal A-site interactions with near-cognate tRNAs drive stop codon readthrough": Sup methods and tables

### SUPPLEMENTARY MATERIALS AND METHODS

#### Construction of yeast strains

List of all yeast strains used throughout this study can be found in the **Supplementary Table 1.**

To create PBH159 strain, PBH140 was transformed with *YCp22-g/TIF35-KLF-TRP1* (B670) and the uracil auxotrophy was regained by growing the cells on SD plates containing 5-fluoro-orotic acid (5-FOA) resulting in loss of *YEp-TIF35-U*RA3 plasmid.

To create ZH20 strain, H541 / L1751 was transformed with the gene disruption cassette created as a PCR product (template *FA6a-kanMX4* [1]; primers 6z and 7z) to obtain *rps25AΔ::kanMX4*. The resulting strain was transformed firstly with *pRS316-RPS25A-URA3* (ZPB43) and then with the gene disruption cassette created as a PCR product (template pZC4 carrying *lox2272-natNT2-lox2272* [2]; primers 8z and 9z) to obtain *rps25BΔ::natNT2*. Then the uracil auxotrophy was regained by growing the cells on SD plates containing 5-FOA resulting in loss of ZPB43.

To create ZH31 strain, H541 / L1751 was firstly transformed with the plasmid *pRS426-tQ(C*UG)B-URA3* (PBB243) and then with the gene disruption cassette created as a PCR product (template *FA6a-kanMX4* [1]; primers 18z and 19z) to obtain *tQ(CUG)MΔ::kanMX4*.

To create ZH40 strain, ZH31 was transformed with *YEplac181-tQ(CUG)M-LEU2* (ZPB58) and the uracil auxotrophy was regained by growing the cells on SD plates containing 5-FOA resulting in loss of *pRS426-tQ(C*UG)B-URA3* (PBB243) plasmid.

To create ZH43 strain, ZH31 was transformed with *YEplac181-tQ(CUG)M_A42G-LEU2* (ZPB59) and the uracil auxotrophy was regained by growing the cells on SD plates containing 5-FOA resulting in loss of *pRS426-tQ(C*UG)B-URA3* (PBB243) plasmid.

To create ZH51 strain, H541 / L1751 was transformed firstly with *YEplac195-pRPS28B+FLAG-RPS15-URA3* (ZPB53) and then with the gene disruption cassette created as a PCR product (template pZC4 carrying *lox2272-natNT2-lox2272* [2]; primers 31z and 32z) to obtain *rps15Δ::natNT2*.

To create ZH54 strain, H541 / L1751 was transformed with *YEplac112-TRP1.*

To create ZH56 strain, ZH20 was transformed with the plasmid *YEplac112-TRP1*.

To create ZH60 strain, ZH20 was transformed with the plasmid *YEplac112-RPS25A wt-FLAG-TRP1* (ZPB73).

To create ZH62 strain, ZH51 was transformed with *YEplac112-pRPS28B+FLAG-RPS15-TRP1* (ZPB51) and the uracil auxotrophy was regained by growing the cells on SD plates containing 5-FOA resulting in loss of *YEplac195-pRPS28B+FLAG-RPS15-URA3* (ZPB53) plasmid.

To create ZH65 strain, ZH51 was transformed with *YEplac112-pRPS28B+FLAG-RPS15_R130A-TRP1* (ZPB52) and the uracil auxotrophy was regained by growing the cells on SD plates containing 5-FOA resulting in loss of *YEplac195-pRPS28B+FLAG-RPS15-URA3* (ZPB53) plasmid.

To create ZH68 strain, ZH51 was transformed with *YEplac112-pRPS28B+FLAG-RPS15_R130A_S136A_R137A_K142A-TRP1* (ZPB64) and the uracil auxotrophy was regained by growing the cells on SD plates containing 5-FOA resulting in loss of *YEplac195-pRPS28B+FLAG-RPS15-URA3* (ZPB53) plasmid.

To create ZH71 strain, ZH51 was transformed with *YEplac112-pRPS28B+FLAG-RPS15_G132S-TRP1* (ZPB66) and the uracil auxotrophy was regained by growing the cells on SD plates containing 5-FOA resulting in loss of *YEplac195-pRPS28B+FLAG-RPS15-URA3* (ZPB53) plasmid.

To create ZH77 strain, ZH51 was transformed with *YEplac112-pRPS28B+FLAG-RPS15_128AAAAAAA134-TRP1* (ZPB65) and the uracil auxotrophy was regained by growing the cells on SD plates containing 5-FOA resulting in loss of *YEplac195-pRPS28B+FLAG-RPS15-URA3* (ZPB53) plasmid.

To create ZH80 strain, ZH20 was transformed with the plasmid *YEplac112-RPS25A_Δ2-13-FLAG* (ZPB74).

To create ZH112 strain, H541 / L1751 was transformed with the gene disruption cassette created as a PCR product (template pUG73 carrying *loxP–LEU2–loxP* [3]; primers 48z and 49z) to obtain *rps30BΔ::LEU2*. Then the *LEU2* disruption cassette was removed by Cre recombinase expressed from the plasmid pSH47 as described in [3]). The uracil auxotrophy was regained by growing the cells on SD plates containing 5-FOA resulting in loss of pSH47. The resulting strain was transformed firstly with *YEplac195-RPS30B-URA3* (ZPB94) and then with the gene disruption cassette created as a PCR product (template pZC4 carrying *lox2272-natNT2-lox2272* [2]; primers 46z and 47z) to obtain *rps30AΔ::natNT2*.

To create ZH117 strain, ZH112 was transformed with *YEplac112-RPS30B-FLAG-TRP1* (ZPB95) and the uracil auxotrophy was regained by growing the cells on SD plates containing 5-FOA resulting in loss of *YEplac195-RPS30B-URA3* (ZPB94) plasmid.

To create ZH121 strain, ZH112 was transformed with *YEplac112-RPS30B-Δ2-6-FLAG-TRP1* (ZPB96) and the uracil auxotrophy was regained by growing the cells on SD plates containing 5-FOA resulting in loss of *YEplac195-RPS30B-URA3* (ZPB94) plasmid.

To create ZH125 strain, ZH112 was transformed with *YEplac112-RPS30B-5AAAAAA10-FLAG-TRP1* (ZPB97) and the uracil auxotrophy was regained by growing the cells on SD plates containing 5-FOA resulting in loss of *YEplac195-RPS30B-URA3* (ZPB94) plasmid.

To create ZH129 strain, ZH112 was transformed with *YEplac112-RPS30B-A2G-FLAG-TRP1* (ZPB98) and the uracil auxotrophy was regained by growing the cells on SD plates containing 5-FOA resulting in loss of *YEplac195-RPS30B-URA3* (ZPB94) plasmid.

To create ZH133 strain, ZH112 was transformed with *YEplac112-RPS30B-K3G-FLAG-TRP1* (ZPB99) and the uracil auxotrophy was regained by growing the cells on SD plates containing 5-FOA resulting in loss of *YEplac195-RPS30B-URA3* (ZPB94) plasmid.

To create ZH137 strain, ZH112 was transformed with *YEplac112-RPS30B-H5G-FLAG-TRP1* (ZPB100) and the uracil auxotrophy was regained by growing the cells on SD plates containing 5-FOA resulting in loss of *YEplac195-RPS30B-URA3* (ZPB94) plasmid.

To create ZH141 strain, ZH112 was transformed with *YEplac112-RPS30B-R10A-FLAG-TRP1* (ZPB101) and the uracil auxotrophy was regained by growing the cells on SD plates containing 5-FOA resulting in loss of *YEplac195-RPS30B-URA3* (ZPB94) plasmid.

To create ZH267 strain, ZH112 was transformed with *YEplac112-RPS30B-V4A-FLAG-TRP1* (ZPB145) and the uracil auxotrophy was regained by growing the cells on SD plates containing 5-FOA resulting in loss of *YEplac195-RPS30B-URA3* (ZPB94) plasmid.

To create ZH275 strain, ZH112 was transformed with *YEplac112-RPS30B-H5D-FLAG-TRP1* (PMB4) and the uracil auxotrophy was regained by growing the cells on SD plates containing 5-FOA resulting in loss of *YEplac195-RPS30B-URA3* (ZPB94) plasmid.

To create ZH276 strain, ZH112 was transformed with *YEplac112-RPS30B-H5K-FLAG-TRP1* (PMB5) and the uracil auxotrophy was regained by growing the cells on SD plates containing 5-FOA resulting in loss of *YEplac195-RPS30B-URA3* (ZPB94) plasmid.

To create ZH278 strain, ZH112 was transformed with *YEplac112-RPS30B-R10D-FLAG-TRP1* (PMB6) and the uracil auxotrophy was regained by growing the cells on SD plates containing 5-FOA resulting in loss of *YEplac195-RPS30B-URA3* (ZPB94) plasmid

**Construction of yeast plasmids**

List of all plasmids, PCR primers and GeneArt Strings DNA Fragments (Invitrogen) used throughout this study can be found in **Supplementary Tables 2 – 4**.

ZPB67 was created by inserting the *Sal*I-*Not*I digested PCR product obtained with primers 42z and PB43 using pTH477 as a template into *Sal*I-*Not*I digested pTH477.

ZPB68 was created by inserting the *Sal*I-*Not*I digested PCR product obtained with primers 43z and PB43 using pTH477 as a template into *Sal*I-*Not*I digested pTH477.

ZPB70 was created by inserting *AlwN*I-*Nsi*I digested ZPB67 (4457 bp) into *AlwN*I-*Nsi*I digested YEplac181 (4174 bp).

ZPB71 was created by inserting *AlwN*I-*Nsi*I digested ZPB68 (4457 bp) into *AlwN*I-*Nsi*I digested YEplac181 (4174 bp).

ZPB78 was created by inserting the *Sal*I-*Not*I digested PCR product obtained with primers 44z and PB43 using PBB37 as a template into *Sal*I-*Not*I digested PBB37.

ZPB80 was created by inserting the *Sal*I-*Not*I digested PCR product obtained with primers PB78 and PB43 using PBB37 as a template into *Sal*I-*Not*I digested PBB37.

PBB238 was created using fusion PCR with PBB150 serving as a template and the following combinations of primers: (i) PB140 + PB189 and (ii) PB188 + PB141. Primers PB140 and PB141 were then used in the third reaction using PCR products from the first and second reactions in the 1:1 ratio as templates. The resulting PCR product was digested with *Xho*I and *BamH*I and inserted into *Xho*I-*BamH*I cut PBB150.

PBB239 was created using fusion PCR with PBB150 serving as a template and the following combinations of primers: (i) PB140 + PB191 and (ii) PB190 + PB141. Primers PB140 and PB141 were then used in the third reaction using PCR products from the first and second reactions in the 1:1 ratio as templates. The resulting PCR product was digested with *Xho*I and *BamH*I and inserted into *Xho*I-*BamH*I cut PBB150.

PBB240 was created using fusion PCR with PBB150 serving as a template and the following combinations of primers: (i) PB140 + PB193 and (ii) PB192 + PB141. Primers PB140 and PB141 were then used in the third reaction using PCR products from the first and second reactions in the 1:1 ratio as templates. The resulting PCR product was digested with *Xho*I and *BamH*I and inserted into *Xho*I-*BamH*I cut PBB150.

PBB241 was created using fusion PCR with PBB150 serving as a template and the following combinations of primers: (i) PB140 + PB195 and (ii) PB194 + PB141. Primers PB140 and PB141 were then used in the third reaction using PCR products from the first and second reactions in the 1:1 ratio as templates. The resulting PCR product was digested with *Xho*I and *BamH*I and inserted into *Xho*I-*BamH*I cut PBB150.

PBB242 was created using fusion PCR with PBB150 serving as a template and the following combinations of primers: (i) PB140 + PB197 and (ii) PB196 + PB141. Primers PB140 and PB141 were then used in the third reaction using PCR products from the first and second reactions in the 1:1 ratio as templates. The resulting PCR product was digested with *Xho*I and *BamH*I and inserted into *Xho*I-*BamH*I cut PBB150.

PBB243 was created using fusion PCR with PBB148 serving as a template and the following combinations of primers: (i) PB134 + PB199 and (ii) PB198 + PB135. Primers PB134 and PB135 were then used in the third reaction using PCR products from the first and second reactions in the 1:1 ratio as templates. The resulting PCR product was digested with *Xho*I and *BamH*I and inserted into *Xho*I-*BamH*I cut PBB150.

ZPB4 was created by inserting *Xho*I-*BamH*I digested *tQ(UUG)B_tQ-M UTRs* (GeneArt String DNA Fragment; Invitrogen) into *Xho*I-*BamH*I digested PBB150.

ZPB5 was created by inserting *Xho*I-*BamH*I digested *tQ(UUG)E2_tQ-M UTRs* (GeneArt String DNA Fragment; Invitrogen) into *Xho*I-*BamH*I digested PBB150.

ZPB6 was created using fusion PCR with PBB150 serving as a template and the following combinations of primers: (i) PB140 + 5z and (ii) PB190 + PB141. Primers PB140 and PB141 were then used in the third reaction using PCR products from the first and second reactions in the 1:1 ratio as templates. The resulting PCR product was digested with *Xho*I and *BamH*I and inserted into *Xho*I-*BamH*I cut PBB150.

ZPB22 was created by inserting *Xho*I-*BamH*I digested *tQ(UUG)M_A51G* (GeneArt String DNA Fragment; Invitrogen) into *Xho*I-*BamH*I digested PBB150.

ZPB23 was created by inserting *Xho*I-*BamH*I digested *tQ(UUG)M_A42G_G66A* (GeneArt String DNA Fragment; Invitrogen) into *Xho*I-*BamH*I digested PBB150.

ZPB25 was created by inserting *Xho*I-*BamH*I digested *tQ(UUG)M_A42G* (GeneArt String DNA Fragment; Invitrogen) into *Xho*I-*BamH*I digested PBB150.

ZPB26 was created by inserting *Xho*I-*BamH*I digested *tQ(UUG)M_G66A* (GeneArt String DNA Fragment; Invitrogen) into *Xho*I-*BamH*I digested PBB150.

ZPB28 was created by inserting *Xho*I-*BamH*I digested *tQ(CUG)M_A42G_G66A* (GeneArt String DNA Fragment; Invitrogen) into *Xho*I-*BamH*I digested PBB150.

ZPB36 was created by inserting *Xho*I-*BamH*I digested *tQ(CUG)M_U28A_A42G* (GeneArt String DNA Fragment; Invitrogen) into *Xho*I-*BamH*I digested PBB150.

ZPB37 was created by inserting *Xho*I-*BamH*I digested *tQ(CUG)M_U28C_A42G* (GeneArt String DNA Fragment; Invitrogen) into *Xho*I-*BamH*I digested PBB150.

ZPB42 was created by inserting *Xho*I-*BamH*I digested *tQ(CUG)M_U28G_A42C* (GeneArt String DNA Fragment; Invitrogen) into *Xho*I-*BamH*I digested PBB150.

ZPB43 was created by inserting the *BamH*I-*EcoR*I digested PCR product obtained with primers 12z and 13z using genomic DNA obtained from the yeast strain PBH156 as template into *BamH*I-*EcoR*I digested pRS316.

ZPB44 was created by inserting the *BamH*I-*SphI* digested PCR product obtained with primers 14z and 15z using ZPB43 as template into *BamH*I-*Sph*I digested B937.

ZPB45 was created by inserting *EcoR*I-*Sph*I digested *ZPB44* into *EcoR*I-*Sph*I digested YEplac112.

ZPB51 was created by inserting the *BamH*I-*Sph*I digested PCR product obtained with primers PB247 and PB248 using genomic DNA obtained from the yeast strain as template into *BamH*I-*Sph*I digested ZPB45.

ZPB52 was created using fusion PCR with ZPB50 serving as a template and the following combinations of primers: (i) PB249 + PB248 and (ii) PB247 + PB250. Primers PB247 and PB248 were then used in the third reaction using PCR products from the first and second reactions in the 1:1 ratio as templates. The resulting PCR product was digested with *Sph*I and *BamH*I and inserted into *Sph*I-*BamH*I cut ZPB45.

ZPB53 was created by inserting the *BamH*I-*SphI* digested PCR product obtained with primers PB247 and PB248 using using genomic DNA obtained from the yeast strain as template into *BamH*I-*Sph*I digested B937.

ZPB58 was created by inserting *Kpn*I-*BamH*I digested *PBB150 (189 bp)* into *Kpn*I-*BamH*I digested YEplac181 (5728 bp).

ZPB59 was created by inserting *Kpn*I-*BamH*I digested *PBB240 (189 bp)* into *Kpn*I-*BamH*I digested YEplac181 (5728 bp).

ZPB64 was created using fusion PCR with ZPB50 serving as a template and the following combinations of primers: (i) 35z + PB248 and (ii) PB247 + 36z. Primers PB247 and PB248 were then used in the third reaction using PCR products from the first and second reactions in the 1:1 ratio as templates. The resulting PCR product was digested with *Sph*I and *BamH*I and inserted into *Sph*I-*BamH*I cut ZPB45.

ZPB65 was created using fusion PCR with ZPB50 serving as a template and the following combinations of primers: (i) 37z + PB248 and (ii) PB247 + 38z. Primers

PB247 and PB248 were then used in the third reaction using PCR products from the first and second reactions in the 1:1 ratio as templates. The resulting PCR product was digested with *Sph*I and *BamH*I and inserted into *Sph*I-*BamH*I cut ZPB45.

ZPB66 was created using fusion PCR with ZPB50 serving as a template and the following combinations of primers: (i) 39z + PB248 and (ii) PB247 + 40z. Primers PB247 and PB248 were then used in the third reaction using PCR products from the first and second reactions in the 1:1 ratio as templates. The resulting PCR product was digested with *Sph*I and *BamH*I and inserted into *Sph*I-*BamH*I cut ZPB45.

ZPB73 was created using fusion PCR with ZPB43 serving as a template and the following combinations of primers: (i) 12z + 30z and (ii) 29z + 13z. Primers 12z and 13z were then used in the third reaction using PCR products from the first and second reactions in the 1:1 ratio as templates. The resulting PCR product was digested with *EcoR*I and *BamH*I and inserted into *EcoR*I-*BamH*I cut YEplac112.

ZPB74 was created using fusion PCR with ZPB73 serving as a template and the following combinations of primers: (i) 12z + 22z and (ii) 23z + 13z. Primers 12z and 13z were then used in the third reaction using PCR products from the first and second reactions in the 1:1 ratio as templates. The resulting PCR product was digested with *EcoR*I and *BamH*I and inserted into *EcoR*I-*BamH*I cut YEplac112.

ZPB82 was created by inserting *Xho*I-*BamH*I digested *tQ(CUG)M_A42C* (GeneArt String DNA Fragment; Invitrogen) into *Xho*I-*BamH*I digested PBB150.

ZPB83 was created by inserting *Xho*I-*BamH*I digested *tQ(UUG)M_U28G* (GeneArt String DNA Fragment; Invitrogen) into *Xho*I-*BamH*I digested PBB150.

ZPB94 was created by inserting the *BamH*I-*EcoR*I digested PCR product obtained with primers 52z and 53z using genomic DNA obtained from the yeast strain H541 / L1751 as template into *BamH*I-*EcoR*I digested YEplac195.

ZPB95 was created using fusion PCR with ZPB94 serving as a template and the following combinations of primers: (i) 52z + 55z and (ii) 53z + 54z. Primers 52z and 53z were then used in the third reaction using PCR products from the first and second reactions in the 1:1 ratio as templates. The resulting PCR product was digested with *EcoR*I and *BamH*I and inserted into *EcoR*I-*BamH*I cut YEplac112.

ZPB96 was created using fusion PCR with ZPB95 serving as a template and the following combinations of primers: (i) 52z + 63z and (ii) 53z + 62z. Primers 52z and 53z were then used in the third reaction using PCR products from the first and second reactions in the 1:1 ratio as templates. The resulting PCR product was digested with *EcoR*I and *BamH*I and inserted into *EcoR*I-*BamH*I cut YEplac112.

ZPB97 was created using fusion PCR with ZPB95 serving as a template and the following combinations of primers: (i) 52z + 65z and (ii) 53z + 64z. Primers 52z and 53z were then used in the third reaction using PCR products from the first and second reactions in the 1:1 ratio as templates. The resulting PCR product was digested with *EcoR*I and *BamH*I and inserted into *EcoR*I-*BamH*I cut YEplac112.

ZPB98 was created using fusion PCR with ZPB95 serving as a template and the following combinations of primers: (i) 52z + 67z and (ii) 53z + 66z. Primers 52z and 53z were then used in the third reaction using PCR products from the first and second reactions in the 1:1 ratio as templates. The resulting PCR product was digested with *EcoR*I and *BamH*I and inserted into *EcoR*I-*BamH*I cut YEplac112.

ZPB99 was created using fusion PCR with ZPB95 serving as a template and the following combinations of primers: (i) 52z + 69z and (ii) 53z + 68z. Primers 52z and 53z were then used in the third reaction using PCR products from the first and second reactions in the 1:1 ratio as templates. The resulting PCR product was digested with *EcoR*I and *BamH*I and inserted into *EcoR*I-*BamH*I cut YEplac112.

ZPB100 was created using fusion PCR with ZPB95 serving as a template and the following combinations of primers: (i) 52z + 71z and (ii) 53z + 70z. Primers 52z and 53z were then used in the third reaction using PCR products from the first and second reactions in the 1:1 ratio as templates. The resulting PCR product was digested with *EcoR*I and *BamH*I and inserted into *EcoR*I-*BamH*I cut YEplac112.

ZPB101 was created using fusion PCR with ZPB95 serving as a template and the following combinations of primers: (i) 52z + 73z and (ii) 53z + 72z. Primers 52z and 53z were then used in the third reaction using PCR products from the first and second reactions in the 1:1 ratio as templates. The resulting PCR product was digested with *EcoR*I and *BamH*I and inserted into *EcoR*I-*BamH*I cut YEplac112.

ZPB145 was created using fusion PCR with ZPB95 serving as a template and the following combinations of primers: (i) 52z + 112z and (ii) 53z + 111z. Primers 52z and 53z were then used in the third reaction using PCR products from the first and second reactions in the 1:1 ratio as templates. The resulting PCR product was digested with *EcoR*I and *BamH*I and inserted into *EcoR*I-*BamH*I cut YEplac112.

ZPB147 was created by inserting *Xho*I-*BamH*I digested *tC(GCA)P1_C27U_G41* (GeneArt String DNA Fragment; Invitrogen) into *Xho*I-*BamH*I digested PBB150.

PMB4 was created using fusion PCR with ZPB95 serving as a template and the following combinations of primers: (i) 52z + 10p and (ii) 53z + 9p. Primers 52z and 53z were then used in the third reaction using PCR products from the first and second reactions in the 1:1 ratio as templates. The resulting PCR product was digested with *EcoR*I and *BamH*I and inserted into *EcoR*I-*BamH*I cut YEplac112.

PMB5 was created using fusion PCR with ZPB95 serving as a template and the following combinations of primers: (i) 52z + 8p and (ii) 53z + 7p. Primers 52z and 53z were then used in the third reaction using PCR products from the first and second reactions in the 1:1 ratio as templates. The resulting PCR product was digested with *EcoR*I and *BamH*I and inserted into *EcoR*I-*BamH*I cut YEplac112.

PMB6 was created using fusion PCR with ZPB95 serving as a template and the following combinations of primers: (i) 52z + 4p and (ii) 53z + 3p. Primers 52z and 53z were then used in the third reaction using PCR products from the first and second reactions in the 1:1 ratio as templates. The resulting PCR product was digested with *EcoR*I and *BamH*I and inserted into *EcoR*I-*BamH*I cut YEplac112.

**Plasmids used for *T. brucei* transfection**

All plasmids were purchased synthetized from Gene Universal into pLew100[4] for expression in *T. brucei* 29-13 cells[5]. The corresponding sequences of the constructs used for *T. brucei* transfection (5' and 3´UTRs and wt or mutated t-RNA$^{Gln}$ can be found in **Supplementary Table 5.**

**Construction of *T. brucei* recombinant cell lines**

A list of *T. brucei* transgenic cell lines generated here can be found at **Supplementary Table 6.**

To generate TbtQ(CUG)U28A42:UAG-C cell line, TbUAG-C, a 29-13 procyclic strain bearing the dual luciferase cassette with an in-frame UAG-C stop codon[6] was transfected with the plasmid pLew100:tQ(CUG)U28A42, a pLew100 derivative bearing the M t-RNA$^{Gln}$ iso-acceptor flanked by *T. brucei* endogenous Gln tRNA 5' and 3' UTRs. Stably transfected cell lines were obtained by selection with Phleomycin marker encoded in pLew100 (Phleo) at a final concentration of 2.5 μg/ml.

To generate TbtQ(CUG)U28A42:DLC-C cell line, TbDLC-C, a 29-13 procyclic strain bearing the dual luciferase cassette without in-frame stop codons[6] was transfected with the plasmid pLew100:tQ(CUG)U28A42, a pLew100 derivative bearing the M tRNA$^{Gln}$ iso-acceptor flanked by *T. brucei* endogenous Gln t-RNA 5' and 3' UTRs. Stably transfected cell lines were selected as above.

To generate Tb*Q(CUG)U28GXA42C:UAG-C cell line, TbUAG-C, a 29-13 procyclic strain bearing the dual luciferase cassette with an in-frame UAG-C stop codon was transfected with plasmid pLew100:*Q(CUG)U28GXA42C, a pLew100 derivative bearing the mutated tRNA$^{Gln}$ iso-acceptor flanked by *T. brucei* endogenous Gln tRNA 5' and 3' UTRs. Stably transfected cell lines were selected as above.

To generate Tb*Q(CUG)U28GXA42C:DLC-C cell line, TbDLC-C, a 29-13 procyclic strain bearing the dual luciferase cassette without in-frame stop codon was transfected with plasmid pLew100:*Q(CUG)U28XA42G, a pLew100 derivative bearing the mutated t-RNA$^{Gln}$ iso-acceptor flanked by *T. brucei* endogenous Gln tRNA 5' and 3' UTRs. Stably transfected cell lines were selected as above.

To generate TbQ(CUG)C28G42:DLC-C cell line, TbDLC-C, a 29-13 procyclic strain bearing the dual luciferase cassette without in-frame stop codon was transfected with plasmid pLew100:Q(CUG)C28G42C, a pLew100 derivative bearing the WT t-RNA$^{Gln}$ iso-acceptor flanked by *T. brucei* endogenous Gln tRNA 5' and 3' UTRs. Stably transfected cell lines were selected as above.

To generate TbQ*(CUG)C28GXG42C:DLC-C cell line, TbDLC-C, a 29-13 procyclic strain bearing the dual luciferase cassette without in-frame stop codon was transfected with plasmid pLew100:Q*(CUG)C28GXG42C, a pLew100 derivative bearing the mutated t-RNA$^{Gln}$ iso-acceptor flanked by *T. brucei* endogenous Gln tRNA 5' and 3' UTRs. Stably transfected cell lines were selected as above.

To generate TbQ*(CUG)C28UXG42A:DLC-C cell line, TbDLC-C, a 29-13 procyclic strain bearing the dual luciferase cassette without in-frame stop codon was transfected with plasmid pLew100:Q*(CUG)C28UXG42A, a pLew100 derivative bearing the mutated t-RNA$^{Gln}$ iso-acceptor flanked by *T. brucei* endogenous Gln tRNA 5' and 3' UTRs. Stably transfected cell lines were selected as above.

To generate TbQ*(CUG)C28UXG42:DLC-C cell line, TbDLC-C, a 29-13 procyclic strain bearing the dual luciferase cassette without in-frame stop codon was transfected with plasmid pLew100:Q*(CUG)C28UXG42, a pLew100 derivative bearing the mutated t-$RNA^{Gln}$ iso-acceptor flanked by *T. brucei* endogenous Gln tRNA 5' and 3' UTRs. Stably transfected cell lines were selected as above.

**Generation of *T. brucei* transgenic lines**

For the analysis of SC-RT in *T. brucei*, procyclic cells of 29-13 strain[5] bearing either UAG-C or DLC cassette without stop codon were transfected with approximately 10 µg of either pLew100tQ(CUG)U28A42 or pLew100*Q(CUG)U28GXA42C (synthetized by Gene Universal) NotI digested plasmids (Supplementary Table 5) using Amaxa electroporator (program X-114). Approximately 16 hours after transfection, cells were placed into selective media supplemented with Phleomycin at a final concentration of 2.5 µg/ml and plated into 24-well plates to obtain semi-clonal populations. After approximately 15 days, positive clones were selected and assayed for dual luciferase activity. A list of generated *T. brucei* recombinant cell lines is summarized in Supplementary Table 6.

For the analysis of RTi mutations in *T. brucei* endogenous tQ(CUG) variants, pLew100tQ(CUG)C28G42, pLew100tQ(CUG)G28C42, pLew100tQ(CUG)U28A42, and pLew100tQ(CUG)U28G42 (synthetized by Gene Universal) NotI digested (Supplementary Table 5) plasmids were used as above for dual luciferase activity measurement.

### SUPPLEMENTARY TABLES

**Supplementary Table 1. Yeast strains used in this study.**

| Strain | Genotype | Source of reference |
|---|---|---|
| H541 / L1751[a] | *MATa ade1-14 trp1-289 his3-Δ200 leu2-3,112 ura3-52* | [7] |
| H543 / L2521[a] | *MATa ade1-14 trp1-289 his3-Δ200 leu2-3,112 ura3-52 sup45-Y410S* | [8] |
| PBH140[a] | *MATa ade1-14 trp1-289 his3-Δ200 leu2-3,112 ura3-52 tif35Δ (YEp-TIF35-URA3)* | [9] |
| PBH156[a] | *MATa ade1-14 trp1-289 his3-Δ200 leu2-3,112 ura3-52 tif35Δ (YCp22-g/TIF35-TRP1)* | [10] |
| PBH159[a] | *MATa ade1-14 trp1-289 his3-Δ200 leu2-3,112 ura3-52 tif35Δ (YCp22-g/tif35-KLF-TRP1)* | This study |
| ySLDN1-1 / PBH291[b] | *MATa his3Δ1 leu2Δ0 ura3Δ0 met15Δ0* | [11] |
| ySLDN1-15 / PBH292[b] | *MATa his3Δ1 leu2Δ0 ura3Δ0 met15Δ0 ncs2Δ::HIS3MX* | [11] |
| ySLDN1-16 / PBH293[b] | *MATa his3Δ1 leu2Δ0 ura3Δ0 met15Δ0 ncs6Δ::HIS3MX* | [11] |
| YSL.KB04_DN1 / PBH294[b] | *MATa his3Δ1 leu2Δ0 ura3Δ0 met15Δ0 elp2Δ::HIS3MX* | [11] |
| YSL.KB04_DN3 / PBH295[b] | *MATa his3Δ1 leu2Δ0 ura3Δ0 met15Δ0 elp6Δ::HIS3MX* | [11] |
| ZH20 | *MATa ade1-14 trp1-289 his3-Δ200 leu2-3,112 ura3-52 rps25AΔ rps25BΔ* | This study |
| ZH31 [a] | *MATa ade1-14 trp1-289 his3-Δ200 leu2-3,112 ura3-52 tQ(CUG)MΔ [pRS426-tQ(C*UG)B-URA3]* | This study |
| ZH40 [a] | *MATa ade1-14 trp1-289 his3-Δ200 leu2-3,112 ura3-52 tQ(CUG)MΔ [YEplac181-tQ(CUG)M-LEU2]* | This study |
| ZH43 [a] | *MATa ade1-14 trp1-289 his3-Δ200 leu2-3,112 ura3-52 tQ(CUG)MΔ [YEplac181-tQ(CUG)M_A42G-LEU2]* | This study |
| ZH51 | *MATa ade1-14 trp1-289 his3-Δ200 leu2-3,112 ura3-52 rps15Δ (YEplac195-pRPS28B+FLAG-RPS15-URA3)* | This study |
| ZH54[a] | *MATa ade1-14 trp1-289 his3-Δ200 leu2-3,112 ura3-52 (YEplac112-TRP1)* | This study |

| | | |
|---|---|---|
| ZH56 | *MATa ade1-14 trp1-289 his3-Δ200 leu2-3,112 ura3-52 rps25AΔ rps25BΔ (YEplac112-TRP1)* | This study |
| ZH60 | *MATa ade1-14 his3 leu2 trp1 ura3 rps25AΔ_rps25BΔ (pRPS25A-FLAG -TRP1)* | This study |
| ZH62 | *MATa ade1-14 trp1-289 his3-Δ200 leu2-3,112 ura3-52 rps15Δ (YEplac112-pRPS28B+FLAG-RPS15-TRP1)* | This study |
| ZH65 | *MATa ade1-14 trp1-289 his3-Δ200 leu2-3,112 ura3-52 rps15Δ (YEplac112-pRPS28B+FLAG-RPS15_R130A-TRP1)* | This study |
| ZH68 | *MATa ade1-14 trp1-289 his3-Δ200 leu2-3,112 ura3-52 rps15Δ (YEplac112-pRPS28B+FLAG-RPS15_ R130A_S136A_R137A_K142A-TRP1)* | This study |
| ZH71 | *MATa ade1-14 trp1-289 his3-Δ200 leu2-3,112 ura3-52 rps15Δ (YEplac112-pRPS28B+FLAG-RPS15_G132S-TRP1)* | This study |
| ZH77 | *MATa ade1-14 trp1-289 his3-Δ200 leu2-3,112 ura3-52 rps15Δ (YEplac112-pRPS28B+FLAG-RPS15_128AAAAAAA134-TRP1)* | This study |
| ZH80 | *MATa ade1-14 trp1-289 his3-Δ200 leu2-3,112 ura3-52 rps25AΔ rps25BΔ (YEplac112-Δ2-13-FLAG-TRP1)* | This study |
| ZH112[a] | *MATa ade1-14 trp1-289 his3-Δ200 leu2-3,112 ura3-52 rps30AΔ::natNT2 rps30BΔ (YEplac195-RPS30B-URA3)* | This study |
| ZH117[a] | *MATa ade1-14 trp1-289 his3-Δ200 leu2-3,112 ura3-52 rps30AΔ::natNT2 rps30BΔ (YEplac112-RPS30B-FLAG-TRP1)* | This study |
| ZH121[a] | *MATa ade1-14 trp1-289 his3-Δ200 leu2-3,112 ura3-52 rps30AΔ::natNT2 rps30BΔ (YEplac112-RPS30B-Δ2-6-FLAG-TRP1)* | This study |
| ZH125[a] | *MATa ade1-14 trp1-289 his3-Δ200 leu2-3,112 ura3-52 rps30AΔ::natNT2 rps30BΔ (YEplac112-RPS30B-5AAAAAA10-FLAG-TRP1)* | This study |
| ZH129[a] | *MATa ade1-14 trp1-289 his3-Δ200 leu2-3,112 ura3-52 rps30AΔ::natNT2 rps30BΔ (YEplac112-RPS30B-A2G-FLAG-TRP1)* | This study |

| | | |
|---|---|---|
| ZH133[a] | *MATa ade1-14 trp1-289 his3-Δ200 leu2-3,112 ura3-52 rps30AΔ::natNT2 rps30BΔ (YEplac112-RPS30B-K3G-FLAG-TRP1)* | This study |
| ZH137[a] | *MATa ade1-14 trp1-289 his3-Δ200 leu2-3,112 ura3-52 rps30AΔ::natNT2 rps30BΔ (YEplac112-RPS30B-H5G-FLAG-TRP1)* | This study |
| ZH141[a] | *MATa ade1-14 trp1-289 his3-Δ200 leu2-3,112 ura3-52 rps30AΔ::natNT2 rps30BΔ (YEplac112-RPS30B-R10A-FLAG-TRP1)* | This study |
| ZH267[a] | *MATa ade1-14 trp1-289 his3-Δ200 leu2-3,112 ura3-52 rps30AΔ::natNT2 rps30BΔ (YEplac112-RPS30B-V4A-FLAG-TRP1)* | This study |
| ZH275[a] | *MATa ade1-14 trp1-289 his3-Δ200 leu2-3,112 ura3-52 rps30AΔ::natNT2 rps30BΔ (YEplac112-RPS30B-H5D-FLAG-TRP1)* | This study |
| ZH276[a] | *MATa ade1-14 trp1-289 his3-Δ200 leu2-3,112 ura3-52 rps30AΔ::natNT2 rps30BΔ (YEplac112-RPS30B-H5K-FLAG-TRP1)* | This study |
| ZH278[a] | *MATa ade1-14 trp1-289 his3-Δ200 leu2-3,112 ura3-52 rps30AΔ::natNT2 rps30BΔ (YEplac112-RPS30B-R10D-FLAG-TRP1)* | This study |

[a] indicates isogenic strain background

[b] indicates isogenic strain background

**Supplementary Table 2. Plasmids used in this study.**

| Plasmid | Description | Source of reference |
|---|---|---|
| PBB37 (YEp-R/T-CAAC-L) | high copy PGK-Renilla-Firefly R/T cassette (stop codon of Renilla is replaced with the CAA-C coding triplet for control readthrough measurements) in *LEU2* plasmid from YEplac181 | [12] |
| PBB78 (YEp-R/T-UAAC-L) | high copy PGK-Renilla-Firefly R/T cassette (stop codon of Renilla is UAA-C) in *LEU2* plasmid from YEplac181 | [9] |
| PBB79 (YEp-R/T-UAGC-L) | high copy PGK-Renilla-Firefly R/T cassette (stop codon of Renilla is UAG-C) in *LEU2* plasmid from YEplac181 | [9] |
| PBB158 (YEp-R/T-UAGU-L) | high copy PGK-Renilla-Firefly R/T cassette (stop codon of Renilla is UAG-U) in *LEU2* plasmid from YEplac181 | [10] |
| PBB159 (YEp-R/T-UAGA-L) | high copy PGK-Renilla-Firefly R/T cassette (stop codon of Renilla is UAG-A) in *LEU2* plasmid from YEplac181 | [10] |
| PBB160 (YEp-R/T-UAGG-L) | high copy PGK-Renilla-Firefly R/T cassette (stop codon of Renilla is UAG-G) in *LEU2* plasmid from YEplac181 | [10] |
| ZPB67 (YEp-R/T-UAG-BSC4-U) | high copy PGK-Renilla-Firefly R/T cassette (stop codon of Renilla is UAG-CAACTA) in *URA3* plasmid from YEplac195 | This study |
| ZPB68 (YEp-R/T-UAG-TMV-U) | high copy PGK-Renilla-Firefly R/T cassette (stop codon of Renilla is UAG-CAATTA) in *URA3* plasmid from YEplac195 | This study |
| ZPB70 (YEp-R/T-UAG-BSC4-L) | high copy PGK-Renilla-Firefly R/T cassette (stop codon of Renilla is UAG-CAACTA) in *LEU2* plasmid from YEplac181 | This study |
| ZPB71 (YEp-R/T-UAG-TMV-L) | high copy PGK-Renilla-Firefly R/T cassette (stop codon of Renilla is UAG-CAATTA) in *LEU2* plasmid from YEplac181 | This study |
| ZPB78 (YEp-R/T-UAA-BSC4-L) | high copy PGK-Renilla-Firefly R/T cassette (stop codon of Renilla is UAA-CAACTA) in *LEU2* plasmid from YEplac181 | This study |
| ZPB80 (YEp-R/T-UGA-BSC4-L) | high copy PGK-Renilla-Firefly R/T cassette (stop codon of Renilla is UGA-CAACTA) in *LEU2* plasmid from YEplac181 | This study |
| YEplac195 | high copy cloning vector, *URA3* | [13] |

| | | |
|---|---|---|
| PBB148 | high copy *S. cerevisiae* wild-type *tQ(UUG)B* in *URA3* plasmid from pRS426 | [10] |
| PBB150 | high copy *S. cerevisiae* wild-type *tQ(CUG)M* in *URA3* plasmid from pRS426 | [10] |
| PBB238 | high copy *S. cerevisiae tQ(U*UG)M_C34U* in *URA3* plasmid from pRS426 | This study |
| PBB239 | high copy *S. cerevisiae tQ(CUG)M_A42G_A51G_G66A [=tQ(C*UG)L_U34C]* in *URA3* plasmid from pRS426 | This study |
| PB240 | high copy *S. cerevisiae tQ(CUG)M_A42G* in *URA3* plasmid from pRS426 | This study |
| PBB241 | high copy *S. cerevisiae tQ(CUG)M_A51G* in *URA3* plasmid from pRS426 | This study |
| PBB242 | high copy *S. cerevisiae tQ(CUG)M_G66A* in *URA3* plasmid from pRS426 | This study |
| PBB243 | high copy *S. cerevisiae tQ(C*UG)B_U34C* in *URA3* plasmid from pRS426 | This study |
| ZPB4 | high copy *S. cerevisiae* wild-type *tQ(UUG)B* in *URA3* plasmid from pRS426 [tRNA in UTRs of *tQ(CUG)M]* | This study |
| ZPB5 | high copy *S. cerevisiae* wild-type *tQ(UUG)E2* in *URA3* plasmid from pRS426 [tRNA in UTRs of *tQ(CUG)M]* | This study |
| ZPB6 | high copy *S. cerevisiae* wild-type *tQ(UUG)L* in *URA3* plasmid from pRS426 [tRNA in UTRs of *tQ(CUG)M]* | This study |
| ZPB22 | high copy *S. cerevisiae tQ(UUG)L_G42A_A66G* in *URA3* plasmid from pRS426 | This study |
| ZPB23 | high copy *S. cerevisiae tQ(UUG)L_G51A* in *URA3* plasmid from pRS426 | This study |
| ZPB25 | high copy *S. cerevisiae tQ(UUG)L_G51A_A66G* in *URA3* plasmid from pRS426 | This study |
| ZPB26 | high copy *S. cerevisiae tQ(UUG)L _G42A_G51A* in *URA3* plasmid from pRS426 | This study |
| ZPB28 | high copy *S. cerevisiae tQ(CUG)M_ A42G_G66A* in *URA3* plasmid from pRS426 | This study |

| | | |
|---|---|---|
| ZPB36 | high copy *S. cerevisiae tQ(CUG)M_U28A_A42U* in *URA3* plasmid from pRS426 | This study |
| ZPB37 | high copy *S. cerevisiae tQ(CUG)M_U28C_A42G* in *URA3* plasmid from pRS426 | This study |
| ZPB42 | high copy *S. cerevisiae tQ(CUG)M_U28G_A42C* in *URA3* plasmid from pRS426 | This study |
| pRS316 | low copy cloning vector, *URA3* | [14] |
| ZPB43 (pRS316-*RPS25A*) | low copy *S. cerevisiae* wild-type *RPS25A* in *URA3* plasmid from pRS316 | This study |
| ZPB44 (YEplac195-pRPS28B+FLAG-*RPS25A*) | high copy *S. cerevisiae* wild-type *RPS25A* with FLAG tag at N-terminus under RPS28B promoter in *URA3* plasmid YEplac195 | This study |
| ZPB45 (YEplac112-pRPS28B+FLAG-*RPS25A*) | high copy *S. cerevisiae* wild-type *RPS25A* with FLAG tag at N-terminus under RPS28B promoter in *TRP1* plasmid YEplac112 | This study |
| pJD1055_RPS15 / ZPB50 | low copy *S. cerevisiae* wild-type *RPS15* in *HIS3* plasmid | [15] |
| ZPB51 (YEplac112-pRPS28B+FLAG-*RPS15*) | high copy *S. cerevisiae* wild-type *RPS15* with FLAG tag at N-terminus under RPS28B promoter in *TRP1* plasmid YEplac112 | This study |
| ZPB52 (YEplac112-pRPS28B+FLAG-*RPS15_R130A*) | high copy *S. cerevisiae* mutant *RPS15_R130A* with FLAG tag at N-terminus under RPS28B promoter in *TRP1* plasmid | This study |
| ZPB53 (YEplac195-pRPS28B+FLAG-*RPS15*) | high copy *S. cerevisiae* wild-type *RPS15* with FLAG tag at N-terminus under RPS28B promoter in *URA3* plasmid | This study |
| ZPB58 | high copy *S. cerevisiae* wild-type *tQ(CUG)M* in *LEU2* plasmid from YEplac181 | This study |
| ZPB59 | high copy *S. cerevisiae* wild-type *tQ(CUG)M_A42G* in *LEU2* plasmid from YEplac181 | This study |
| ZPB64 (YEplac112-pRPS28B+FLAG- | high copy *S. cerevisiae* mutant *RPS15_R130A_S136A_R137A_K142A* | This study |

| | | |
|---|---|---|
| *RPS15_R130A_S136 A_R137A_K142A*) | with FLAG tag at N-terminus under RPS28B promoter in *TRP1* plasmid | |
| ZPB65 (YEplac112-pRPS28B+FLAG-*RPS15_ 128AAAAAAA134*) | high copy *S. cerevisiae* mutant *RPS15_128AAAAAAA134* with FLAG tag at N-terminus under RPS28B promoter in *TRP1* plasmid | This study |
| ZPB66 (YEplac112-pRPS28B+FLAG-*RPS15 G132S*) | high copy *S. cerevisiae* mutant *RPS15_G132S* with FLAG tag at N-terminus under RPS28B promoter in *TRP1* plasmid | This study |
| ZPB74 (YEplac112-*RPS25A_Δ2-13*-FLAG*)* | high copy *S. cerevisiae* mutant *RPS25A_Δ2-13* with FLAG tag at C-terminus in *TRP1* plasmid YEplac112 | This study |
| ZPB82 | high copy *S. cerevisiae tQ(CUG)M_ A42C* in *URA3* plasmid from pRS426 | This study |
| ZPB83 | high copy *S. cerevisiae tQ(CUG)M_ U28G* in *URA3* plasmid from pRS426 | This study |
| ZPB84 | high copy *S. cerevisiae* tW(CCA)G1_ A42U in *URA3* plasmid from pRS426 | |
| PBB90 | high copy *S. cerevisiae* wild-type *tY(GUA)J2* in *URA3* plasmid from pRS426 | 9 |
| B670 YCp22-g/TIF35-KLF | single copy *TIF35-KLF-His* in*TRP1* plasmid from YCplac22 | 16 |
| pTH477 | high copy PGK-Renilla-Firefly R/T cassette (stop codon of Renilla is UGA-CCGUUC; for read-through measurements) in *URA3* plasmid from YEplac195 | 17 |
| YEplac112 | high copy cloning vector, *TRP1* | 13 |
| YEplac181 | high copy cloning vector, *LEU2* | 13 |
| pUG73 | loxP-flanked marker gene deletion cassette: *loxP-pKlLEU2-KlLEU2-tKlLEU2-loxP;* selectable phenotype: leucine prototr. | 3 |
| pSH47 | Cre-expressing (pGAL1-cre) CEN/ARS plasmid, marker gene: *pScURA3-ScURA3-tScURA3* | 3 |
| pZC4 | Lox2272-flanked marker gene deletion cassette: *lox2272-natNT2-lox2272;* selectable phenotype: nourseothricin resist. | 2 |

| | | |
|---|---|---|
| ZPB94 (YEplac195-*RPS30B*) | high copy *S. cerevisiae* wild-type *RPS30B* in *URA3* plasmid YEplac195 | This study |
| ZPB95 (YEplac112-*RPS30B*-FLAG) | high copy *S. cerevisiae* wild-type *RPS30B* with FLAG tag at C-terminus in *TRP1* plasmid YEplac112 | This study |
| ZPB96 (YEplac112-*RPS30B_Δ2-6*-FLAG) | high copy *S. cerevisiae* mutant *RPS30B_Δ2-6* with FLAG tag at C-terminus in *TRP1* plasmid YEplac112 | This study |
| ZPB97 (YEplac112-*RPS30B_5AAAAAA10*-FLAG) | high copy *S. cerevisiae* mutant *RPS30B_5AAAAAA10* with FLAG tag at C-terminus in *TRP1* plasmid YEplac112 | This study |
| ZPB98 (YEplac112-*RPS30B_A2G*-FLAG) | high copy *S. cerevisiae* mutant *RPS30B_A2G* with FLAG tag at C-terminus in *TRP1* plasmid YEplac112 | This study |
| ZPB99 (YEplac112-*RPS30B_K3G*-FLAG) | high copy *S. cerevisiae* mutant *RPS30B_K3G* with FLAG tag at C-terminus in *TRP1* plasmid YEplac112 | This study |
| ZPB100 (YEplac112-*RPS30B_H5G*-FLAG) | high copy *S. cerevisiae* mutant *RPS30B_H5G* with FLAG tag at C-terminus in *TRP1* plasmid YEplac112 | This study |
| ZPB101 (YEplac112-*RPS30B_R10A*-FLAG) | high copy *S. cerevisiae* mutant *RPS30B_R10A* with FLAG tag at C-terminus in *TRP1* plasmid YEplac112 | This study |
| ZPB106 | high copy *S. cerevisiae* tW(CCA)G1 in *URA3* plasmid from pRS426 | |
| ZPB145 (YEplac112-*RPS30B-V4A*-FLAG-TRP1) | high copy *S. cerevisiae* mutant *RPS30B-V4A* with FLAG tag at C-terminus in *TRP1* plasmid YEplac112 | This study |
| PMB4 (YEplac112-*RPS30B_H5D*-FLAG) | high copy *S. cerevisiae* mutant *RPS30B_H5D* with FLAG tag at C-terminus in *TRP1* plasmid YEplac112 | This study |
| PMB5 (YEplac112-*RPS30B_H5K*-FLAG) | high copy *S. cerevisiae* mutant *RPS30B_H5K* with FLAG tag at C-terminus in *TRP1* plasmid YEplac112 | This study |
| PMB6 (YEplac112-*RPS30B_R10D*-FLAG) | high copy *S. cerevisiae* mutant *RPS30B_R10D* with FLAG tag at C-terminus in *TRP1* plasmid YEplac112 | This study |
| ZPB147 | high copy *S. cerevisiae* *tC(GCA)P1_C27U_G41* in *URA3* plasmid from pRS426 | |

**Supplementary Table 3. Primers (Eurofins Genomics) used in this study.**

| Primer name | Primer sequence (5´to 3´) |
|---|---|
| 5z (tQM-LTTG-R) | TCCTACCCGGATTCGAACCGGGGTTGTCCGGATCAAAACCGAAAGTGATAACC |
| 6z (RPS25A + N5_F) | TAACAAGTATGTTTTTACTTTTTACTTTATCATAGAACATTTAATAAATCCTATAGAACGCGGCCGCCAG |
| 7z (RPS25A + C3_R) | AATATTATTGAAAATACAGATTATTTAAAATTATATAACCCGTTCCCTGTCACTATAGGGAGACCGGCAG |
| 8z (RPS25B + N5_F) | ACACTAACATCAATTTTCACCCTTTCTTACGTCTCAGAACATAATATAACCTATAGAACGCGGCCGCCAG |
| 9z (RPS25B + C3_R) | AAAAAAAGAAGAAGACAAACAACTTATTACCTTATTTCTTGATTTTATATCACTATAGGGAGACCGGCAG |
| 12z (RPS25A_F1) | AAATAAGGATCCAACGAAGGACGCCTTCTAC |
| 13z (RPS25A_R1) | AATAAGAATTCCCGTCACAATGCAACAGAAC |
| 14z (RPS25A_F2) | AATAAGGATCCATGCCTCCAAAGCAACAATTATCTAAAGC |
| 15z (RPS25A_R2) | AATAAGCATGCGCTACACTTACTTGCGTACTC |
| 18z (tQ(CUG)M + N5_F) | AGGTTCCATAAAACCGGAAGTTTTAGTGTACACTAACAACAGAAGAAAAACTATAGAACGCGGCCGCCAG |
| 19z (tQ(CUG)M + C3_R) | AAAAAAAAAAATGATGGTTTAAATTTCGTAAAATACGAAAAATGAAGGGACACTATAGGGAGACCGGCAG |
| 22z (RPS25_del2-13_R4) | ACCAGCAAGGGCAGCAGCCATGATTTATTAAATGTTCTATG |
| 23z (RPS25_del2-13_F5) | CATAGAACATTTAATAAATCATGGCTGCTGCCCTTGCTGGTG |
| 29z (F7_RPS25A_FLAG) | GATTACAAGGATGACGACGATAAGTAAACAGGGAACGGGTTATA |
| 30z (R6_RPS25A_FLAG) | TTACTTATCGTCGTCATCCTTGTAATCTTCAGAAGCAGTAGCTC |
| 31z (RPS15 + N5_F) | TGTAGAATAAGACGAACTAGAAGTACACAACAAGATAATCACGACCGATCCTATAGAACGCGGCCGCCAG |
| 32z (RPS15 + C3_R) | TTATATATAAAAAAGATACAAGGATGACTGCGATTCTGTTATTAGGGAGCCACTATAGGGAGACCGGCAG |
| 35z (RPS15_F2) | ACTACTGCTGCTTTCATCCCATTGGCTTAAGCTCCCTAATAACAGAATC |
| 36z (RPS15_R2) | GATGAAAGCAGCAGTAGTAGCACCGGCAGCACCATGTCTGACTGGGGTG |
| 37z (RPS15_F3) | GCTGCTGCCGCTGCTGCTACTTCCCGTTTCATCCCATTG |

| | |
|---|---|
| 38z (RPS15_R3) | GCAGCAGCGGCAGCAGCAGCTCTGACTGGGGTGTAAG TAATGGAG |
| 39z (RPS15_F4) | GTAGAGCCAGCGCTACTACTTCCCGTTTC |
| 40z (RPS15_R4) | GTAGTAGCGCTGGCTCTACCATGTCTGAC |
| 42z (BSC4_UAG_F) | CAAATGTCGACGTGCGATTAGCAACTAGGATCCTTCAAC TTCCCTG |
| 43z (TMV_UAG_F) | CAAATGTCGACGTGCGATTAGCAATTAGGATCCTTCAAC TTCCCTG |
| 44z (BSC4_UAA_F) | CAAATGTCGACGTGCGATTAACAACTAGGATCCTTCAAC TTCCCTG |
| 46z *(RPS30A + N5_F)* | ATTTGCCACTGTAATAATCTTCCATATCCCCATACAAAAA CTACGCAAATCTATAGAACGCGGCCGCCAG |
| 47z (RPS30A + C3_R) | ATTTTGTTTCTTGTTACTTTTAAGCTCCTTTTCCACAACTG TTAATTTTCCACTATAGGGAGACCGGCAG |
| 48z (RPS30B + N5_F) | TCTGAATTAGCACTCCTTCCTCAAGATACTAGATAAACAA ATTATACAAACTATAGAACGCGGCCGCCAG |
| 49z (RPS30B + C3_R) | TGTTTAATTACACATGTAGAACAATAAAAGTATAGAATTT TAGATAGTATCACTATAGGGAGACCGGCAG |
| 52z (RPS30B_F1) | AAATAAGGATCCCGAAATGTTGTGGTTCCTCTC |
| 53z (RPS30B_R1) | AATAAGAATTCCTTCAATTAGAGGGGCAGGTGG |
| 54z (RPS30B-FLAG tag_F2) | GATTACAAGGATGACGACGATAAGTAGATACTATCTAAA ATTC |
| 55z (RPS30B-FLAG tag_R2) | CTTATCGTCGTCATCCTTGTAATCTTGGACAGATGGACC TGGG |
| 62z (RPS30B_del2-6_F5) | TCTCCTCAGTCTCTAGCTCGTGCTGGT |
| 63z (RPS30B_del2-6_R5) | AGCTAGAGACTGAGGAGAATTTGAAAA |
| 64z (RPS30B_5AAAAAA 10_F6) | GCCGCTGCTGCTGCTGCTGCTGGTAAAGTTAAGTCTC |
| 65z (RPS30B_5AAAAAA 10_R6) | AGCAGCAGCAGCAGCGGCAACTTTAGCCTGAGGAGA |
| 66z (RPS30B_A2G_F7) | GGTAAAGTTCACGGTTCTCTAG |
| 67z | AGAACCGTGAACTTTACCCTGAGGAGAATTTGAAA |

| | |
|---|---|
| (RPS30B_A2G_R7) | |
| 68z (RPS30B_K3G_F8) | GCTGGAGTTCACGGTTCTCTAGC |
| 69z (RPS30B_K3G_R8) | AGAACCGTGAACTCCAGCCTGAGGAGAATTTGAAAA |
| 70z (RPS30B_H5G_F9) | CTAAAGTTGGCGGTTCTCTAGCTCGTGC |
| 71z (RPS30B_H5G_R9) | GAGAACCGCCAACTTTAGCCTGAGGAGA |
| 9p (RPS30B_H5D_F) | CTAAAGTGTCCGGTTCTCTAGCTCGTGC |
| 10p (RPS30B_H5D_R) | GAGAACCCAGAACTTTAGCCTGAGGAGA |
| 7p (RPS30B_H5K_F) | CTAAAGTTTTCGGTTCTCTAGCTCGTGC |
| 8p (RPS30B_H5K_R) | GAGAACCAAAAACTTTAGCCTGAGGAGA |
| 72z RPS30B_R10A_F10 | CTCTAGCTGCTGCTGGTAAAGTTAAGTC |
| 73z RPS30B_R10A_R10 | TACCAGCAGCAGCTAGAGAACCGTGAAC |
| 3p (RPS30B_R10D_F) | CTCTAGCGTCTGCTGGTAAAGTTAAGTC |
| 4p (RPS30B_R10D_R) | TACCAGCCAGAGCTAGAGAACCGTGAAC |
| 111z (eS30B_V4A_F11) | GGCTAAAGCTCACGGTTCTCTAGCTCG |
| 112z (eS30B_V4A_R11) | GAACCGTGAGCTTTAGCCTGAGGAGAAT |
| PB43 (PBRFNotI) | CTCGAAGCGGCCGCTCTAGAATTACAC |
| PB78 (BSC4) | CAAATGTCGACGTGCGATTGACAACTAGGATCCTTCAAC TTCCCTGAGCTCG |
| PB134 (PBtQBXhoI) | AATAACTCGAGTTTTCAAACCACTCAATTTAAAAAATTGT CAG |
| PB135 (PBtQBBamHI-R) | AATAAGGATCCATCTCCATTCTAAGAGTGTCCGATAATT CATG |

| | |
|---|---|
| PB140 (PBtQMXhoI) | AATAACTCGAGAGGTTCCATAAAACCGGAAGTTTTAGTG |
| PB141 (PBtQMBamHI-R) | AATAAGGATCCAAAAAAAAAAATGATGGTTTAAATTTCGTAAAATACG |
| PB188 (tQMTTG-F) | GTGGTTATCACTTTCGGTTTTGATCCGAACAACCCC |
| PB189 (tQMTTG-R) | GGGGTTGTTCGGATCAAAACCGAAAGTGATAACCAC |
| PB190 (tQM-LCTG-F) | CCGGACAACCCCGGTTCGAATCCGGGTAGGACCTTCCCTTCATTTTTCG |
| PB191 (tQM-LCTG-R) | TCCTACCCGGATTCGAACCGGGGTTGTCCGGATCAGAACCGAAAGTGATAACC |
| PB192 (tQM42G-F) | CACTTTCGGTTCTGATCCGGACAACCCCAGTTCGAATCCG |
| PB193 (tQM42G-R) | CGGATTCGAACTGGGGTTGTCCGGATCAGAACCGAAAGTG |
| PB194 (tQM51G-F) | GTTCTGATCCGAACAACCCCGGTTCGAATCCGGGTGGG |
| PB195 (tQM51G-R) | CCCACCCGGATTCGAACCGGGGTTGTTCGGATCAGAAC |
| PB196 (tQM66A-F) | CCCAGTTCGAATCCGGGTAGGACCTTCCCTTCATTTTTCG |
| PB197 (tQM66A-R) | CGAAAAATGAAGGGAAGGTCCTACCCGGATTCGAACTGGG |
| PB198 (tQBCTG-F) | GTGGTTATCACTTTCGGTTCTGATCCGGACAACCCC |
| PB199 (tQBCTG-R) | GGGGTTGTCCGGATCAGAACCGAAAGTGATAACCAC |
| PB248 (PBuS19_r) | GCTTGCATGCCTGCAGGAGTCCGTGACAGTAAATG |
| PB249 (PBuS19R130A_f) | CCCAGTCAGACATGGTGCCGCCGGTGCTACTACTT |
| PB250 (PBuS19R130A_r) | AAGTAGTAGCACCGGCGGCACCATGTCTGACTGGG |
| KQ (*S.c.* tQ probe) | AGGTCCCACCCGGATTCGAACTGGGGTTGT - $^{32}P$ labelled |
| KQ (*S.c.* tQ probe) | DIG labelled - AGGTCCCACCCGGATTCGAACTGGGGTTGT |
| 79z (*S.c.* 5.8S rRNA) | GCTGCGTTCTTCATCGATGCGAGAACCAA - $^{32}P$ labelled |
| 79z (*S.c.* 5.8S rRNA) | DIG labelled - GCTGCGTTCTTCATCGATGCGAGAACCAA |
| Tb 5.8S | ATTGGGCAATGAAATGATTCTG - $^{32}P$ labelled |

| | |
|---|---|
| (*T.b.* 5.8S rRNA) | |
| ZP710R-TbGlnCUG | CTGGACTCGAACCAGGGTTATCG - $^{32}$P labelled |

**Supplementary Table 4. GeneArt Strings DNA Fragments (Invitrogen) used in this study.**

| | |
|---|---|
| tQ(UUG)B_tQ-M UTRs | TTGGGTACCGGGCCCCCCCTCGAGAGGTTCCATA<br>AAACCGGAAGTTTTAGTGTACACTAACAACAGAAG<br>AAAAAGGTTTTATAGTGTAGTGGTTATCACTTTCGG<br>TTTTGATCCGGACAACCCCGGTTCGAATCCGGGTA<br>AGACCTTCCCTTCATTTTTCGTATTTTACGAAATTT<br>AAACCATCATTTTTTTTTTTGGATCCACTAGTTCTA<br>GAGCGGCCGCCA |
| tQ(UUG)E2_tQ-M UTRs | TTGGGTACCGGGCCCCCCCTCGAGAGGTTCCATA<br>AAACCGGAAGTTTTAGTGTACACTAACAACAGAAG<br>AAAAAGGTTTTATAGTGTAGTGGTTATCACTTTCGG<br>TTTTGATCCGAACAACCCCGGTTCGAATCCGGGTA<br>AGACCTTCCCTTCATTTTTCGTATTTTACGAAATTT<br>AAACCATCATTTTTTTTTTTGGATCCACTAGTTCTA<br>GAGCGGCCGCCA |
| tQ(UUG)M_A51G | TTGGGTACCGGGCCCCCCCTCGAGAGGTTCCATA<br>AAACCGGAAGTTTTAGTGTACACTAACAACAGAAG<br>AAAAAGGTCCTATAGTGTAGTGGTTATCACTTTCG<br>GTTTTGATCCGAACAACCCCGGTTCGAATCCGGGT<br>GGGACCTTCCCTTCATTTTTCGTATTTTACGAAATT<br>TAAACCATCATTTTTTTTTTTGGATCCACTAGTTCTA<br>GAGCGGCCGCCA |
| tQ(UUG)M_A42G_G66A | TTGGGTACCGGGCCCCCCCTCGAGAGGTTCCATA<br>AAACCGGAAGTTTTAGTGTACACTAACAACAGAAG<br>AAAAAGGTCCTATAGTGTAGTGGTTATCACTTTCG<br>GTTTTGATCCGGACAACCCCAGTTCGAATCCGGGT<br>AGGACCTTCCCTTCATTTTTCGTATTTTACGAAATT<br>TAAACCATCATTTTTTTTTTTGGATCCACTAGTTCTA<br>GAGCGGCCGCCA |
| tQ(UUG)M_A42G | TTGGGTACCGGGCCCCCCCTCGAGAGGTTCCATA<br>AAACCGGAAGTTTTAGTGTACACTAACAACAGAAG<br>AAAAAGGTCCTATAGTGTAGTGGTTATCACTTTCG<br>GTTTTGATCCGGACAACCCCAGTTCGAATCCGGGT<br>GGGACCTTCCCTTCATTTTTCGTATTTTACGAAATT<br>TAAACCATCATTTTTTTTTTTGGATCCACTAGTTCTA<br>GAGCGGCCGCCA |
| tQ(UUG)M_G66A | TTGGGTACCGGGCCCCCCCTCGAGAGGTTCCATA<br>AAACCGGAAGTTTTAGTGTACACTAACAACAGAAG<br>AAAAAGGTCCTATAGTGTAGTGGTTATCACTTTCG<br>GTTTTGATCCGAACAACCCCAGTTCGAATCCGGGT<br>AGGACCTTCCCTTCATTTTTCGTATTTTACGAAATT |

| | |
|---|---|
| | TAAACCATCATTTTTTTTTTTGGATCCACTAGTTCTA<br>GAGCGGCCGCCA |
| tQ(CUG)M_A42G_G66A | TTGGGTACCGGGCCCCCCCTCGAGAGGTTCCATA<br>AAACCGGAAGTTTTAGTGTACACTAACAACAGAAG<br>AAAAAGGTCCTATAGTGTAGTGGTTATCACTTTCG<br>GTTCTGATCCGGACAACCCCAGTTCGAATCCGGG<br>TAGGACCTTCCCTTCATTTTTCGTATTTTACGAAAT<br>TTAAACCATCATTTTTTTTTTTGGATCCACTAGTTCT<br>AGAGCGGCCGCCA |
| tQ(CUG)M_U28A_A42U | TTGGGTACCGGGCCCCCCCTCGAGAGGTTCCATA<br>AAACCGGAAGTTTTAGTGTACACTAACAACAGAAG<br>AAAAAGGTCCTATAGTGTAGTGGTTATCACTTACG<br>GTTCTGATCCGTACAACCCCAGTTCGAATCCGGGT<br>GGGACCTTCCCTTCATTTTTCGTATTTTACGAAATT<br>TAAACCATCATTTTTTTTTTTGGATCCACTAGTTCTA<br>GAGCGGCCGCCA |
| tQ(CUG)M_U28C_A42G | TTGGGTACCGGGCCCCCCCTCGAGAGGTTCCATA<br>AAACCGGAAGTTTTAGTGTACACTAACAACAGAAG<br>AAAAAGGTCCTATAGTGTAGTGGTTATCACTTCCG<br>GTTCTGATCCGGACAACCCCAGTTCGAATCCGGG<br>TGGGACCTTCCCTTCATTTTTCGTATTTTACGAAAT<br>TTAAACCATCATTTTTTTTTTTGGATCCACTAGTTCT<br>AGAGCGGCCGCCA |
| tQ(CUG)M_U28G_A42C | TTGGGTACCGGGCCCCCCCTCGAGAGGTTCCATA<br>AAACCGGAAGTTTTAGTGTACACTAACAACAGAAG<br>AAAAAGGTCCTATAGTGTAGTGGTTATCACTTGCG<br>GTTCTGATCCGCACAACCCCAGTTCGAATCCGGGT<br>GGGACCTTCCCTTCATTTTTCGTATTTTACGAAATT<br>TAAACCATCATTTTTTTTTTTGGATCCACTAGTTCTA<br>GAGCGGCCGCCA |
| tQ(CUG)M_A42C | TTGGGTACCGGGCCCCCCCTCGAGAGGTTCCATA<br>AAACCGGAAGTTTTAGTGTACACTAACAACAGAAG<br>AAAAAGGTCCTATAGTGTAGTGGTTATCACTTTCG<br>GTTCTGATCCGCACAACCCCAGTTCGAATCCGGGT<br>GGGACCTTCCCTTCATTTTTCGTATTTTACGAAATT<br>TAAACCATCATTTTTTTTTTTGGATCCACTAGTTCTA<br>GAGCGGCCGCCA |
| tQ(UUG)M_U28G | TTGGGTACCGGGCCCCCCCTCGAGAGGTTCCATA<br>AAACCGGAAGTTTTAGTGTACACTAACAACAGAAG<br>AAAAAGGTCCTATAGTGTAGTGGTTATCACTTGCG<br>GTTCTGATCCGAACAACCCCAGTTCGAATCCGGGT<br>GGGACCTTCCCTTCATTTTTCGTATTTTACGAAATT |

| | |
|---|---|
| | TAAACCATCATTTTTTTTTTTGGATCCACTAGTTCTAGAGCGGCCGCCA |
| tC(GCA)P1_C27U_G41 | TTGGGTACCGGGCCCCCCCTCGAGTTGCGTGGATAAGTGTTATTATTCTATTGCCTTAAAACTATACAACAAAAGCTCGTATGGCGCAGTGGTAGCGCAGTAGATTGCAAATCTGTTGGTCCTTAGTTCGATCCTGAGTGCGAGCTTTCTTTTTTTCCAAAGAATTTTAATTATATACGTTCGCCTGTACGGCTTTAGGATCCACTAGTTCTAGAGCGGCCGCCA |

**Supplementary Table 5. Gene Universal DNA Fragments used for the generation of *T. brucei* strains.**

| | |
|---|---|
| tQ(CUG)U28A42 | CGGGGTACCTCCTATTTTCGACCCCTAACCCGCCT GCCGGCACGCCCCAAATGGCGCGGCCCACATTTG ACGTAGAAACCATTTGCATATATGACTCTCCCGAG GACCCGCGCCCCGGGCAGACTCGAAGGGGGGAC GTTCTGCGCATAACGCTAAGTGGCGGCGAACAGC AAGCCTCCCGTAACGTCCCTGTAAACGCGGCCGA CCCGATCCAATAGTAAAACACAAGGGCCACGTCCT CTTCCCGCCCCCGCCGCAGCAGTAAGGTTATGCG TGAGAGGTTTTGACTGCCACTTGGAACGCTGAACC GTGGCGTAAGAGTCTCCGCAAAGGAAAACGAATG CTCCCGGCGGGGTTCGAACCCGCGATATTCGGTT CATAAGACCAACGTCCTAACCAACTAGACCACGGG AGCATTGGCGCCGGCCACTTCGACAGATACAGAG GAAAGGACCGCCGAGTAAACTTCTTCGTTTGTATT AGAAAAAAACAGCCTAACGGCAGTGGCCGGGTCC TATAGTGTAGTGGTTATCACTTTCGGTTCTGATCC GAACAACCCCAGTTCGAATCCGGGTGGGACCTAA ATTTTCGTTTTTTTCAATTCACTAACTATGGTGTCTT TTCACCATCTGCGGCAGTGCGTGGGTGATCTAGT GGTTATGATGTCTGCTTTACACGCAGAACGTCGCG GGTTCGAACCCCGCCCCGCGTAGCGTTTTCCAGT TTGCGGCTGCGTCTCTTCATGTATGGTTCCGATTA ATGAAATTGTTGAGGAACACGTAGCCCTTCTAGCT CAGTCGGTAGAGCGCACGGCTCTTAACCGTGTGG TCGTGGGTTCGAGCCCCACGGGGGGTGCTTTTCT CCGATTTCCCTGATTTAACTTCCACAATGGGGTGC CTCGAGTGCTGTCCCGTAAGGCGGTGTGATGTGT CAACAGGCCTTCCTTCAGCAGCGTCACAGTGATAT CGTCGTCACTAAAGGGGAAGTGCCTTCTTCAGGTT GTTGATCTAACATGTTATTGGCCGAAGGCTCCTGC GGTGGACGGCGCATCCCACCACCACCTCCCCCCC CCCCCCCGAGAGAGGATCCGCG |
| *Q(CUG)U28GXA42C | CGGGGTACCTCCTATTTTCGACCCCTAACCCGCCT GCCGGCACGCCCCAAATGGCGCGGCCCACATTTG ACGTAGAAACCATTTGCATATATGACTCTCCCGAG GACCCGCGCCCCGGGCAGACTCGAAGGGGGGAC GTTCTGCGCATAACGCTAAGTGGCGGCGAACAGC AAGCCTCCCGTAACGTCCCTGTAAACGCGGCCGA CCCGATCCAATAGTAAAACACAAGGGCCACGTCCT CTTCCCGCCCCCGCCGCAGCAGTAAGGTTATGCG TGAGAGGTTTTGACTGCCACTTGGAACGCTGAACC GTGGCGTAAGAGTCTCCGCAAAGGAAAACGAATG |

| | |
|---|---|
| | CTCCCGGCGGGGTTCGAACCCGCGATATTCGGTT CATAAGACCAACGTCCTAACCAACTAGACCACGGG AGCATTGGCGCCGGCCACTTCGACAGATACAGAG GAAAGGACCGCCGAGTAAACTTCTTCGTTTGTATT AGAAAAAAACAGCCTAACGGCAGTGGCCGGGTCC TATAGTGTAGTGGTTATCACTTGCGGTTCTGATCC GCACAACCCCAGTTCGAATCCGGGTGGGACCTAA ATTTTCGTTTTTTTCAATTCACTAACTATGGTGTCTT TTCACCATCTGCGGCAGTGCGTGGGTGATCTAGT GGTTATGATGTCTGCTTTACACGCAGAACGTCGCG GGTTCGAACCCCGCCCCGCGTAGCGTTTTCCAGT TTGCGGCTGCGTCTCTTCATGTATGGTTCCGATTA ATGAAATTGTTGAGGAACACGTAGCCCTTCTAGCT CAGTCGGTAGAGCGCACGGCTCTTAACCGTGTGG TCGTGGGTTCGAGCCCCACGGGGGGTGCTTTTCT CCGATTTCCCTGATTTAACTTCCACAATGGGGTGC CTCGAGTGCTGTCCCGTAAGGCGGTGTGATGTGT CAACAGGCCTTCCTTCAGCAGCGTCACAGTGATAT CGTCGTCACTAAAGGGGAAGTGCCTTCTTCAGGTT GTTGATCTAACATGTTATTGGCCGAAGGCTCCTGC GGTGGACGGCGCATCCCACCACCACCTCCCCCCC CCCCCCCGAGAGAGGATCCGCG |
| tQ(CUG)C28G42 | GGTACCAAGCTTTGTCGTCACGACCCTTTACTGTG TAATATACGGTGCTGCTTTGGCTGGGCCGCACCG TTGCTCCTATAGCTCAGTCGGTTAGAGCGTGGGTC TAATAAGCCCAAGGTCACAGGTTCGACCCCTGTTG GGAGCACGTTTTTCAGTGCGGGAGACTGTGCTTG TTGAAGCTAAAGGCAAAAAACTGCTGAGGGTGGG GTTCGAACCCACGAAGCAACCGCATGAGAACTTAA GTCTCACCCCTTTGACCAACTCGGGAACCCCAGC GGCACGTTCGCCCGGCTAATCAAACTAGGGCCAA CATTCTCCACACAAACTAATTTCCAACAGATTTGAG CACTGTGGCGTTATGTGTCAAAAAGCACTCCCACC TGGACTCGAACCAGGGTTATCGGATTCAGAGTCC GAGGTGATAACCGCTACACTATGGGAGCGACGGT CCCAACGACTACTTGAAGCAACTATCAAAACCGAA TTAAAGATATACAGAAAGGAACATTTTGTACATTTT CGCTGCGCCCGCGCTCCCATAGTGTAGCGGTTAT CACCTCGGACTCTGAATCCGATAACCCTGGTTCGA GTCCAGGTGGGAGTGCTTTTTGACACATAACGCCA CAGTGCTCAAATCTGTTGGAAATTAGTTTGTGTGAT TGGGTATTGCAAGAGACCCCGCAGCTCCTATAGCT CAGTCGGTTAGAGCGTGGGTCTAATAAGCCCAAG GTCACAGGTTCGACCCCTGTTGGGAGCACTTTTTC |

| | |
|---|---|
| | TCTACCTATCAGTTCGTTGTTTTCACGTGTAGCTCT GAACGTTCGCGTGCTGGGCTCGTGTTCTTGGAAG GAAACTGACTGGCTCAGCATTCCCGTCGTTGCGA CCAATTTTTCATAGTTTCATAAAAGAAGCAAGCTGA AACATTTTACTTAGTGAAGCTGTTTCCTGCAACGTC CAGAAAACGAATATCTAAAGGAGGGCGGCTGTTTA CCGCGCTAGGGGCCTTTTGGATCC |
| tQ(CUG)G28C42 | GGTACCAAGCTTTGTCGTCACGACCCTTTACTGTG TAATATACGGTGCTGCTTTGGCTGGGCCGCACCG TTGCTCCTATAGCTCAGTCGGTTAGAGCGTGGGTC TAATAAGCCCAAGGTCACAGGTTCGACCCCTGTTG GGAGCACGTTTTTCAGTGCGGGAGACTGTGCTTG TTGAAGCTAAAGGCAAAAAACTGCTGAGGGTGGG GTTCGAACCCACGAAGCAACCGCATGAGAACTTAA GTCTCACCCCTTTGACCAACTCGGGAACCCCAGC GGCACGTTCGCCCGGCTAATCAAACTAGGGCCAA CATTCTCCACACAAACTAATTTCCAACAGATTTGAG CACTGTGGCGTTATGTGTCAAAAAGCACTCCCACC TGGACTCGAACCAGGGTTATCGGATTCAGAGTCC GAGGTGATAACCGCTACACTATGGGAGCGACGGT CCCAACGACTACTTGAAGCAACTATCAAAACCGAA TTAAAGATATACAGAAAGGAACATTTTGTACATTTT CGCTGCGCCCGCGCTCCCATAGTGTAGCGGTTAT CACCTGGGACTCTGAATCCCATAACCCTGGTTCGA GTCCAGGTGGGAGTGCTTTTTGACACATAACGCCA CAGTGCTCAAATCTGTTGGAAATTAGTTTGTGTGAT TGGGTATTGCAAGAGACCCCGCAGCTCCTATAGCT CAGTCGGTTAGAGCGTGGGTCTAATAAGCCCAAG GTCACAGGTTCGACCCCTGTTGGGAGCACTTTTTC TCTACCTATCAGTTCGTTGTTTTCACGTGTAGCTCT GAACGTTCGCGTGCTGGGCTCGTGTTCTTGGAAG GAAACTGACTGGCTCAGCATTCCCGTCGTTGCGA CCAATTTTTCATAGTTTCATAAAAGAAGCAAGCTGA AACATTTTACTTAGTGAAGCTGTTTCCTGCAACGTC CAGAAAACGAATATCTAAAGGAGGGCGGCTGTTTA CCGCGCTAGGGGCCTTTTGGATCC |
| tQ(CUG)U28A42 | GGTACCAAGCTTTGTCGTCACGACCCTTTACTGTG TAATATACGGTGCTGCTTTGGCTGGGCCGCACCG TTGCTCCTATAGCTCAGTCGGTTAGAGCGTGGGTC TAATAAGCCCAAGGTCACAGGTTCGACCCCTGTTG GGAGCACGTTTTTCAGTGCGGGAGACTGTGCTTG TTGAAGCTAAAGGCAAAAAACTGCTGAGGGTGGG GTTCGAACCCACGAAGCAACCGCATGAGAACTTAA |

| | |
|---|---|
| | GTCTCACCCCTTTGACCAACTCGGGAACCCCAGC GGCACGTTCGCCCGGCTAATCAAACTAGGGCCAA CATTCTCCACACAAACTAATTTCCAACAGATTTGAG CACTGTGGCGTTATGTGTCAAAAAGCACTCCCACC TGGACTCGAACCAGGGTTATCGGATTCAGAGTCC GAGGTGATAACCGCTACACTATGGGAGCGACGGT CCCAACGACTACTTGAAGCAACTATCAAAACCGAA TTAAAGATATACAGAAAGGAACATTTTGTACATTTT CGCTGCGCCCGCGCTCCCATAGTGTAGCGGTTAT CACCTTGGACTCTGAATCCAATAACCCTGGTTCGA GTCCAGGTGGGAGTGCTTTTTGACACATAACGCCA CAGTGCTCAAATCTGTTGGAAATTAGTTTGTGTGAT TGGGTATTGCAAGAGACCCCGCAGCTCCTATAGCT CAGTCGGTTAGAGCGTGGGTCTAATAAGCCCAAG GTCACAGGTTCGACCCCTGTTGGGAGCACTTTTTC TCTACCTATCAGTTCGTTGTTTTCACGTGTAGCTCT GAACGTTCGCGTGCTGGGCTCGTGTTCTTGGAAG GAAACTGACTGGCTCAGCATTCCCGTCGTTGCGA CCAATTTTTCATAGTTTCATAAAAGAAGCAAGCTGA AACATTTTACTTAGTGAAGCTGTTTCCTGCAACGTC CAGAAAACGAATATCTAAAGGAGGGCGGCTGTTTA CCGCGCTAGGGGCCTTTTGGATCC |
| tQ(CUG)U28G42 | GGTACCAAGCTTTGTCGTCACGACCCTTTACTGTG TAATATACGGTGCTGCTTTGGCTGGGCCGCACCG TTGCTCCTATAGCTCAGTCGGTTAGAGCGTGGGTC TAATAAGCCCAAGGTCACAGGTTCGACCCCTGTTG GGAGCACGTTTTTCAGTGCGGGAGACTGTGCTTG TTGAAGCTAAAGGCAAAAAACTGCTGAGGGTGGG GTTCGAACCCACGAAGCAACCGCATGAGAACTTAA GTCTCACCCCTTTGACCAACTCGGGAACCCCAGC GGCACGTTCGCCCGGCTAATCAAACTAGGGCCAA CATTCTCCACACAAACTAATTTCCAACAGATTTGAG CACTGTGGCGTTATGTGTCAAAAAGCACTCCCACC TGGACTCGAACCAGGGTTATCGGATTCAGAGTCC GAGGTGATAACCGCTACACTATGGGAGCGACGGT CCCAACGACTACTTGAAGCAACTATCAAAACCGAA TTAAAGATATACAGAAAGGAACATTTTGTACATTTT CGCTGCGCCCGCGCTCCCATAGTGTAGCGGTTAT CACCTTGGACTCTGAATCCGATAACCCTGGTTCGA GTCCAGGTGGGAGTGCTTTTTGACACATAACGCCA CAGTGCTCAAATCTGTTGGAAATTAGTTTGTGTGAT TGGGTATTGCAAGAGACCCCGCAGCTCCTATAGCT CAGTCGGTTAGAGCGTGGGTCTAATAAGCCCAAG GTCACAGGTTCGACCCCTGTTGGGAGCACTTTTTC |

| | |
|---|---|
| | TCTACCTATCAGTTCGTTGTTTTCACGTGTAGCTCT GAACGTTCGCGTGCTGGGCTCGTGTTCTTGGAAG GAAACTGACTGGCTCAGCATTCCCGTCGTTGCGA CCAATTTTTCATAGTTTCATAAAAGAAGCAAGCTGA AACATTTTACTTAGTGAAGCTGTTTCCTGCAACGTC CAGAAAACGAATATCTAAAGGAGGGCGGCTGTTTA CCGCGCTAGGGGCCTTTTGGATCC |

**Supplementary Table 6. *T. brucei* strains used in this study.**

| Strain | Description | Source of reference |
|---|---|---|
| TbUGA-C | *T7 Pol/Tet inducible expression of UAG-C dual luciferase system* | [6] |
| TbDLC-C | *T7 Pol/Tet inducible expression of DLC-C dual luciferase system* | [6] |
| TbtQ(CUG)U28A42:UAG-C | *T7 Pol/Tet inducible expression of UAG-C dual luciferase system*[6]<br>*Constitutive expression of S.c. tQ(CUG)U28A42 (pLew100, Phleo)* | This study |
| TbtQ(CUG)U28A42:DLC-C | *T7 Pol/Tet inducible expression of DLC-C dual luciferase system*[6]<br>*Constitutive expression of S.c. tQ(CUG)U28A42 (pLew100, Phleo)* | This study |
| Tb*Q(CUG)U28GXA42C:UAG-C | *T7 Pol/Tet inducible expression of UAG-C dual luciferase system*[6]<br>*Constitutive expression of S.c. *Q(CUG)U28GXA42C (pLew100, Phleo)* | This study |
| Tb*Q(CUG)U28GXA42C:DLC-C | *T7 Pol/Tet inducible expression of DLC-C dual luciferase system*[6]<br>*Constitutive expression of S.c. *Q(CUG)U28GXA42C (pLew100, Phleo)* | This study |
| TbQ(CUG)C28G42:DLC-C | *T7 Pol/Tet inducible expression of DLC-C dual luciferase system*[6]<br>*Constitutive expression of Tb* Q(CUG)C28G42 *(pLew100, Phleo)* | This study |
| TbQ*(CUG)C28GXG42C:DLC-C | *T7 Pol/Tet inducible expression of DLC-C dual luciferase system*[6] | This study |

| | | |
|---|---|---|
| | *Constitutive expression of Tb* Q(CUG)C28GXG42C *(pLew100, Phleo)* | |
| TbQ*(CUG)C28UXG42A:DLC-C | *T7 Pol/Tet inducible expression of DLC-C dual luciferase system*[6] *Constitutive expression of Tb* Q(CUG) C28UXG42A *(pLew100, Phleo)* | This study |
| TbQ*(CUG)C28UXG42:DLC-C | *T7 Pol/Tet inducible expression of DLC-C dual luciferase system*[6] *Constitutive expression of Tb* Q(CUG)C28UXG42 *(pLew100, Phleo)* | This study |
